# Reconfiguration of brain-wide neural activity after early life adversity

**DOI:** 10.1101/2023.09.10.557058

**Authors:** Taylor W. Uselman, Russell E. Jacobs, Elaine L Bearer

## Abstract

Early life adversity (ELA) predisposes individuals to physical and mental disorders lifelong. How ELA affects brain functions, leading to these vulnerabilities, is a mystery. To understand ELA’s impacts, investigations into neural activity affected by ELA must go beyond localized areas toward simultaneous recordings from multiple widely distributed regions over time. Such studies will expose relative activity between regions and discover shifts in regional activity in response to different experiences. Here, we performed longitudinal manganese-enhanced magnetic resonance imaging (MEMRI) to measure degrees of brain-wide neural activity in ELA-exposed mice across a series of experiences in adulthood. To ascertain whether ELA resulted in atypical brain activity, results were compared to those of the standard mouse (Std). MEMRI captured activity in the freely moving home cage condition, and short– and long-term after exposure to TMT, a naturalistic predator threat. Images were normalized and aligned then analyzed with statistical mapping and automated segmentation. We found that neural activity in the home cage was greater in ELA compared to Std in multiple striatal-pallidal and hypothalamic regions. Upon acute threat, neural activity in Std increased in these regions to become more similar to that in ELA, while new hyperactive responses in ELA emerged in the midbrain and hindbrain. Nine days after acute threat, heightened neural activity in ELA persisted within locus coeruleus and increased within posterior amygdala, ventral hippocampus, and dorsomedial and ventromedial hypothalamus. These results reveal functional imbalances that arise between multiple brain-systems after ELA, which are dependent upon context and cumulative experiences into adulthood.

**Significance Statement:** Early life adversity (ELA) is a crucial determinant of adult health. Yet the neurobiological basis for this remains elusive. Localized brain regions display atypical neural activity in rodents who experienced ELA, but how this contributes to overall brain state dynamics has been unknown. Here, we used longitudinal manganese-enhanced MRI to detect brain-wide activities altered by ELA in the adult across a series of conditions: freely moving, experiencing threat or its aftermath. Computational analyses revealed that ELA produced widespread dynamic reconfiguration of brain-wide neural activity in response to threat, shown here for the first time. Dynamic brain state remodeling after ELA provides critical insights into human mental health vulnerabilities associated with adverse childhood experiences (ACEs), and suggests plausible targets for intervention.

## Introduction

Early life adversity (ELA) poses risks for neuropsychiatric disorders, especially after a second traumatic event, such as a life-threatening experience (1–5). ELA’s long-term consequences on global neural activity lasting into adulthood could serve as a biological basis for poor outcomes (6–8). We are studying the brain states of adult mice who experienced ELA to test whether neural activity in the home cage has been affected and whether their responses to threat are normal. To ascertain how a potentially normal mouse is affected by ELA into adulthood, we compared ELA-exposed mice to the standard mouse reared under generalized conditions (9, 10).

We invoked ELA in mice using a well-established paradigm: fragmented maternal care induced by limited bedding (11, 12). Most studies using preclinical paradigms focus on localized brain regions with invasive methods (13–27). Only a few have examined ELA’s brain-wide consequences (28–31). Lacking are any studies that examine the brain-wide effects of threat in the adult previously exposed to ELA. Many questions remain that cannot be answered by localized techniques, such as whether brain-wide activity is changed in an adult who has experienced ELA, and whether activity in response to threat is also affected. To investigate these questions and overcome inherent limitations of other techniques, we employed longitudinal manganese-enhanced MRI (MEMRI) together with computational data-driven analysis. This provides quantitative information about global brain activity, which sometimes is described with qualitative terms for directionality or to generalize an aggregate of similar data.

Longitudinal MEMRI detects brain-wide activity before, during and after acute threat in awake freely moving mice (32, 33). Predator odor is a widely accepted naturalistic paradigm to induce innate threat responses in rodents (56–58). MEMRI does not require invasive or destructive procedures (33). Mn(II) is safe – even continuous infusion of low doses has minimal effects on behavior (33, 34). We chose to deliver Mn(II) via two single bolus injections in order to avoid surgical procedures required for continuous infusion and to allow for Mn(II) clearance between imaging sessions (32). Systemic Mn(II) distributes throughout the brain parenchyma (35), enters active neurons through voltage-gated calcium channels (36–38) and accumulates in neurons in an activity-dependent manner (33, 39, 40). MR signal intensity is proportional to the frequency of action potentials (40). Thus, MEMRI represents a quantitative readout of neural activity. Patterns of activity correspond between MEMRI and IEG expression before and after challenge (32, 38, 41). Regional accumulations of Mn(II) throughout the brain produce robust hyperintense MR signals with large effect sizes at 100 μm^3^ resolution (32, 33, 35, 41–43). These signals represent a composite picture of regional neural activity occurring between injection and imaging timepoints. Alignment of multiple MR images, collected repeatedly from large cohorts of individuals as they progress in parallel through a series of experiences, produces a dataset that enables extraction of statistically significant voxel-wise information about brain-wide activity over time, not possible with most other techniques (32, 33, 44–55). Such processing is a well-accepted practice in human imaging that yields much useful information and is routinely repurposed for rodent imaging (52, 56–59). To ensure reliability and reproducibility, computational processing is verified and validated at each step. Thus, longitudinal MEMRI can uniquely image enduring consequences of ELA on brain-wide activity in the adult and its dynamics in response to threat.

Here, we examine the consequences of ELA on adult brain-wide activity during normal home cage exploration in the adult and investigate how this activity changes in response to acute threat, short and long term, with data-driven and hypothesis-based analyses of MEMRI images captured longitudinally. Our alignments and computational processing pipeline were rigorously verified at every step. We validated Mn(II) dose by comparing signal-to-noise ratios across all individuals. We verified that Mn(II) signal replicated neural activity detected by other methods by measuring intensities in circuits known to be involved in predator odor responses and by correlation of Mn(II)-dependent enhancements with IEG (c-fos) expression. We used within– and between-group statistical parametric mapping and automated segmentation with our new *InVivo* mouse brain atlas to quantify relative activity within and between 102 segments brain-wide under each condition. We found that volumes of activities in multiple segments throughout the brain differed significantly between standard mice and those exposed to ELA undergoing the same experiential timeline.

## Results

To witness neural activity in the home cage condition (HC) of adult mice who experienced ELA, and their responses to predator stress short– and long-term, we adapted our longitudinal imaging procedure (**Fig. 1A**) (32). We compared ELA mice (ELA, n = 12) to pool-weaned JAX-reared mice of the same strain (Standard, Std, n = 12) to determine whether brain-wide activity of adult mice who had experienced ELA differed from a phenotypically “normal” adult mouse and the degree of difference.

**Fig. 1.**
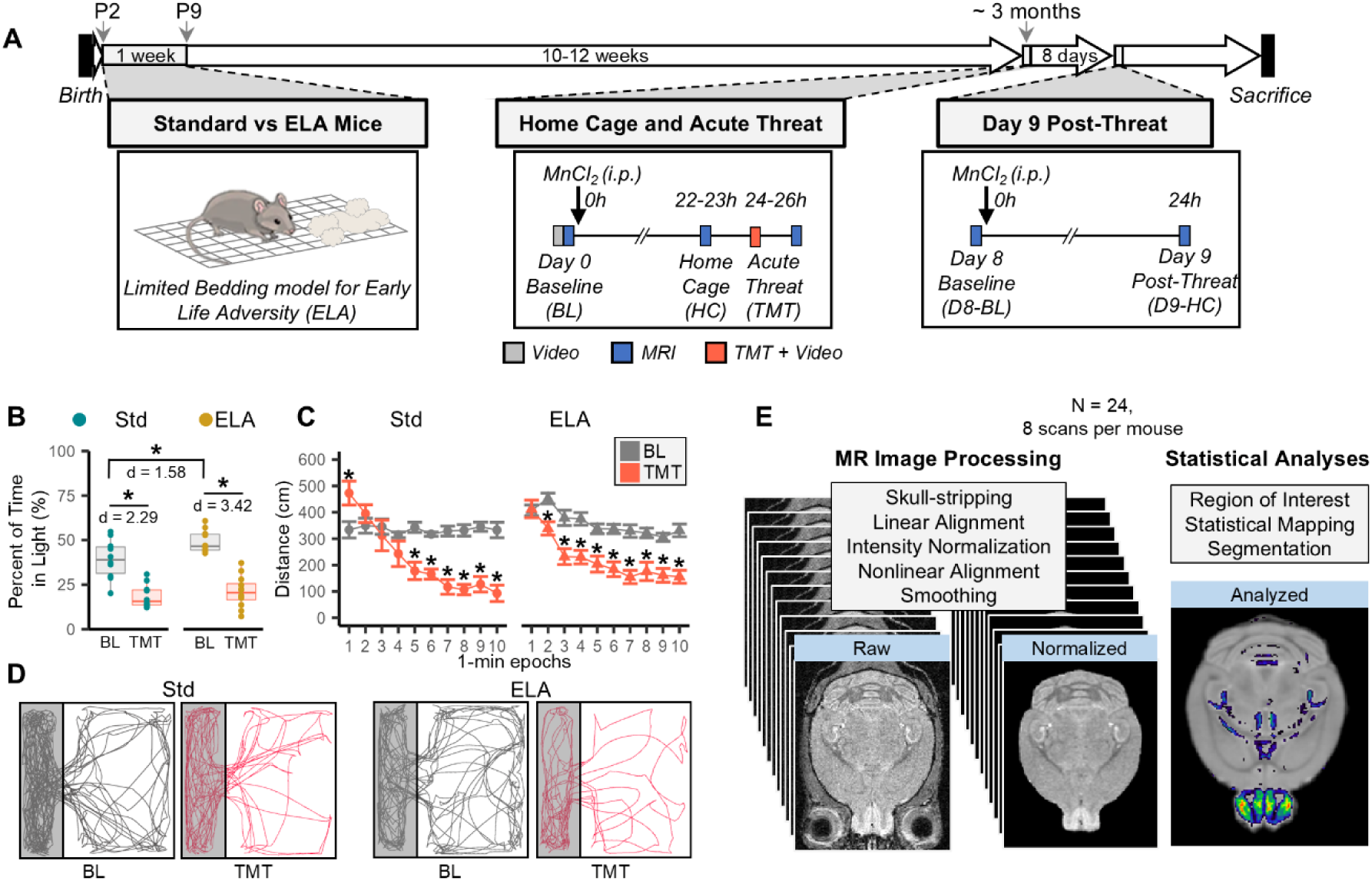
Longitudinal imaging procedures. *(A)* Schematic of the overall timeline (open arrows, top). Boxes diagram events. *Left box*: ELA paradigm; *Middle box*: Home Cage and Acute Threat, timeline of MR and video capture: Baseline, BL; Home Cage, HC; Acute Threat, TMT. *Right box*: Day 8 Baseline, D8-BL; Day 9 Post-Threat, D9-H. *(B-D)* Video tracking results: *(B)* Percent time spent in the light (quartile boxplots, individuals indicated by dot plot overlays) (cyan, Std; yellow, ELA; BL and TMT as per schematic). *(C)* Distance traveled after imaging during TMT exposure. Line graphs of distance travelled per 1-min epochs with error bars representing standard error of the mean. Repeated measures ANOVA with fixed effects of group and condition. For (B-C), paired and unpaired t-tests with FDR correction for multiple comparisons; Cohen’s D effect sizes (d); * p < 0.05, n = 22 (10 Std, 12 ELA). *(D)* Representative tracking of one mouse from each group, Std and ELA, at BL (gray) and TMT (red). Shaded areas represent dark side of the arena. *(E)* Overview of MR data processing and analysis used in this study (also see ***SI Appendix* Fig. S1-2**) (33, 50, 52).

### Successful Replication of Behavioral Paradigms

To test whether our protocols for these two paradigms, fragmented care (ELA) and predator stress (TMT), faithfully reproduced previously reported behaviors (60–62), we compared videos of adult mice as they explored a custom arena (**Fig. 1B-D**). For the effect of ELA experience, we measured the time ELA mice spent in the light at baseline (BL) and compared that to Std mice. As expected, adult ELA mice spent 10% more time in the light at BL than Std, demonstrating a persisting effect of ELA experience in an unprovoked situation (p < 0.05 FDR, Cohen’s d = 1.58) (**Fig. 1B**). For the effect of TMT, we measured time spent in the light and compared between BL and TMT. As expected, TMT decreased time spent in the light in both ELA and Std mice by more than 20%, demonstrating an acute effect of TMT (p < 0.05 FDR, Cohen’s d_Std_ = 2.29, d_ELA_ = 3.42). To assess mouse recovery from MR anesthesia before being presented with TMT, we measured distance traveled in 1 min epochs and tracked movements from the beginning of a 10 min exposure to TMT (**Fig. 1C-D**). During the first minute of TMT-exposure, distance traveled was equal to or greater than that at BL. Motility then decreased by 50% or more over the 10 min TMT-exposure period (p < 0.05, FDR). These data support the contention that our procedures faithfully reproduced ELA and TMT paradigms, giving us confidence in assigning behavioral characteristics to unknown brain-wide activity.

### Longitudinal Imaging

Our MR imaging protocol captures 8 sequential images across 12 individuals for each group (ELA and Std), yielding a total of 192 raw images with >2 million voxels each (**Fig. 1E**). To ensure no differences between groups were introduced by processing, all images were processed as a batch through our semi-automated processing pipeline (32, 33). This pipeline yielded a combined dataset of skull-stripped, intensity normalized and spatially aligned images ready for statistical evaluation. Quality assurance considerations included reviewing imaging parameters, performing SNR measurements to confirm similar Mn(II) doses, visualizing modal scaling output in voxel-by-intensity histograms to monitor equivalent intensity normalization, calculating Jaccard Similarity and Mutual Information Indices for alignment accuracy, and determining that smoothing does not significantly change regional signal intensities. All images reported here met our quality assurance standards (***SI Appendix* Fig. S1-2** and **Table S1**).

### Brain States in Mice Exposed to ELA

To detect Mn(II)-enhancements in ELA mice across conditions from the home cage (HC) to post-threat (TMT) to nine days after threat (D9), we compared post-Mn(II) images for each condition to each mouse’s respective pre-Mn(II) image using voxel-wise statistical parametric mapping (SPM) (**Fig. 2**). SPM of images at HC revealed large (16.9% of total) volumes of statistically significant signals in multiple brain regions identified by anatomy, demonstrating that the unprovoked ELA brain was already highly active (T = 2.72, p < 0.01, cluster corr.) (**Fig. 2A**). Visual inspection of SPMs revealed larger volumes of signal within regions of the olfactory bulb, striatum/pallidum, hypothalamus, hindbrain, and cerebellum, and smaller volumes within subregions of hippocampus, amygdala, and hindbrain. Immediately after TMT, volumes of statistically significant voxels increased by +1.4% overall. SPM volumes within these regions were maintained or expanded, with sizable increases in the mid– and hindbrain (e.g., LC) (**Fig. 2B**). By D9, activity decreased in olfactory, hindbrain, and cerebellum, or increased in hippocampus, amygdala, and midbrain (e.g., PAG), leading to an overall decrease of –2.9% from HC (**Fig. 2C**). Other subregions also changed, such as a decrease of the volume of statistical signal in the lateral septum (LS) and appearance of new signal in the paraventricular nucleus of the thalamus (PVT). Direct SPM paired t-test comparisons between ELA post-Mn(II) images at each condition added to these findings from activation volumes, to also identify statistical differences in signal intensities within many regions across conditions (***SI Appendix* Fig. S3**). These findings indicate that ELA-exposed brains are already highly active before a second provocation in adulthood, and that short– and long-term brain responses to TMT have unique patterns of activity, with regions emerging or disappearing between conditions.

**Fig. 2.**
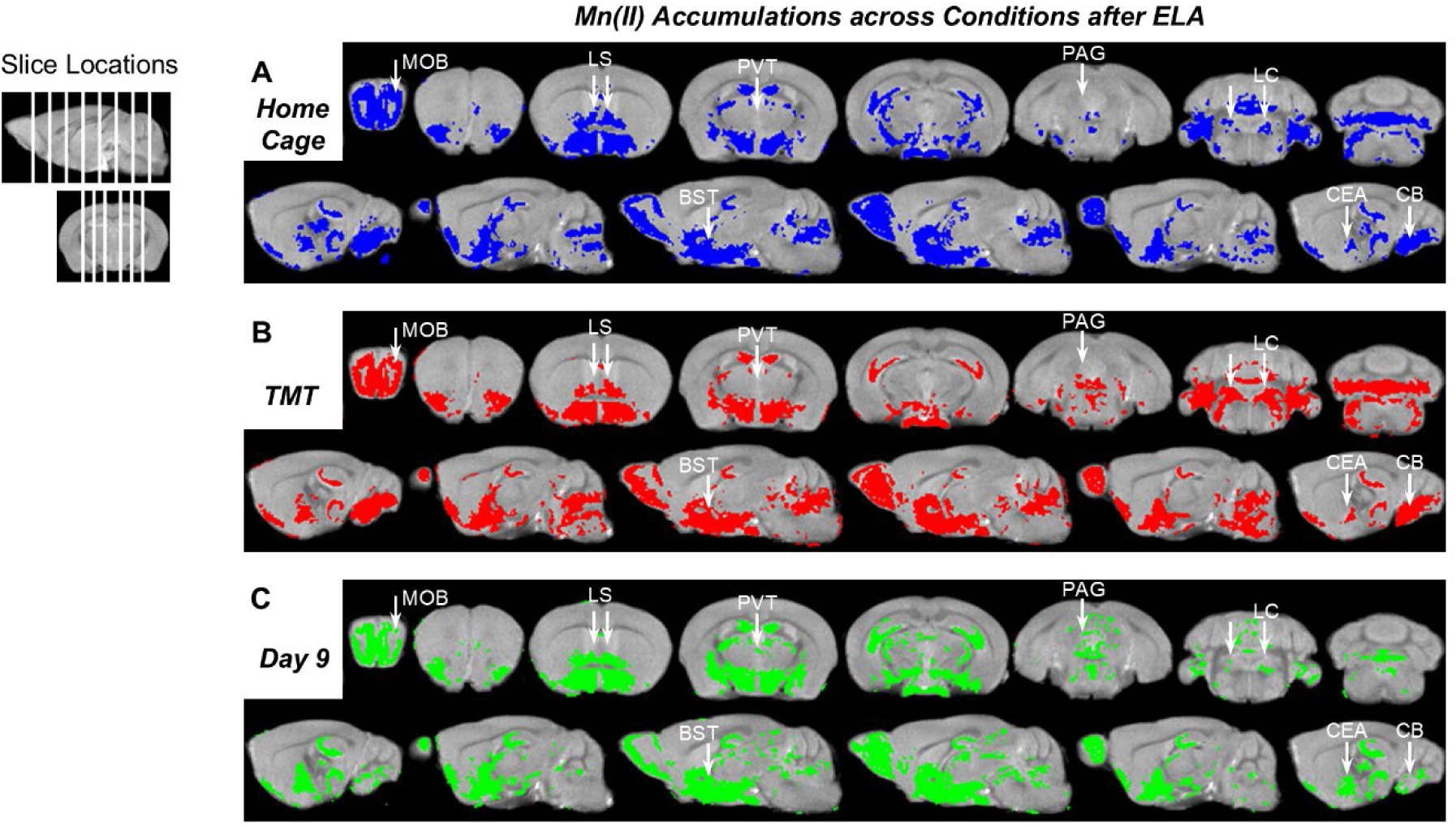
Statistical parametric maps of MEMRI in ELA mice change as they progress from home cage to acute threat to post threat. *(A-C)* Shown are statistical maps of voxel-wise paired t-tests of post-Mn(II) images of ELA mice at each successive condition compared to their pre-Mn(II) images. Maps are overlaid on the grayscale dataset average of all pre-Mn(II) images (coronal slices, top rows; sagittal, bottom rows). Slice position is indicated on the grayscale image (top left, white lines). White arrows indicate notable differences between conditions. MOB, main olfactory bulb; LS, lateral septum; BST, bed nuclei of the stria terminalis; PVT, paraventricular thalamus; HIP, hippocampus; LC, locus coeruleus; CEA, central amygdala; and CB, cerebellum. For t-tests, n = 12, p < 0.01, T = 2.72, cluster corr. For comparisons between conditions of ELA mice see ***SI Appendix***, **Fig. S3**.

### Magnitude of Signal Enhancements

To determine whether these statistical maps actually represent changes in intensity values across conditions, we measured intensities within 3 x 3 x 3 voxel regions of interest (ROI) in TMT-responsive regions and compared between HC and TMT (**Fig. 3A**, and ***SI Appendix* Tables S2** and **S3**) (63, 64). Our voxel-wise SPM of ELA images at HC and TMT revealed significantly enhanced signal intensities after TMT in all four expected regions (p < 0.05 cluster corr.) (**Fig. 3A**, *top*). At HC, SPMs show that dlOB-DII and vlAON were already enhanced and only minimally changed across conditions. Region of interest (ROI) measurements within these locations revealed that SPM signals at HC represented large signal differences over pre-Mn(II) (6.8%, dlOB-DII; 8.0%, vlAON) with standard deviations of similar magnitude, leading to large Cohen’s D effect sizes (d = 1.6, dlOB-DII; d = 1.5, vlAON; p < 0.05 FDR). Standard deviations are shown in **Fig. 3A** and listed in ***SI Appendix* Table S3**. In aOC and aCEA, SPM signals were minimal at HC and increased from HC to TMT (**Fig. 3A**, *top*). These SPM signal changes were recapitulated in ROI comparisons, with aOC and aCEA showing small magnitudes of signal enhancement over pre-Mn(II) at HC (1.4% and 3.0%), and larger magnitudes at TMT (3.5% and 6.2%) (**Fig. 3A**, *bottom*). Thus, in this case SPMs represented signal intensity differences. Signal intensity increased in all four regions expected to respond to TMT (+2.1-3.9%, d = 0.85-1.20, p < 0.05 FDR), demonstrating robust TMT-induced enhancement.

**Fig. 3.**
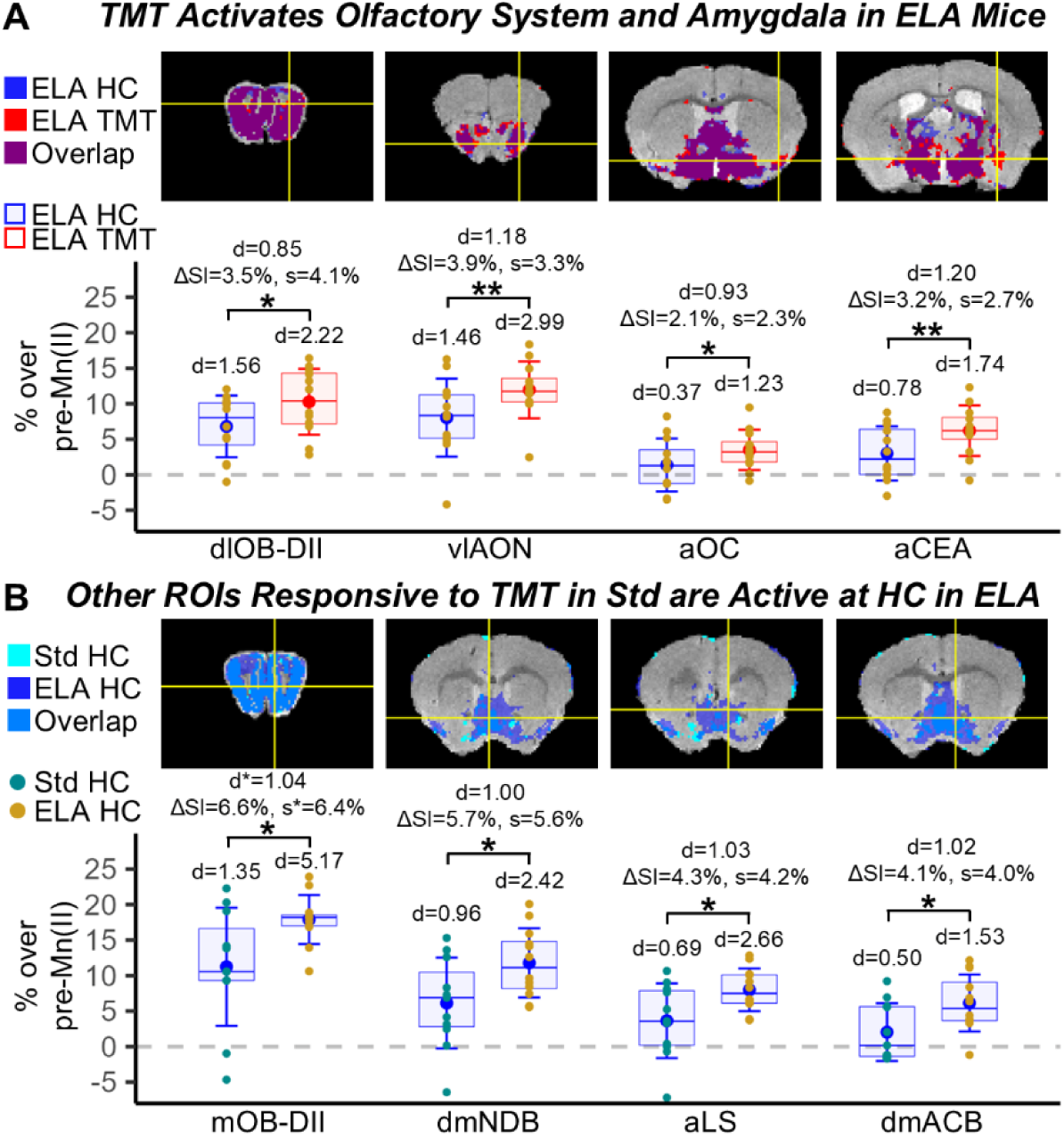
Measurements of MEMRI signal intensities in candidate regions of interest in systems known to be involved in predator odor-type threat responses. (A) Top: Representative slices from two voxel-wise statistical maps (ELA HC & ELA TMT) of post-Mn(II) > pre-Mn(II) superimposed onto the grayscale InVivo atlas. SPM paired t-tests: p < 0.01 cluster corr. ROI locations selected where TMT signal did not overlap with HC (yellow crosshairs). Bottom: Quartile boxplots of signal intensity as a percent over pre-Mn(II) measured in each ROI. Error bars: Mean ± standard deviation. Effect sizes of the differences in signal between conditions at each ROI (Cohen’s D, d), average signal intensity differences (ΔSI) between the two conditions, and standard deviations of ΔSI are indicated. Paired t-test, * p < 0.05, ** p < 0.01, FDR. dlOB-DII, dorsal lateral DII olfactory subdomain; vlAON, ventrolateral anterior olfactory nucleus; aOC, anterior olfactory cortex; and aCEA, anterior central amygdala. (B) Top: Similar comparison but between groups at one condition, HC. Slices of paired t-tests of HC greater than pre-Mn(II) from either Std or ELA are superimposed on the grayscale atlas. Crosshairs indicate ROI locations, selected where ELA signal did not overlap with Std. Bottom: Quartile boxplots of unpaired t-tests of ROI measurements. Error bars: Mean ± standard deviation. Welch’s t-test and non-pooled variance-based d* and s* were used for mOB-DII. * p < 0.05, FDR. mOB-DII, medial DII olfactory subdomain; dmNDB, dorsomedial diagonal band nucleus; aLS, anterior lateral septum; dmACB, dorsomedial nucleus accumbens. See ROI coordinates and summary statistics in ***SI Appendix,* Tables S2-4**.

Statistical mapping of ELA at HC (**Fig. 2** above) indicated signal enhancements within regions normally only responsive after TMT in Std (32). To determine the degree of signal intensity differences between ELA and Std at HC and test whether ELA signal truly exceeded Std in these regions, we measured and compared signal intensities within 3 x 3 x 3 voxel ROIs (**Fig. 3B**, ***SI Appendix* Tables S2** and **S4**). Magnitudes of signal enhancement at HC over pre-Mn(II) corresponded to the presence or absence of significant SPM signal for both Std and ELA (non-enhanced: SI = 2.0-3.6%, d = 0.5-0.7; enhanced: SI = 6.1-17.9%, d = 0.96-5.17). Direct comparisons between groups revealed that ELA intensities were heightened at HC in all four measured regions (SI = 4.1-6.6%, d > 1.00, p < 0.05 FDR). Standard deviations are shown in **Fig. 3B** and listed in ***SI Appendix* Table S4**. Thus, ELA signal intensities within threat-responsive regions exceeded that of Std even before threat. ELA also displayed a lower variance than Std.

Together these observations support the conclusion that differences in SPM-identified activation volumes, at p < 0.01 and cluster size ≥ 8 voxels, reliably represent intensity differences between post-Mn(II) images. However, when SPM analysis reveals overlapping anatomy of activation volumes between groups or conditions, direct measurements of post-Mn(II) intensities may be necessary to uncover potential differences in activity levels that are not detected by activation volumes.

### Mn(II)-Enhanced Signal Corresponds to c-fos Expression

We next sought to test whether differences in Mn(II)-enhancement corresponded quantitatively to differences in neural excitation, as detected by c-fos expression (33, 41, 66, 67) (**Fig. 4**). We previously reported a correspondence between c-fos and MEMRI before and after challenge (predator odor) and found similar changes in activity between the two modalities (32). Here, we compare intensity values from *in vivo* longitudinal MEMRI with positive nuclei counts from c-fos immunohistochemistry, which is an endpoint where longitudinal dynamics are not feasible. A subset of the mice that underwent the longitudinal MEMRI protocol were processed for c-fos immunohistochemistry (n_Std_ = 5 and n_ELA_ = 6). We selected pairs of regions in sections stained for c-fos where anatomy could be cross-correlated with MR slices for each mouse: hypothalamus (HYP) and thalamus (THA); superior central raphe (CS) and midbrain (MB) (**Fig. 4A-B**). First, we compared regions for each modality separately (**Fig. 4C-D**). Statistical comparison of Mn(II)-enhanced signal intensities in HYP and CS were statistically greater than in THA and MB, respectively (**Fig. 4C**, p < 0.001). Numbers of c-fos+ nuclei revealed similar differences between regions (**Fig. 4D**, p < 0.001). Robust effect sizes were observed for both modalities (c-fos: MEMRI: d = 1.88-6.37; d = 3.75-5.03). Statistical differences in MR signal intensities from this subset of mice were replicated in a second subset from each group, demonstrating test-retest reliability (d = 2.94-8.43, p < 0.001, FDR) (***SI Appendix* Fig. S4**). To test further whether intensities correlated linearly with c-fos counts, we calculated Pearson’s pairwise correlation coefficient for MEMRI versus c-fos data across the four regions with bootstrap resampling and permutation testing. In both groups, correlations were statistically significant (Std: r = 0.636, p < 0.0001; ELA: r = 0.582, p < 0.0013) (**Fig. 4E** and ***SI Appendix* Fig. S5**).

**Fig. 4.**
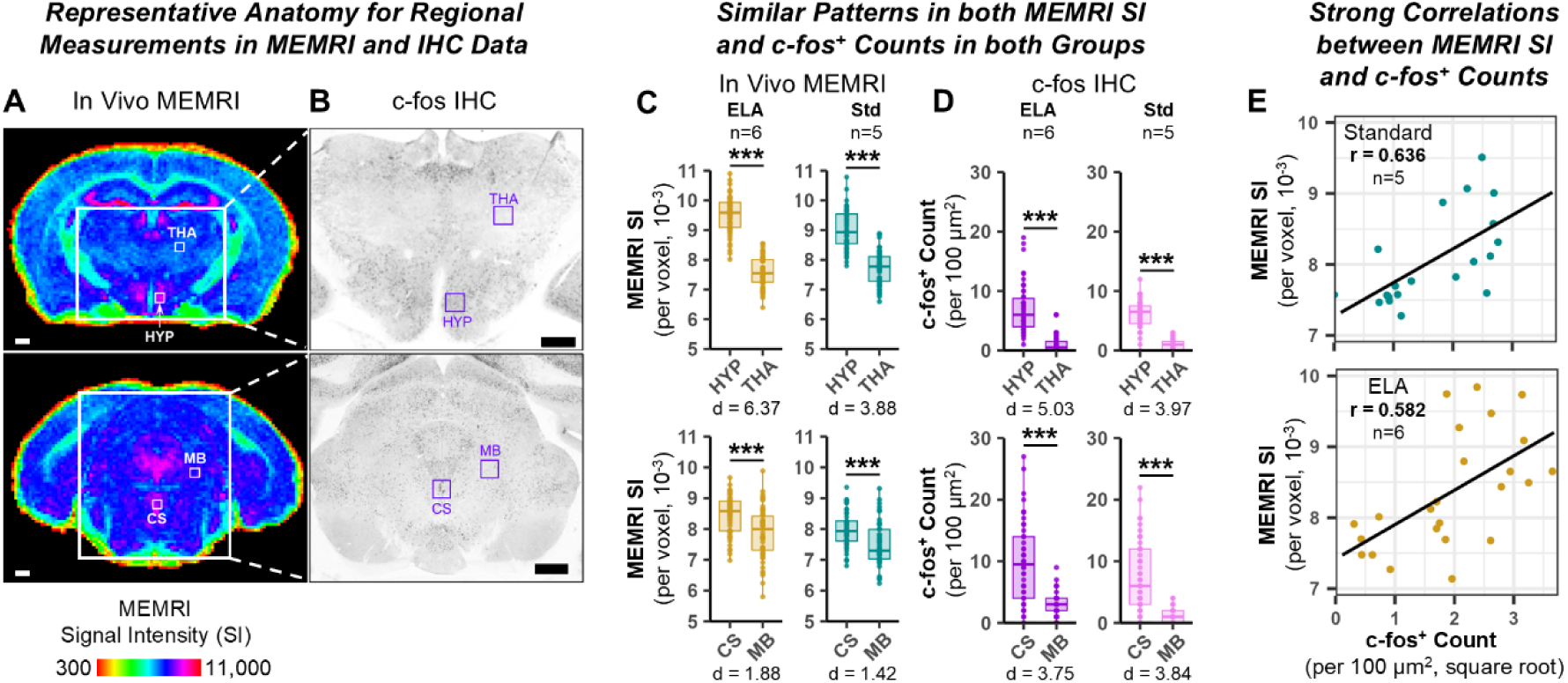
MEMRI and c-fos staining correlate. (*A*) Coronal slices of the average intensities of day-9 post-Mn(II) images (color code, bottom) (n = 6). Minimum (300 a.u.) based average noise and maximum (11,000 a.u.) from intensity histograms. Small white boxes indicate regions (3 x 3 x 1 voxels) where signal intensities (SI) were measured. HYP, hypothalamus; THA, thalamus; CS, superior central raphe; MB, midbrain. Mag bar = 500 μm. (*B*) Two representative histological sections from one of the mice contributing to *(A)* stained for c-fos corresponding to anatomy of slices in *(A)* (large white boxes). Purple boxes indicate locations of 3 x 3 grid of 100 μm^2^ squares where numbers of c-fos+ nuclei were counted. Mag bar = 500 μm. See ***SI Appendix* Fig. S5A-B** for corresponding images from Std. *(C)* Statistical comparisons of MEMRI SI in four locations for both groups. Shown are quartile boxplots with dot plot overlays of MEMRI SI from ELA (yellow, left) and Std (cyan, right). Paired t-tests were performed between regions within the same slice. Note large Cohen’s D effect sizes (d) for paired t-tests between active and inactive regions. For evidence of test-retest reliability of MEMRI SI, see ***SI Appendix* Fig. S4**. *(D)* c-fos counts paralleled MEMRI SI with high or low numbers of c-fos+ nuclei. Shown are quartile boxplots with dot plot overlay of raw c-fos+ counts from ELA (purple, left) and Std (pink, right). A square root transformation linearized c-fos+ counts for statistics. Note similar large Cohen’s D effect sizes (d) between active and inactive regions. *(E)* Strong Pearson correlations between MEMRI SI and numbers of c-fos+ nuclei (linearized) when paired by individual. Scatter plots with linear regressions (black lines) of MEMRI SI (vertical axes) by c-fos+ counts (horizontal axes). Points represent the average values for each of the 4 regions across 5 Std (*top*, *cyan*) and 6 ELA (*bottom*, *yellow*) brains. Note strong correlations in both groups (r_Std_ = 0.636, r_ELA_ = 0.582). For paired t-tests in *(C-D)*, *** p < 0.001, FDR corrected. For *(E)*, p < 0.001 by permutation testing and bootstrapping. Also see ***SI Appendix* Fig. S5C-D**.

A between-group comparison of c-fos and MEMRI at D9 for each region found that both modalities found increased activity in HYP (d_c-fos_ = 0.38; d_MEMRI_ = 2.21) and CS (d_c-fos_ = 1.07; d_MEMRI_ = 2.02) in ELA relative to Std. While MEMRI differences were statistically significant in HYP and CS (p < 0.05, FDR), c-fos differences were not, demonstrating that MEMRI could detect statistically significant differences with only 5-6 mice possibly due to larger values and smaller variance in MEMRI data. Comparison of c-fos at D9 with MEMRI from all three conditions in these regions revealed statistically significant correlations (r > 0.582, p < 0.0013). This result shows that the relative levels of activity were consistent in these regions across the longitudinal imaging procedure. Thus, our data provide evidence that differences in the degree of MEMRI signal intensities reflect differences in c-fos expression for the most part.

### Deeper Dives into Brain States of Adult Mice Exposed to ELA

To assign neural activity in ELA mice to specific anatomical regions, we aligned statistical maps of enhanced voxels with our *InVivo* Atlas (***SI Appendix* Table S5**). Then we quantified activity in those anatomically defined segments at each condition with a new approach, fractional activation volumes (FAV) (32) (**Fig. 5**). The FAV is the fraction of activated to total voxels in each segment. Activated voxels were those found to be statistically enhanced in SPM comparisons of post-Mn(II) to pre-Mn(II) images at each condition. FAVs were visualized as column graphs. Comparing column graphs between conditions demonstrated changes in FAVs as mice progressed through successive conditions. For conciseness, we summarize key regional differences from this quantitative analysis using qualitative terms for direction, magnitude, and aggregate patterns. Quantitative information underlying these terms is available in ***SI Appendix* Tables S6-8**.

**Fig. 5.**
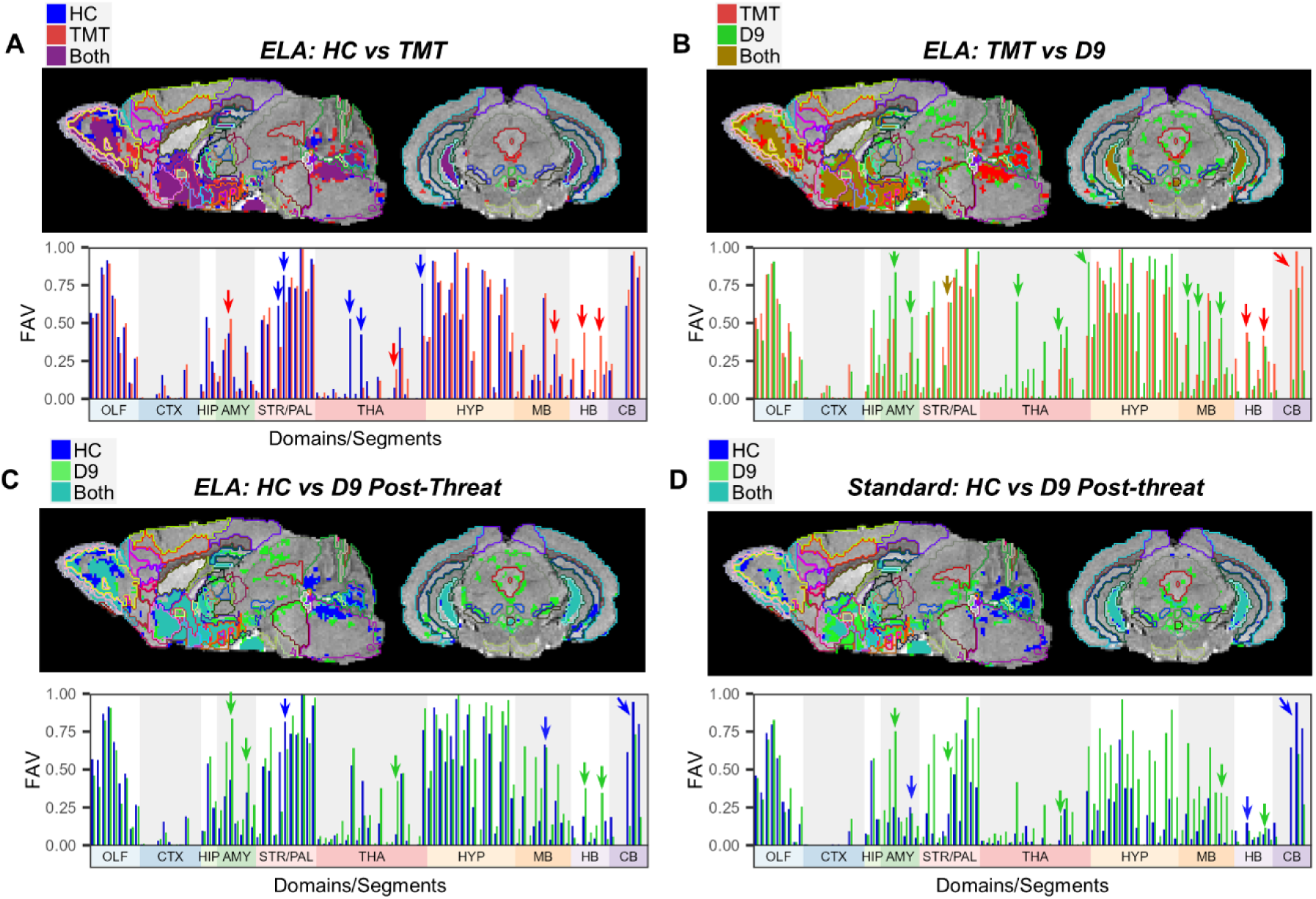
Statistical maps show progressive modifications of activity across sequential conditions. *(A-D)* Voxel-wise paired t-tests of post-Mn(II) compared to pre-Mn(II) images at two conditions: in ELA mice, *(A)* HC (blue) and TMT (red), *(B)* TMT (red) and D9 (green), and *(C)* HC (blue) and D9 (green); and in Std mice, *(D)* HC (blue) and D9 (green). *Top*: Pairs of statistical maps were projected onto the grayscale *InVivo* Atlas with segments outlined to show the anatomical overlaps between conditions. *Bottom*: Corresponding column graphs show segment-wise fractional activation volumes (FAV, vertical axes) quantified from segmentations of statistical maps. Anatomical domains (color-coded on horizontal axes) are divided into 102 segments. Arrows point to examples of segments with FAV alterations between conditions and color-coded to indicate direction of change. For paired t-tests, p < 0.01, cluster *corr.* for multiple comparisons. ELA, T(_11_) = 2.72, d ≥ 0.79, n = 12; Std, T(_10_) = 2.76, d ≥ 0.83, n = 11. See ***SI Appendix* Table S5** for list of domains and segments, and **Table S6A-B** for FAV values.

Overlays of statistical maps show that much of the global activity in ELA at HC (**Fig. 5A, blue**) versus pre-Mn(II) coincides with that of TMT (**Fig. 5A, red**) in the same group (**Fig. 5A, overlap purple**) (p < 0.01 cluster corr., d = 0.79-8.19) (***SI Appendix* Table. S6A**). Some regions expand while a few others contract from HC to TMT. By FAV analysis of ELA, we see that at HC large FAVs were prominent in various domains (OLF, STR/PAL, HYP, and CB) with some segments (MOBmi, 92%; MS, 100%; BST, 92%; and AHN, 97%) almost totally active. Immediately after TMT, FAV values were retained or increased, with the largest increases in CEA (+9%) and LC (+42%). Several STR/PAL and THA segments decreased in FAV after TMT, including LSc (−28%), LGd (−53%) and MH (−34%). No relationships between segment volume and FAV were observed. Differences in signal intensities between conditions in these segments were also statistically significant by SPM comparisons between successive post-Mn(II) images, with the greatest differences in the LC (p < 0.05 cluster corr.) (***SI Appendix* Fig. S6** and **Table S6B**).

We then compared SPM overlays and segmental FAVs between TMT (**Fig. 5B, red**) and D9 (**Fig. 5B, green; overlap, brown**) of ELA mice to learn how TMT affected brain-wide activity long term (**Fig. 5B**). Segmentation analysis revealed globally that FAV decreased in three domains (OLF, HB, and CB; –14% on average) with the most substantial change in CB (−19%). Most other segments either retained or further increased FAV, particularly within AMY, THA, HYP and MB domains (+11% on average). In many segments where FAV had increased following TMT, FAV at D9 either mostly persisted (LC, 34%) or increased (CEA, +29%), suggesting a longer lasting effect of the threat experience. Segments where FAV decreased acutely after TMT typically remained low or further decreased (LSc, –12%). Others increased at D9 (MH, +14%). These results suggest that brain activity in some segments does not return to pre-threat levels.

To test whether activities return or not to pre-threat levels, we compared HC with D9 using a similar strategy in both ELA and Std mice (**Fig. 5C and 5D**, also ***SI Appendix* Table S6A**). Both overlays of SPMs (**Fig. 5C and 5D, overlap, turquoise**) and columns graphs of FAV show that at D9 after threat, activity in both ELA and Std was increased substantially compared to HC in many domains (AMY, STR/PAL, HYP, and MB). The magnitude of FAV increase in these domains was twice as large on average in Std than in ELA (+28% vs +11%). This pattern also appeared in pairwise comparisons of post-Mn(II) intensities between conditions (***SI Appendix* Fig. S7** and **Table S6B**). Neither the brain-wide activity of ELA nor Std returned to pre-threat levels at D9. These data suggest regions in the brain remain hyperactive one week or more after acute threat.

Visual comparison of pairwise statistical maps within group at HC and D9 suggested statistical differences between groups. At HC, before threat, FAVs of ELA were substantially greater than those of Std across brain segments (31% vs 17% on average). These larger FAVs in ELA could not be attributed to procedural differences, such as MnCl_2_ dosing, MR parameters, imaging timing, image processing, segmentation analysis (***SI Appendix* Fig. S2** and **Table S1**), differences in pre-Mn(II) signal intensities between groups (***SI Appendix* Fig. S8**) or differences in anatomy between groups (***SI Appendix* Fig. S9**). While Std exhibited larger FAV increases from HC to D9 than ELA in most segments (+12% vs +5% on average), ELA exhibited larger increases than Std in a subset of segments, including RE (+35% vs +16%) and LC (+34% vs +4%). In other segments, ELA displayed the opposite directional change from HC to D9 compared to Std, including PA (+19% vs –4%), LSr (−18% vs +31%), IPN (−2% vs +33%), and PCG (+19% vs –5%). Except for IPN, these segments also responded acutely to TMT in ELA (**Fig. 5B**). These findings suggest that brain-wide activity and its evolution from threat-naïve (HC) to threat-experienced (D9) conditions differs between groups and that ELA mice resemble a post-threat-like state already at HC. Furthermore, those segments that respond to TMT in ELA appear to be major contributors to differences between groups at D9. These observations prompted direct between-group statistical analyses for each condition.

### ELA Affected Many Segments Within Each Anatomical Domain

To determine whether signal intensities between ELA and Std differed statistically between groups, and to map the anatomical distribution of those differences, we performed direct SPM comparisons. First, we performed an unpaired t-test comparing baseline pre-Mn(II) images between groups, which revealed small intensity differences (***SI Appendix* Fig. S7**). For subsequent comparisons, post-Mn(II) images were adjusted to correct for these baseline differences. We then compared ELA and Std post-Mn(II) images at each condition by an unpaired SPM t-test (p < 0.05, cluster corr., d = 0.70-5.30) (**Fig. 6** and **Movie S1**). Segmentations of SPM difference maps at each condition were used to obtain the number of statistically enhanced voxels (in either ELA > Std or Std > ELA) per segment, which were then divided by the total number of voxels in each segment producing the fractional difference volume (FDV) (**Fig. 6A-D**, and ***SI Appendix* Table S7**). At all three conditions, ELA exhibited greater signal intensities than Std (i.e., heightened activity) in most segments but to a lesser extent after TMT (HC, 17%; TMT 11%; D9, 12% on average), supporting observations from the within-group SPMs shown in **Fig. 5**. From HC to D9, the total volume of enhanced voxels in ELA relative to Std decreased by 4.4%.

**Fig. 6.**
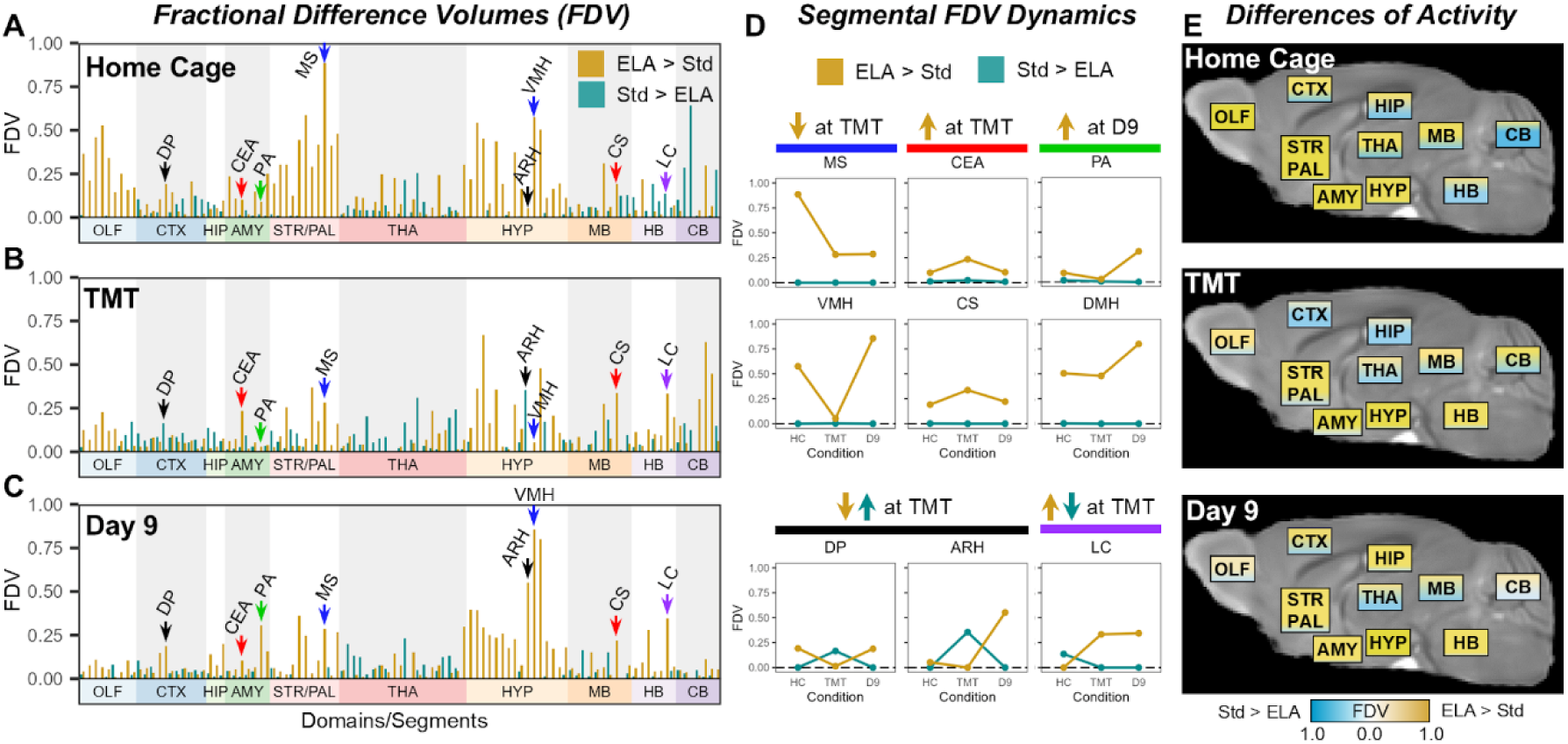
ELA brain states differ from Std at every condition. *(A-C)* Fractional Difference Volumes (FDVs) from segmentation of unpaired SPM t-tests between signal intensities of ELA and Std at *(A)* HC, *(B)* TMT and *(C)* D9 conditions. All ELA images were adjusted for the voxel-wise average differences between groups’ pre-Mn(II) images. FDV color-code represents where ELA signal intensities were heightened (yellow) or diminished (blue) relative to Std. Arrows in column graphs point to example segments with notable FDV dynamics as in *(D)*: blue, MS and VMH; red, CEA and CS; green, PA and DMH; black, DP and ARH; and purple, LC. *(D)* Shown are examples of various dynamics in FDV in 9 segments, which were highlighted by arrows in column graphs *(A-C)*. Line graphs show FDVs (vertical axes) of each segment across three successive conditions (horizontal axes). Graphs are grouped and color-coded according to the dynamical pattern of ELA and Std FDV values. *(A-D)* Unpaired t-tests, p < 0.05, cluster corr.; HC: T(_21_) = 1.72, n_Std_ = 11, n_ELA_ = 12; TMT and D9: T(_22_) = 1.72, n = 12 per group. *(E)* Three schematics summarizing differences in activity between ELA and Std across the 10 segmented anatomical domains at each condition: HC (*top*), TMT (*middle*) and D9 (*bottom*). Each color gradient corresponds to magnitudes and direction of segmental fractional difference volumes (FDV), and proportion of coloring indicating proportion of segments following that pattern within each domain; See **SI Appendix Movie 1** for a slice-by-slice walkthrough of between group SPMs at each condition projected on brain anatomy, **Table S8** for FDV values and **Table S6** for list of domains and segments.

Closer examination of FDVs within different anatomical domains identified heightened activity at HC in ELA compared to Std within OLF (29% on average), STR/PAL (40% on average), and HYP (26% on average) (**Fig. 6A**, yellow columns). At TMT, differences in these domains decreased (−17% on average), although heightened HYP activity in ELA was somewhat preserved (−7%) and new differences became noticeable in the CB (+16%) (**Fig. 6B**). At D9, heightened activity in the HYP domain of ELA returned or even increased over HC (+13% on average) (**Fig. 6C**). While heightened activity was present in most HYP segments for each condition (26% at HC vs 19% and 32% at TMT and D9), many other domains in ELA exhibited smaller volumes of heightened activity over Std at TMT and D9, particularly segments within STR/PAL (40% at HC vs 14% at both TMT and D9). In contrast, Std exhibited heightened activity compared to ELA within only a few segments of CTX, THA, HYP, MB and CB domains (4.6% on average across all segments and conditions), with the most differences occurring at TMT (6%) (**Fig. 6A-C**, blue columns). Thus, ELA in general displayed heightened brain-wide activity over Std at HC, with FDV for ELA > Std of 10% or greater in 53 of 102 total segments. Of the 53 segments, the degree of differences lessened upon experiencing threat in most but not in all (TMT, 31; D9, 30). Next, we will take a deeper dive into segmental activity dynamics.

Examining segment-wise FDV across conditions revealed at least five patterns of segmental dynamics (**Fig. 6D**). To facilitate interpretation of activity differences across conditions, we plotted FDVs indicated by arrows in **Fig. 6A-C** as line graphs (**Fig. 6D**) and FDV values listed in ***SI Appendix* Table S7**. These graphs exemplify five dynamic patterns across conditions of heightened activity in ELA > Std (yellow) or Std > ELA (blue). *One* pattern was heightened activity in ELA at HC, which became more similar between groups at TMT (**Fig. 6D**, blue), as exemplified by MS and VMH and observed across 26 segments. Some of these segments (18) continued to be similar between groups at D9 (e.g., MS) while others returned (7) to heightened activity in ELA (e.g., VMH). A *second* pattern followed an opposite dynamic in response to TMT. In these segments (8), volumes of heightened ELA activity expanded acutely with TMT and returned to HC levels at D9 (CEA and CS, red). A *third* pattern appeared in a few segments (8), where FDVs of ELA rose only at D9 (PA and DMH, green). A *fourth* pattern emerged in 5 segments (e.g., DP and ARH, black). Similar to pattern one or three, ELA activities in these segments were greater than of Std at HC or D9. Dissimilar to pattern one or three, ELA activities decreased in these segments below those of Std at TMT. A comparison between male and female found that ELA females might contribute to between-group differences at D9 in VMH, ARH and PA more than ELA males, with sex-differences in PA also present in pre-Mn(II) images (***SI Appendix* Fig. S10**). A single segment followed a *fifth* pattern (LC, purple), where activity was greater in Std at HC and then reversed after TMT through D9 to become heightened in ELA. Together these data demonstrate that brain-wide activity of ELA mice is reconfigured across conditions and that dynamics in response to threat, both brain-wide and segment-wise, differ dramatically from Std.

## Discussion

In this study we mapped the spatial-temporal transformations that ELA imposes on brain-wide activity, as detected initially when in a home cage (HC) environment and then short and long-term after challenge by an acute threat as compared to a Standard, phenotypically “normal,” mouse. Three different brain states (68), as defined here by widely distributed patterns of fractional activation volumes among segments, emerged within each group. In all three conditions, brain states of adult mice reared with the fragmented maternal care model for ELA differed from brain states of adult Std mice, with greater overall neural activity after ELA (**Fig. 6E** and ***SI Appendix* Fig. S11**). Even without provocation, ELA mice at HC had heightened fractional activation volumes in regions known to be activated by predator odor or altered in reward processing: olfactory and hypothalamic segments (67, 69–74), as well as striatum-pallidum (32, 75, 76). Distribution of heightened activity expanded immediately after threat to engage subcortical structures as well as monoaminergic nuclei and deeper brainstem segments in ELA, as also occurred in Std (32). Neither group regained HC activity patterns nine days after threat. Across all conditions, activity in hypothalamus of ELA was higher than Std. At D9, new differences emerged in the ventral hippocampus and posterior amygdala, where activity was greater in ELA. Our longitudinal brain-wide imaging strategy coupled with our computational processing and analytics empowered us to follow these brain states in the same individuals as they progress through this series of experiences in sample sizes sufficient for statistical significance. Our processing pipeline, aligning 192 whole-brain images captured with 100μm^3^ resolution, allows precise localization of statistically significant activity common across multiple individuals. This unique capability is not possible with most other physiological approaches.

We found dynamic changes in segment-wise fractional activation volumes between conditions throughout the brain in both cohorts. When a segment transitions from lower to higher FAV/FDV, we predict that segment to have a more dominant functional role within its connectome. For example in ELA, the increased volume of activity In CEA (from 43% to 53%) as the mouse progresses from home cage to immediately after threat may contribute to heightened activity throughout threat circuity (77). That ELA mice sustained heightened CEA activity at all three conditions, suggests that ELA leads to persistent engagement of threat circuitry. This sustained engagement of threat circuitry after ELA may sensitize CS and LC to threat, promoting monoaminergic modulation of segmental activities toward the patterns observed at D9 (78–81). Differing levels of activity among segments of a single network across different conditions could produce different internal states (82, 83) and/or outcomes from that same network, be they actions, decisions, emotions, thoughts, memories, or creative works, such as music, poetry, or art.

Results presented here should be considered in light of several limitations. 1) Theoretically, small variations in rearing environment could contribute to differences between groups not attributable to ELA. However, multi-site studies in mice find that different breeding/rearing facilities have no effect on stress reactivity (84) and minimal explanatory value for at least 14 other physiological measures (85). JAX-reared mice are the phenotypical standard and the most commonly used mouse by researchers globally (9), making them an invaluable comparison group to generalize findings across research facilities (86). 2) Although we found minimal sex differences between male and female ELA mice, we cannot fully attribute heightened activity in HYP and vHIP/AMY segments to ELA because we only had a small sample size and because our Std group was all male. Other studies suggest a combinatorial effect of ELA and sex is likely within these threat responsive regions (73, 87, 88). 3) Small anatomical variations could have influenced our results (89–92). However, our Jacobian deformation analysis reported minimal alterations in anatomical volumes, and our SPM analysis accounted for any signal differences by preserving intensity concentrations within voxels (93, 94). 4) Atlas segmentation yields volumes that vary widely, ∼10^2^-10^4^ voxels. This may miss some large segments where small-to medium-sized activations only produce small fractional volumes. 5) We recognize that statistical methods are limited due to the choice of thresholding and pre-preprocessing, which is particularly difficult with MRI (33, 95–97). We minimized these potential issues by rigorously examining each processing step for image quality, by ensuring sample size was sufficient to detect moderate to large signal enhancements from Mn(II), and by selecting statistical thresholds based on effect size and false positive rate corrections. Fractional volumes, pioneered here, provide quantitative detail about the volumetric extent of Mn(II)-enhanced signals, and are limited by their dependence on statistical parametric mapping and segmentation, and could miss variability and discrete locations of signal intensities within segments. Caveats of MEMRI technology include pre-existing tissue enhancement, intensity normalization procedures, and differences in dosing and imaging parameters. We have dealt with each of these in detail in this study, as described above.

To what degree does Mn(II) signal faithfully represent neural activity? Staining by c-fos shown here and previous reports, including IEG and non-IEG methodology, contend that Mn(II) tags active neurons (32, 36–41, 98, 99). Patterns of activity in response to challenge as seen by MEMRI reproduce those detected by IEG expression (32, 98). Both MEMRI and c-fos are dependent on calcium entry into neurons, though minor non-voltage-gated calcium channel dependent mechanisms may exist for MEMRI (99, 100). MEMRI acts as a direct, quantitative reporter of electrochemical flux and neural depolarization (40). Axonal transport from an active segment to another inactive segment cannot explain the timing and pattern of accumulations across the brain. Therefore, we believe that Mn(II)-enhanced signals are due to activity and not secondary to transport or non-voltage dependent uptake. Spatiotemporal resolution of MEMRI is less granular than some other methods. At 100 µm^3^ voxels, our MEMRI has higher spatial resolution than current BOLD imaging but still not at the level of single neurons studied with electrophysiology. MEMRI’s temporal resolution is less than electrophysiology or BOLD imaging. However, MEMRI’s capacity for repeated longitudinal brain-wide imaging across a long time span of neural activity in awake, freely moving mice makes these tradeoffs worthwhile. Thus, Mn(II)-enhanced MR signal intensities and statistical activation volumes are quantitative readouts of localized neural activity across the brain. MEMRI pinpoints regions and conditions worthy of study at greater resolutions by other techniques.

ELA is a naturalistic translationally relevant paradigm that provides an experimental platform to test neuromodulatory interventions for human adverse childhood experiences (ACEs) and the impacts of subsequent life-threatening events. That brain-wide destabilization exists in adult mice after ELA provides insight into neurobiological observations of mental disorders arising after ACEs in humans (101, 102), where similar longitudinal studies are not feasible. Regions highlighted as being particularly affected, such as locus coeruleus, raphe nuclei, and hypothalamic nuclei, suggest plausible neuro-chemical or –physiological targets.

In conclusion, ELA has a brain-wide impact on balance of neural activity among widely distributed sub-regions and their dynamics across a series of experiences, from freely moving exploration in the home cage to immediate– and post-threat conditions. We identify specific subcomponents of this brain-wide network that are differentially activated depending upon condition in adult mice who experienced ELA and compare these with the standard, commonly studied, adult mouse. Future studies dissecting the functional connections suggested by these MEMRI results will provide valuable information on coordination of coactivated regions. A next step will be to determine the functional significance of these disparate segmental activities, how they interact in a serially dependent manner to reconfigure future brain states, and whether these interactions contribute to vulnerabilities for mental health disorders and/or compulsive drug use. Ultimately this longitudinal MEMRI technique will provide insights into neurophysiological/biochemical mechanisms contributing to the reconfiguration of brain states by ELA and inspire reparative/restorative interventions to protect from the harm of ELA and secondary traumatic events later in adulthood.

## Materials and Methods

### Mice

Animal protocols were approved by Institutional Animal Care and Use Committees at both California Institute of Technology (Caltech) and University of New Mexico. Numbers were determined by statistical power analyses. Both Std and ELA reared groups of C57BL/6J mice (Stock No: 000664) were bred at Jackson Labs (JAX). Adults from each group went through parallel acute threat timelines, were handled by the same experimentalists, and imaged at Caltech (n = 24, 12 in each group), in a series of experiments performed over a period of 3.8 years. For ELA, we followed a fragmented care paradigm (11, 12). Longitudinal MEMRI commenced at ages 11-13 wk. This age was chosen because brain development is considered largely complete by 3 months (103). Equal numbers of male (n = 6) and female (n = 6) ELA pups were used from 5 weaning groups. Twelve male Std mice selected from 5 different weaning cages were subjected to the longitudinal experimental protocol. No litter/cage effects were detected for either group.

### Video Recordings and Acute Threat Exposure

We followed our previously published longitudinal imaging protocol (32). Briefly, prior to imaging and during TMT exposure mice were tracked in a custom arena using Noldus Ethovision (XT15) (32, 104, 105). Prior to TMT exposure, mice were bright, alert and responsive, and displayed normal motility. Percentage of time spent in the light side of the arena and distance travelled were calculated and statistically evaluated with FDR correction in R v4.2.1 (*nlme* and *emmeans* packages) (106–111). Graphs were generated in R (*ggplot2*) (112).

### Longitudinal MEMRI

After capture of a non-enhanced image, mice were injected i.p. with 0.3 mmol/kg of MnCl_2_-4H_2_0 in sterile buffered saline and then imaged with contrast according to the timeline previously reported (32). Between scanning sessions, video recordings, or MnCl_2_ injections, mice were returned to their home cage. Mn(II) kinetics were followed by ROI analysis across the timeline to test for expected intensity patterns. For optimization of MEMRI imaging parameters, refer to our review (33).

### MR Image Pre-Processing

MR images from both groups were pre-processed together in batch according to standardized protocols (33, 50, 52). Briefly, raw MR image quality was visually assessed and SNR quantified. Image artifacts were corrected when needed and outlier images excluded from statistical analyses. Std and ELA pre– and post-Mn(II) images were then skull-stripped, intensity normalized and anatomically aligned together in batch. Each processing step was verified and validated (***SI Appendix* Fig. S1-2**).

### MR Image Analysis

*Comparison of Anatomy Between Groups.* For comparison of anatomy between groups, we calculated Jacobian Determinants of the Deformation fields generated by the affine-nonlinear alignment step. Average Jacobian determinant values were calculated using our atlas-based segmentation process (see below). Two sample t-tests between Std and ELA values were performed for each segment, FDR correction.

*Measurement and Analysis of MR Intensities in ROIs.* Measurements were made from pre– and post-Mn(II)-enhanced MR images that had been skull-stripped, scaled, and aligned. Coordinates (x/y/z) for regions of interest (3 x 3 x 3 voxels) were selected by projecting the annotated atlas labels onto the aligned grayscale template in FSLeyes and placing the crosshairs within segments that appeared different in SPM comparisons to pre-Mn(II) and that corresponded to previously reported TMT-responsive regions (32, 77). Intensities beyond 3 standard deviations from the sample average for each ROI were considered outliers and removed; none were found. Mean intensities from ROI measurements were compared by Student’s paired and two-sample t-tests, or by Welch’s test (if heteroscedastic), in R (*t.test)*. Cohen’s D effect sizes and FDR adjustments (*p.adjust)* were calculated. Cohen’s D for heteroscedastic data were calculated using Welch’s non-pooled standard deviation.

*Statistical Parametric Mapping (SPM).* For SPM, all images were smoothed to meet SPM assumptions with a 0.15 mm^3^ Gaussian kernel, the minimum size allowable (57, 94). We adjusted for small baseline differences between groups in their pre-Mn(II) images by subtracting the average voxel-wise differences of intensities from each of the ELA images. After this adjustment, an unpaired t-test of pre-Mn(II) images yielded no voxels with significant differences between groups. Voxel-wise T-maps of statistically significant differences between images of different conditions within group (paired t-test) or between groups (unpaired t-test) were calculated using SPM12 (94). Statistical maps were cluster corrected (*cluster corr.*) in SPM12 with a minimum cluster size of 8 voxels (94). Fiber tracts and ventricles segments were masked from statistical maps using our *InVivo* Atlas. Setting statistical thresholds included consideration of minimal detectable effects and smallest cluster size needed to remove false positives while minimizing loss of signal.

*Atlas Alignment, Segmentation, and Calculation of Fractional Volumes.* Our updated *InVivo* Atlas grayscale image was inversely aligned to the dataset, and the warp field applied to the label image. This permitted automated segmentation of the images into 117 regions, including 102 neuronal clusters a.k.a. nuclei (32). Segment masks were created and applied using *fslmaths*. The number of statistically enhanced voxels were identified as those from SPM paired or unpaired t-tests exceeding thresholds set according to our criteria described above. The number of total voxels within each segment were counted using *fslstats* (24). The fraction of enhanced voxels from SPM paired (FAV) or unpaired (FDV) t-tests to total voxels within each segment were calculated in R and visualized as column graphs and line plots using *ggplot2* (111–113). Segments were categorized into patterns of FDV dynamics by the criteria of FDV > 0.1 for at least one condition and |ΔFDV| > 0.1 between conditions.

### Immunohistochemistry, c-fos Quantification and Correlation to MEMRI

Brains from mice, euthanized at the conclusion of MR imaging (5 Std and 6 ELA), were prepared for histology according to our protocol (32, 114). We selected 1-3 sections also identifiable in MR slices from each mouse that had regions with both very high and very low numbers of c-fos stained nuclei. Nuclei were counted in a 3 x 3 grid of 100 μm^2^ squares from 1-3 x 35 μm thick sections of each brain, and MR intensities (SI) from each condition measured from 100 µm^3^ voxels in corresponding slices. Counts ranged from 0 to 27. MEMRI SI ranged from ∼6,900-10,100. Counts and SI were visualized in quartile boxplots with R (*ggplot2*). A square root transformation linearized c-fos+ count data for determination of effect size and statistical significance. Regions counted in the same section or measured in the same slice were compared statistically by linear mixed models and by post-hoc paired t-tests with FDR correction. Regions were compared between groups for both modalities by linear mixed models and post-hoc two-sample t-tests with FDR correction. Counts and SI of each group were averaged by region and individual and then correlated, with p-value determined by permutation testing.

Please see ***SI Appendix* Extended Methods** for additional methodological details.

## Acknowledgments

We gratefully acknowledge support from the Biological Imaging Facility/Center and the Beckman Institute at Caltech where imaging was performed, and funding from NIH R01MH096093 (ELB), NIH NIA P20AG068077 (ELB), NIH NINDS F99NS139535 (TWU), Zilkha Neurogenetic Institute (REJ) and Harvey Family Endowment (ELB). We appreciate staff, faculty and trainees at UNM, USC and Caltech including Christopher S. Medina, Daniel Barto, Xiaowei Zhang and Sharon Wu Lin (see ***SI Appendix***). We thank Chiara Padilla for proofreading support. We are grateful to Harry B. Gray, Arnold O. Beckman Professor of Chemistry, at Caltech for insights on Mn(II) metallobiology and advice on the manuscript. See ***SI Appendix* Additional Acknowledgements**.

## Author Contributions

T.W.U. designed and performed research, contributed analytical tools, analyzed data and wrote the manuscript; R.E.J. designed and performed research, developed the imaging protocol, analyzed data and edited the manuscript; E.L.B. designed and performed research, analyzed data, wrote and edited the manuscript, and raised funding for the project.

## Classification

Biological Sciences (Major), Neuroscience (Minor).

## References

1. Felitti VJ, et al. (1998) Relationship of Childhood Abuse and Household Dysfunction to Many of the Leading Causes of Death in Adults. American Journal of Preventive Medicine 14(4):245–258.

2. Merrick M, et al. (2019) Vital Signs: Estimated Proportion of Adult Health Problems Attributable to Adverse Childhood Experiences and Implications for Prevention — 25 States, 2015–2017. MMWR. Morbidity and Mortality Weekly Report 68.

3. Zarse EM, et al. (2019) The adverse childhood experiences questionnaire: Two decades of research on childhood trauma as a primary cause of adult mental illness, addiction, and medical diseases. Cogent Medicine 6(1):1581447.

4. Spadoni AD, et al. (2022) Contribution of early-life unpredictability to neuropsychiatric symptom patterns in adulthood. Depression and Anxiety 39(10-11):706–717.

5. Yehuda R, et al. (2010) Putative biological mechanisms for the association between early life adversity and the subsequent development of PTSD. Psychopharmacology 212(3):405–417.

6. Chen Y & Baram TZ (2016) Toward Understanding How Early-Life Stress Reprograms Cognitive and Emotional Brain Networks. Neuropsychopharmacology 41(1):197–206.

7. Kraaijenvanger EJ, et al. (2020) Impact of early life adversities on human brain functioning: A coordinate-based meta-analysis. Neuroscience & Biobehavioral Reviews 113:62–76.

8. Malave L, Van Dijk MT, & Anacker C (2022) Early life adversity shapes neural circuit function during sensitive postnatal developmental periods. Translational Psychiatry 12(1):306.

9. Garcia-Arocena D (2015) More researchers are using B6J mice than ever before. (The Jackson Laboratory).

10. Laboratory TJ (2024) Mouse Room Conditions.

11. Molet J, Maras PM, Avishai-Eliner S, & Baram TZ (2014) Naturalistic rodent models of chronic early-life stress. Developmental Psychobiology 56(8):1675–1688.

12. Rice CJ, Sandman CA, Lenjavi MR, & Baram TZ (2008) A Novel Mouse Model for Acute and Long-Lasting Consequences of Early Life Stress. Endocrinology 149(10):4892–4900.

13. Bercum FM, Navarro Gomez MJ, & Saddoris MP (2021) Elevated fear responses to threatening cues in rats with early life stress is associated with greater excitability and loss of gamma oscillations in ventral-medial prefrontal cortex. Neurobiol Learn Mem 185:107541.

14. Bolton JL, et al. (2018) Anhedonia Following Early-Life Adversity Involves Aberrant Interaction of Reward and Anxiety Circuits and Is Reversed by Partial Silencing of Amygdala Corticotropin-Releasing Hormone Gene. Biological Psychiatry 83(2):137–147.

15. Bolton JL, et al. (2022) Early stress-induced impaired microglial pruning of excitatory synapses on immature CRH-expressing neurons provokes aberrant adult stress responses. Cell Rep 38(13):110600.

16. Brosens N, Simon C, Kessels HW, Lucassen PJ, & Krugers HJ (2023) Early life stress lastingly alters the function and AMPA-receptor composition of glutamatergic synapses in the hippocampus of male mice. J Neuroendocrinol 35(12):e13346.

17. de Carvalho G, Khoja S, Haile MT, & Chen LY (2023) Early life adversity impaired dorsal striatal synaptic transmission and behavioral adaptability to appropriate action selection in a sex-dependent manner. Front Synaptic Neurosci 15:1128640.

18. Guadagno A, et al. (2018) Reduced resting-state functional connectivity of the basolateral amygdala to the medial prefrontal cortex in preweaning rats exposed to chronic early-life stress. Brain Struct Funct 223(8):3711–3729.

19. Guadagno A, Verlezza S, Long H, Wong TP, & Walker CD (2020) It Is All in the Right Amygdala: Increased Synaptic Plasticity and Perineuronal Nets in Male, But Not Female, Juvenile Rat Pups after Exposure to Early-Life Stress. The Journal of neuroscience: the official journal of the Society for Neuroscience 40(43):8276–8291.

20. Guadagno A, Wong TP, & Walker CD (2018) Morphological and functional changes in the preweaning basolateral amygdala induced by early chronic stress associate with anxiety and fear behavior in adult male, but not female rats. Prog Neuropsychopharmacol Biol Psychiatry 81:25–37.

21. Gunn BG, et al. (2013) Dysfunctional astrocytic and synaptic regulation of hypothalamic glutamatergic transmission in a mouse model of early-life adversity: relevance to neurosteroids and programming of the stress response. The Journal of neuroscience: the official journal of the Society for Neuroscience 33(50):19534–19554.

22. Johnson FK, et al. (2018) Amygdala hyper-connectivity in a mouse model of unpredictable early life stress. Transl Psychiatry 8(1):49.

23. Kooiker CL, Chen Y, Birnie MT, & Baram TZ (2023) Genetic Tagging Uncovers a Robust, Selective Activation of the Thalamic Paraventricular Nucleus by Adverse Experiences Early in Life. Biol Psychiatry Glob Open Sci 3(4):746–755.

24. Levis SC, et al. (2022) Enduring disruption of reward and stress circuit activities by early-life adversity in male rats. Transl Psychiatry 12(1):251.

25. Malter Cohen M, et al. (2013) Early-life stress has persistent effects on amygdala function and development in mice and humans. Proc Natl Acad Sci U S A 110(45):18274–18278.

26. Raineki C, Cortes MR, Belnoue L, & Sullivan RM (2012) Effects of early-life abuse differ across development: infant social behavior deficits are followed by adolescent depressive-like behaviors mediated by the amygdala. The Journal of neuroscience: the official journal of the Society for Neuroscience 32(22):7758–7765.

27. Yan CG, et al. (2017) Aberrant development of intrinsic brain activity in a rat model of caregiver maltreatment of offspring. Transl Psychiatry 7(1):e1005.

28. Holschneider DP, Guo Y, Mayer EA, & Wang Z (2016) Early life stress elicits visceral hyperalgesia and functional reorganization of pain circuits in adult rats. Neurobiol Stress 3:8–22.

29. Islam R, et al. (2024) Early adversity causes sex-specific deficits in perforant pathway connectivity and contextual memory in adolescent mice. Biol Sex Differ 15(1):39.

30. White JD, et al. (2020) Early life stress causes sex-specific changes in adult fronto-limbic connectivity that differentially drive learning. Elife 9.

31. Kooiker CL, et al. (2024) Paraventricular Thalamus Neuronal Ensembles Encode Early-life Adversity and Mediate the Consequent Sex-dependent Disruptions of Adult Reward Behaviors. (Cold Spring Harbor Laboratory).

32. Uselman TW, Barto DR, Jacobs RE, & Bearer EL (2020) Evolution of brain-wide activity in the awake behaving mouse after acute fear by longitudinal manganese-enhanced MRI. Neuroimage 222:116975.

33. Uselman TW, Medina CS, Gray HB, Jacobs RE, & Bearer EL (2022) Longitudinal manganese-enhanced magnetic resonance imaging of neural projections and activity. e4675.

34. Vousden DA, et al. (2018) Continuous manganese delivery via osmotic pumps for manganese-enhanced mouse MRI does not impair spatial learning but leads to skin ulceration. NeuroImage 173:411–420.

35. Aoki I, Wu YJ, Silva AC, Lynch RM, & Koretsky AP (2004) In vivo detection of neuroarchitecture in the rodent brain using manganese-enhanced MRI. Neuroimage 22(3):1046–1059.

36. Bedenk BT, et al. (2018) Mn(2+) dynamics in manganese-enhanced MRI (MEMRI): Cav1.2 channel-mediated uptake and preferential accumulation in projection terminals. Neuroimage 169:374–382.

37. Narita K, Kawasaki F, & Kita H (1990) Mn and Mg influxes through Ca channels of motor nerve terminals are prevented by verapamil in frogs. Brain research 510(2):289–295.

38. Hattori S, et al. (2013) In vivo evaluation of cellular activity in alphaCaMKII heterozygous knockout mice using manganese-enhanced magnetic resonance imaging (MEMRI). Front Integr Neurosci 7:76.

39. Lin YJ & Koretsky AP (1997) Manganese ion enhances T1-weighted MRI during brain activation: an approach to direct imaging of brain function. Magnetic resonance in medicine 38(3):378–388.

40. Svehla P, Bedecarrats A, Jahn C, Nargeot R, & Ciobanu L (2018) Intracellular manganese enhanced MRI signals reflect the frequency of action potentials in Aplysia neurons. J Neurosci Methods 295:121–128.

41. Malkova NV, Gallagher JJ, Yu CZ, Jacobs RE, & Patterson PH (2014) Manganese-enhanced magnetic resonance imaging reveals increased DOI-induced brain activity in a mouse model of schizophrenia. Proc Natl Acad Sci U S A 111(24):E2492–2500.

42. Yu X, Sanes DH, Aristizabal O, Wadghiri YZ, & Turnbull DH (2007) Large-scale reorganization of the tonotopic map in mouse auditory midbrain revealed by MRI. Proceedings of the National Academy of Sciences 104(29):12193–12198.

43. Yu X, Wadghiri YZ, Sanes DH, & Turnbull DH (2005) In vivo auditory brain mapping in mice with Mn-enhanced MRI. Nature Neuroscience 8(7):961–968.

44. Bearer EL, et al. (2007) Role of neuronal activity and kinesin on tract tracing by manganese-enhanced MRI (MEMRI). Neuroimage 37 Suppl 1:S37–46.

45. Bearer EL, Zhang X, & Jacobs RE (2007) Live imaging of neuronal connections by magnetic resonance: Robust transport in the hippocampal–septal memory circuit in a mouse model of Down syndrome. NeuroImage 37(1):230–242.

46. Bearer EL, Zhang X, Janvelyan D, Boulat B, & Jacobs RE (2009) Reward circuitry is perturbed in the absence of the serotonin transporter. Neuroimage 46(4):1091–1104.

47. Zhang X, et al. (2010) Altered neurocircuitry in the dopamine transporter knockout mouse brain. PLoS One 5(7):e11506.

48. Gallagher JJ, Zhang X, Ziomek GJ, Jacobs RE, & Bearer EL (2012) Deficits in axonal transport in hippocampal-based circuitry and the visual pathway in APP knock-out animals witnessed by manganese enhanced MRI. Neuroimage 60(3):1856–1866.

49. Gallagher JJ, et al. (2013) Altered reward circuitry in the norepinephrine transporter knockout mouse. PLoS One 8(3):e57597.

50. Delora A, et al. (2016) A simple rapid process for semi-automated brain extraction from magnetic resonance images of the whole mouse head. J Neurosci Methods 257:185–193.

51. Medina CS, et al. (2017) Hippocampal to basal forebrain transport of Mn(2+) is impaired by deletion of KLC1, a subunit of the conventional kinesin microtubule-based motor. Neuroimage 145(Pt A):44–57.

52. Medina CS, Manifold-Wheeler B, Gonzales A, & Bearer EL (2017) Automated Computational Processing of 3-D MR Images of Mouse Brain for Phenotyping of Living Animals. Curr Protoc Mol Biol 119:29A 25 21-29A 25 38.

53. Bearer EL, et al. (2018) Alterations of functional circuitry in aging brain and the impact of mutated APP expression. Neurobiol Aging 70:276–290.

54. Bearer EL, Barto D, & Jacobs RE (2019) Imaging the evolution acute fear: Longitudinal whole brain imaging in living mice of neural activity with MEMRI. Proceedings of the International Society for Magnetic Resonance in Medicine… Scientific Meeting and Exhibition. International Society for Magnetic Resonance in Medicine. Scientific Meeting and Exhibition, (NIH Public Access).

55. Medina CS, et al. (2019) Decoupling the Effects of the Amyloid Precursor Protein From Amyloid-beta Plaques on Axonal Transport Dynamics in the Living Brain. Front Cell Neurosci 13:501.

56. Phinyomark A, Ibanez-Marcelo E, & Petri G (2017) Resting-State fMRI Functional Connectivity: Big Data Preprocessing Pipelines and Topological Data Analysis. IEEE Transactions on Big Data 3(4):415–428.

57. Flandin G & Friston K (2008) Statistical parametric mapping (SPM). Scholarpedia 3(4):6232.

58. Penny WD, Friston KJ, Ashburner JT, Kiebel SJ, & Nichols TE (2011) Statistical Parametric Mapping: The Analysis of Functional Brain Images (Elsevier Science).

59. Pallast N, et al. (2019) Processing Pipeline for Atlas-Based Imaging Data Analysis of Structural and Functional Mouse Brain MRI (AIDAmri). Frontiers in Neuroinformatics 13.

60. Fendt M, Endres T, Lowry CA, Apfelbach R, & McGregor IS (2005) TMT-induced autonomic and behavioral changes and the neural basis of its processing. Neuroscience and biobehavioral reviews 29(8):1145–1156.

61. Rosen J, Asok A, & Chakraborty T (2015) The smell of fear: innate threat of 2,5-dihydro-2,4,5-trimethylthiazoline, a single molecule component of a predator odor. 9.

62. Gallo M, et al. (2019) Limited Bedding and Nesting Induces Maternal Behavior Resembling Both Hypervigilance and Abuse. 13.

63. Matsukawa M, Yoshikawa M, Katsuyama N, Aizawa S, & Sato T (2022) The Anterior Piriform Cortex and Predator Odor Responses: Modulation by Inhibitory Circuits. Front Behav Neurosci 16:896525.

64. Matsumoto H, et al. (2010) Spatial Arrangement of Glomerular Molecular-Feature Clusters in the Odorant-Receptor Class Domains of the Mouse Olfactory Bulb. 103(6):3490-3500.

65. Cohen J (2013) Statistical Power Analysis for the Behavioral Sciences.

66. Kimbrough A, et al. (2020) Brain-wide functional architecture remodeling by alcohol dependence and abstinence. 117(4):2149-2159.

67. Motta SC, et al. (2009) Dissecting the brain’s fear system reveals the hypothalamus is critical for responding in subordinate conspecific intruders. Proceedings of the National Academy of Sciences 106(12):4870–4875.

68. Greene AS, Horien C, Barson D, Scheinost D, & Constable RT (2023) Why is everyone talking about brain state? Trends Neurosci 46(7):508–524.

69. Ben-Shaul Y, Katz LC, Mooney R, & Dulac C (2010) In vivo vomeronasal stimulation reveals sensory encoding of conspecific and allospecific cues by the mouse accessory olfactory bulb. 107(11):5172–5177.

70. Canteras NS (2002) The medial hypothalamic defensive system: Hodological organization and functional implications. Pharmacology Biochemistry and Behavior 71(3):481–491.

71. Canteras NS, Chiavegatto S, Ribeiro do Valle LE, & Swanson LW (1997) Severe Reduction of Rat Defensive Behavior to a Predator by Discrete Hypothalamic Chemical Lesions. Brain Research Bulletin 44(3):297–305.

72. Janitzky K, D׳Hanis W, Kröber A, & Schwegler H (2015) TMT predator odor activated neural circuit in C57BL/6J mice indicates TMT-stress as a suitable model for uncontrollable intense stress. Brain research 1599:1–8.

73. Kunwar PS, et al. (2015) Ventromedial hypothalamic neurons control a defensive emotion state. eLife 4:e06633.

74. Pérez-Gómez A, et al. (2015) Innate Predator Odor Aversion Driven by Parallel Olfactory Subsystems that Converge in the Ventromedial Hypothalamus. Current Biology 25(10):1340–1346.

75. Koob GF & Volkow ND (2016) Neurobiology of addiction: a neurocircuitry analysis. The lancet. Psychiatry 3(8):760–773.

76. Russo SJ & Nestler EJ (2013) The brain reward circuitry in mood disorders. Nature Reviews Neuroscience 14(9):609–625.

77. Silva BA, Gross CT, & Graff J (2016) The neural circuits of innate fear: detection, integration, action, and memorization. Learn Mem 23(10):544–555.

78. Coletta L, et al. (2020) Network structure of the mouse brain connectome with voxel resolution. 6(51):eabb7187.

79. Salvan P, et al. (2023) Serotonin regulation of behavior via large-scale neuromodulation of serotonin receptor networks. Nature Neuroscience 26(1):53–63.

80. Taylor NL, et al. (2022) Structural connections between the noradrenergic and cholinergic system shape the dynamics of functional brain networks. NeuroImage 260:119455.

81. Zerbi V, et al. (2019) Rapid Reconfiguration of the Functional Connectome after Chemogenetic Locus Coeruleus Activation. Neuron 103(4):702–718.e705.

82. David & Adolphs R (2014) A Framework for Studying Emotions across Species. Cell 157(1):187–200.

83. Flavell SW, Gogolla N, Lovett-Barron M, & Zelikowsky M (2022) The emergence and influence of internal states. Neuron 110(16):2545–2570.

84. Jaric I, et al. (2022) The rearing environment persistently modulates mouse phenotypes from the molecular to the behavioural level. PLoS Biol 20(10):e3001837.

85. Jaric I, et al. (2024) Using mice from different breeding sites fails to improve replicability of results from single-laboratory studies. Lab Anim (NY) 53(1):18–22.

86. Usui T, Macleod MR, McCann SK, Senior AM, & Nakagawa S (2021) Meta-analysis of variation suggests that embracing variability improves both replicability and generalizability in preclinical research. PLoS Biol 19(5):e3001009.

87. Chang C-H & Gean P-W (2019) The Ventral Hippocampus Controls Stress-Provoked Impulsive Aggression through the Ventromedial Hypothalamus in Post-Weaning Social Isolation Mice. Cell Reports 28(5):1195–1205.e1193.

88. Manzano Nieves G, Bravo M, & Bath KG (2023) Early life adversity ablates sex differences in active versus passive threat responding in mice. Stress:1–32.

89. Spring S, Lerch JP, & Henkelman RM (2007) Sexual dimorphism revealed in the structure of the mouse brain using three-dimensional magnetic resonance imaging. NeuroImage 35(4):1424–1433.

90. Bruce MR, et al. (2021) Sexually dimorphic neuroanatomical differences relate to ASD-relevant behavioral outcomes in a maternal autoantibody mouse model. Molecular Psychiatry 26(12):7530–7537.

91. Holz NE, et al. (2023) A stable and replicable neural signature of lifespan adversity in the adult brain. Nature Neuroscience.

92. Lerch JP, et al. (2011) Maze training in mice induces MRI-detectable brain shape changes specific to the type of learning. NeuroImage 54(3):2086–2095.

93. Ashburner J, et al. (2012) SPM8 manual.

94. Ashburner J, et al. (2014) SPM12 manual.

95. Carter CS, Lesh TA, & Barch DM (2016) Thresholds, Power, and Sample Sizes in Clinical Neuroimaging. Biological Psychiatry: Cognitive Neuroscience and Neuroimaging 1(2):99–100.

96. Mumford JA (2012) A power calculation guide for fMRI studies. Social Cognitive and Affective Neuroscience 7(6):738–742.

97. Vandekar SN & Stephens J (2021) Improving the replicability of neuroimaging findings by thresholding effect sizes instead of p-values. 42(8):2393–2398.

98. Lehallier B, et al. (2012) Brain Processing of Biologically Relevant Odors in the Awake Rat, as Revealed by Manganese-Enhanced MRI. PLoS ONE 7(10):e48491.

99. Petrus E, Saar G, Daoust A, Dodd S, & Koretsky AP (2021) A hierarchy of manganese competition and entry in organotypic hippocampal slice cultures. NMR in Biomedicine 34(4).

100. Bartelle BB, Szulc KU, Suero-Abreu GA, Rodriguez JJ, & Turnbull DH (2013) Divalent metal transporter, DMT1: A novel MRI reporter protein. Magnetic resonance in medicine 70(3):842–850.

101. Duffy KA, McLaughlin KA, & Green PA (2018) Early life adversity and health-risk behaviors: proposed psychological and neural mechanisms. 1428(1):151–169.

102. Hakamata Y, Suzuki Y, Kobashikawa H, & Hori H (2022) Neurobiology of early life adversity: A systematic review of meta-analyses towards an integrative account of its neurobiological trajectories to mental disorders. Frontiers in Neuroendocrinology 65:100994.

103. Hammelrath L, et al. (2016) Morphological maturation of the mouse brain: An in vivo MRI and histology investigation. NeuroImage 125:144–152.

104. Brigman JL, Mathur P, Lu L, Williams RW, & Holmes A (2009) Genetic relationship between anxiety-related and fear-related behaviors in BXD recombinant inbred mice. 20(2):204–209.

105. Noldus LPJJ, Spink AJ, & Tegelenbosch RAJ (2001) EthoVision: A versatile video tracking system for automation of behavioral experiments. Behavior Research Methods, Instruments, & Computers 33(3):398–414.

106. Fox J & Weisberg S (2019) An R Companion to Applied Regression.

107. Lenth RV (2022) emmeans: Estimated Marginal Means, aka Least-Squares Means.

108. Pinheiro J, Bates D, & Team RC (2022) nlme: Linear and Nonlinear Mixed Effects Models.

109. Pinheiro JC & Bates DM (2000) Mixed-Effects Models in S and S-PLUS.

110. Posit t (2022) RStudio: Integrated Development Environment for R.

111. Team RC (2022) R: A Language and Environment for Statistical Computing.

112. Wickham H (2016) ggplot2: Elegant Graphics for Data Analysis.

113. Wickham H, et al. (2019) Welcome to the Tidyverse. 4(43):1686.

114. Tyszka JM, Readhead C, Bearer EL, Pautler RG, & Jacobs RE (2006) Statistical diffusion tensor histology reveals regional dysmyelination effects in the shiverer mouse mutant. NeuroImage 29(4):1058–1065.

