## Supplemental Information Appendix for "Reconfiguration of brain-wide neural activity after early life adversity"

PNAS

### Supporting Information for
Reconfiguration of brain-wide neural activity after early life adversity.

Taylor W. Uselman[1], Russell E. Jacobs[2,3], Elaine L Bearer[1,3]

[1] University of New Mexico Health Sciences Center, Albuquerque, NM 87131
[2] Zilkha Neurogenetic Institute, Keck School of Medicine of University of Southern California, Los Angeles, CA 90033
[3] California Institute of Technology, Pasadena, CA 91125

**Corresponding Author:** Elaine L. Bearer

**This PDF file includes:**

Table of Contents
Supporting Text
- A. Additional Acknowledgements
- B. Abbreviations
- C. Extended Methods

SI Figures S1 to S11
SI Tables S1 to S7
Legend for Movie S1
SI References

**Other supporting materials for this manuscript include the following:**

Movie S1

**Table of Contents**:

**Supporting Text**

**A. Additional Acknowledgements**

**B. Abbreviations**

**C. Extended Methods**

**1. Mice.**

**2. Video Recordings and Acute Threat Exposure.**

*2.1. Arena and Predator Odor.*

*2.2. Movement Tracking from Video Recordings.*

**3. Longitudinal MEMRI.**

*3.1. MRI Hardware and Scan Parameters.*

*3.2. Timing.*

*3.3. Mn(II) Accumulation Kinetics.*

**4. MR Image Pre-Processing and Quality Assurance.**

*4.1. Image Pre-Processing Pipeline Overview.*

*4.2. Review of Raw MR Images and Slice Interpolation.*

*4.3. Measurements of Signal-to-Noise Ratio (SNR).*

*4.4. Quality Control of Alignments and Jaccard Similarity.*

*4.5. Modal Scaling Evaluation.*

**5. MR Image Analysis.**

*5.1. Comparison of Anatomy Between Groups, Jacobian Determinant of Deformation Fields.*

*5.2. Statistical Parametric Mapping (SPM): Selection of Statistical Thresholds.*

*5.3. SPM: Adjustment of Post-Mn(II) Images for Average Differences Between Groups' Pre-Mn(II) Images.*

*5.4. InVivo Atlas: Alignment and Automated Segmentation*

*5.5. Formulas Used to Calculate Fractional Volumes.*

*5.6. Evaluation of Sex Differences.*

*5.7. Diagrams of the Balance of Brain Wide Patterns of Activity.*

**6. Immunohistochemistry, c-fos Quantification and Correlation to MEMRI.**

*6.1. Immunohistochemical Staining for c-fos and Microscopy.*

*6.2. Quantification and Correlation c-fos+ counts and MEMRI SI.*

**Figures S1 to S11**

**Fig. S1.** Flow Diagram of MEMRI Image Processing and Analysis Pipeline.

**Fig. S2.** MRI Image Processing Quality Controls and Mn(II)-enhanced Signal Kinetics.

**Fig. S3.** Acute and Longer-term Transitions of ELA Brain States as they Progress across Conditions

**Fig. S4.** Test-retest Reliability in MEMRI Intensities between Subsets of Std and ELA mice.

**Fig. S5.** Correlations between MEMRI SI and c-fos Counts.

**Fig. S6.** Comparison of ROI Intensities before and after Smoothing at a Kernel Size of 0.15 $mm^3$.

**Fig. S7.** Direct Comparisons between Successive Conditions in ELA and in Std.

**Fig. S8.** Comparison of Baseline pre-Mn(II) Images between Groups and Effects of Adjustment on post-Mn(II) Intensity Maps.

**Fig. S9.** Absence of Anatomical Differences between Adult ELA and Std Mice.

**Fig. S10.** Female Responses Differ Slightly from Male at each Condition.

**Fig. S11.** Summary Diagrams of Anatomical Distributions of Neural Activity Patterns Characterizing Brain-States of Both Groups at each of the Three Conditions.

**Tables S1 to S7**

**Table S1.** MRI Hardware and Scan Parameters for ELA and Std Groups

**Table S2.** Voxel Coordinates for Regions of Interest (ROI) Analysis

**Table S3.** Degree and Statistics of SI in Olfactory and Amygdala ROIS of ELA.

**Table S4.** Degree and Statistics of SI of Both Groups in ROIs Responsive to TMT in Standard.

**Table S5.** Key to *InVivo* Atlas (v10.4) Segmental Abbreviations in Anatomical Order, color-coded per domain, as in the main text figures.

**Table S6A.** Segment-wise fractional activation volumes (FAV) in Standard and ELA at each condition.

**Table S6B.** Segment-wise fractional volumes from comparisons between post-Mn(II) images in Standard and ELA mice

**Table S7.** Segment-wise fractional difference volumes (FDV) between groups at each condition, as in Fig. 6 of the main text.

**Legend for Movie S1**

**SI References**

#### Supporting Text

##### A. Additional Acknowledgements.

We appreciate Kathleen Kilpatrick for technical support and histology; Kevin P. Reagan and Daniel Perez-Rodriguez for laboratory support, and James Chavez for IT services at UNM. We thank Professors Jonathan Brigman, Erik B. Erhard, Natalie Adolphi and Kiran Bhaskar for helpful discussions of procedures, data analysis, and interpretations. We thank Evelyn Lockhart for guidance on scientific illustration and visual communication. We are grateful to Joe Gallagher for MRI scanning, video recording and animal handling at Caltech. We are indebted to Scott E. Frasier for the creation and direction of the Biological Imaging Center at Caltech.

##### B. Abbreviations for terminology in Main Text and SI Appendix.

For abbreviations used for anatomical domains and segments in the *InVivo* Atlas, please see ***SI Appendix*** **Table S5** below.

**MEMRI** = Manganese-enhanced magnetic resonance imaging
**Mn(II)** = Manganese II
**ELA** = Early life adversity / Early life adversity-reared mice
**Std** = Standard / Standard mice
**BL** = Baseline condition – behavior followed by imaging prior to Mn(II)
**HC** = Home cage condition – Mn(II) accumulation at 22-23 h after the first injection when the mouse had been in the home care environment
**TMT** = Acute threat condition - predator odor 2,3,5-Trimethyl-3-thiazoline
**D9** = Day 9 post-threat condition
**JAX** = Jackson laboratories
**FDR** = False discovery rate
**SNR** = Signal-to-noise ratio
**SPM** = Statistical parametric mapping
**ROI** = Region of interest
**SI** = Signal intensity
**FAV** = Fractional activation volume
**FDV** = Fractional difference volume
**Caltech** = California Institute of Technology
**UNM** = University of New Mexico
**P** (as in P2, P9, P10) = Post-natal days 2, 9 or 10.
**RF** = Radio frequency
**FLASH** = Fast Low Angle Shot

**TR** = Repetition time
**TE$_{eff}$** = Effective echo time
**FOV** = Field of view
**JSI** = Jaccard similarity index
**NMI** = Normalized mutual information
**NIfTI** = Neuroimaging Informatics Technology Initiative - MR image file type
**MDT** = Minimal Deformation Target
**JD** = Jacobian Determinant

**C. Extended Methods.**

**1. Mice.**

Numbers of mice were determined based on power analysis from MEMRI pilot data, assuming a Type I error rate of α = 0.05 and a Type II error rate of β = 0.2 (i.e., Power = 80%), and estimated a range of sample sizes of 8-12 mice per group based on intensities in regions expected to be enhanced (1, 2). A post-hoc sensitivity analysis of behavioral results demonstrated that this sample size also produced adequate effect sizes (d = 1.58-3.42). Pregnant dams bred at JAX were housed in Caltech's vivarium and monitored daily. For ELA, the number of pups per litter was limited to 5 after birth. On post-natal day P2, dams and pups were placed in a cage with mesh flooring and 1/3$^{rd}$ of standard bedding. Dams and pups remained undisturbed from P2 through P9. On P10, dams and pups were provided standard bedding. Twelve mice selected from each of 5 different litters were subjected to the longitudinal experimental protocol. For our comparison group (Std), we obtained adult C57BL/6J mice from JAX, the most phenotypically standardized mouse with this genotype. Dams and pups were group housed and pool-weaned to minimize any effects of rearing (3-5). Eight to ten week old Std mice from 5 different weaning cages were shipped to Caltech on 4 occasions over a period of 7 months. All mice were housed for at least 2 weeks in the Caltech vivarium before commencing experiments, regularizing their microbiota (6).

**2. Video Recordings and Acute Threat Exposure.**

*2.1. Arena and Predator Odor.* We assembled a custom two-compartment arena at UNM and shipped it to Caltech (1). Compartment 1: A black walled enclosure (38 × 13 × 20 cm) with a dark tinted ceiling that allowed infrared light to pass – the 'Dark' compartment. Compartment 2: A white-walled enclosure (38 × 28 × 20 cm) that had a translucent plastic ceiling that allowed illumination of the chamber – the 'Light' compartment. Mice could move between compartments through a floor level aperture (13 × 8 cm). The arena was maintained within a chemical fume hood and wiped down with 70% ethanol before and after every recording session to reduce odors. The arena contained a gauze pad in an open 100 mm petri dish onto which either saline or TMT (600 µl, 3% aq. solution)

(2,3,5-Trimethyl-3-thiazoline) was dripped (ConTech, Victoria, BC, Canada Cat #v300000368 lot 11125). Video recordings were performed at Caltech in the MR imaging suite.

*2.2. Movement Tracking from Video Recordings.* Ethovision XT15 was used to track the X-Y position of each mouse's center point over each 10-min recording at a sampling rate of 0.033 s. Frames missing the center point were determined manually or via linear interpolation of the adjacent video frames. Positional data for all animals were exported to text files, and subsequently compiled into a single data frame in R. For processing, the top and bottom 2.5% of data points were removed to limit the impact of outliers. Analysis was performed at UNM. Then positional tracks in both x- and y-directions were smoothed separately across 10 consecutive frames (~0.33 s) using kernel smoothing regression with the *ksmooth* function in the R stats package. To account for slight differences in arena positioning, positions were rotated as needed and min-max normalized in both x-y directions to match the physical dimensions of the arena (38 cm x 13 cm, Dark side; 38 cm x 28 cm, Light side). After the processing of positions, two variables were generated: 1) Distance Travelled and 2) Light-Dark. Distance traveled was calculated as the Euclidean distance between positions at adjacent video frames divided by the elapsed time for a single frame (~0.033 s). Cumulative distance travelled over 1-min epochs or over the full 10-min recording were also calculated for graphical and statistical comparisons. The percentage of time spent in light was determined by first calculating a binary 'Light-Dark' variable, where any X-position ≤ 13 cm was labeled as 'Dark', and any X-position > 13 cm was labeled as 'Light', corresponding to the dimensions of our arena. Then the percentage of time spent in the light side of the arena was calculated by dividing the frequency of video frames labeled as 'Light' over all video frames, 'Dark' or 'Light'. A mixed effects model for each group (Std and ELA) was constructed with fixed effects of Condition and Epoch and random effect of individual and rearing litter to assess for differences in distance travelled and percentage of time spent in the light. Post-hoc t-tests were then performed with FDR correction using the *emmeans and p.adjust* R packages: 1) within group, within epoch, between conditions; 2) within group, between conditions; and 3) between groups, within a condition. Cohen's D effect sizes were calculated using the *effect.size* R package (7).

**3. Longitudinal MEMRI.**

*3.1. MRI Hardware and Scan Parameters.* Images were collected at Caltech in an 11.7 T 89 mm vertical bore Bruker BioSpin Avance DRX500 scanner (Bruker BioSpin Inc, Billerica, MA) with a Micro2.5 gradient system and custom 35 mm linear birdcage radio frequency (RF) coil (1). We employed a $T_1$-weighted Fast Low Angle Shot (FLASH) imaging sequence (8, 9) with 8 averages, $TR/TE_{eff}$ = 25 ms/5 ms, and a 20° flip-angle. Voxel resolution was 100 µm isotropic with a matrix size of 200 x 124 x 82, FOV of 20.0 mm x 12.4 mm x 8.2 mm and a scan time of 34 min (1).

*3.2. Timeline.* This longitudinal imaging procedure is according to our previous publication (1). For each mouse, a pre-Mn(II) image was acquired, then the mouse was injected intraperitoneal (i.p.) with 0.3 mmol/kg of $MnCl_2$-$4H_20$ in buffered saline. Mice were then returned to their home cage for ~23-24 h before undergoing the first post-Mn(II), pre-TMT imaging. Mice were recovered from anesthesia for 30-40 min after the first contrast image and then exposed to TMT once fully recovered from anesthesia; after TMT mice were re-imaged at 0.5 h (ELA) and 1.5 h (Std). We found no difference in whole-brain SNR between groups for any imaging time point (***SI Appendix* Fig. S2**). After the post-TMT imaging session, mice were returned to their home cage. Eight days after the first $MnCl_2$-injection, mice were imaged again to capture residual or persistent Mn(II), then injected i.p. with a second dose of 0.3 mmol/kg of $MnCl_2$-$4H_20$ in sterile saline. Mice were returned to their home cage and then the D9 post-Mn(II) image acquired ~24 h later.

*3.3. Mn(II) Accumulation Kinetics.* To follow the kinetics of Mn(II) signal, we calculated mean intensities in three 5x5x5 voxel regions of interest (ROI) within OLF, HIP and thalamus (THA) of aligned and intensity normalized images at pre-Mn(II), HC post-Mn(II), and D9 pre-Mn(II) in each group (***SI Appendix*, Fig. S2G**). FSL functions *fslroi* and *fslstats* were used to measure ROI intensities and calculate the means, respectively. A plot of combined data was generated in R (*ggplot2*) (10). The kinetics of Mn(II)-dependent intensities matched previous reports (1, 11).

**4. MR Image Pre-Processing and Quality Assurance.**

*4.1. Image Pre-Processing Pipeline Overview.* MR images from both groups were pre-processed together in batch according to our protocols (2, 12, 13). Raw Bruker images were first converted to NIfTI format using ImageJ/FIJI and then visually checked for quality. Images were skull-stripped using our semi-automated pipeline (12), intensity normalized by modal scaling with our custom MATLAB software and anatomically aligned to the data set average using SPM. Each processing step was verified and validated (***SI Appendix* Fig. S1-2**). For comparison of anatomy between groups, we calculated Jacobian Determinants of the Deformation fields generated in the warp step (***SI Appendix* Fig. S8**) (14). For SPM t-tests, we first smoothed the images to ~0.3 $mm^3$ using a 0.15 $mm^3$ Gaussian kernel. Region of interest measurements were acquired from pre-smoothed images.

*4.2. Review of Raw MR Images and Slice Interpolation.* To assure quality of images during acquisition, we reviewed the original Bruker images by visualizing them in Fiji/ImageJ (15) and checking parameters using *Image > Type*, *Image > Show Info*, *Image > Properties*. (***SI Appendix* Fig. S2A and S2B**). In various images, small radiofrequency (RF) feed-through artifacts were identified in coronal slices. We replaced these slices with linear interpolation/averaging of adjacent slices, a common procedure (***SI Appendix* Fig. S2C**). We could not remove local signal artifacts in two images (1 pre-Mn(II) image and 1 post-Mn(II) image at HC) from Standard mouse #10 and removed these images from statistical analyses. Note that these images did not stand out as outliers in terms of global measures presented in ***SI Appendix*** **Fig. S1-2**, indicating the need for detailed review of images for quality control as described here. For summary of processing steps, quality controls, and statistical comparisons see ***SI Appendix*** **Fig. S1**.

*4.3. Measurements of Signal-to-Noise Ratio (SNR).* To assure Mn(II) dosing was similar between groups, we measured intensities in largest cubic region possible within brain, muscle, and non-tissue (noise) in unprocessed images from each dataset (N = 24 mice). Noise was calculated as the standard deviation of intensities within the non-tissue block. Mean signal was calculated for each tissue. SNR for each image was the mean signal intensity divided by the noise. We compared SNR across rearing groups using a linear mixed effects model in R (*nlme*) (16) with a main effect of rearing and with interactions of rearing by condition and by tissue (***SI Appendix*** **Fig. S2D**).

*4.4. Quality Control of Alignments and Jaccard Similarity.* To evaluate the quality of image alignments, we performed both qualitative and quantitative assessments. As a qualitative assessment, we averaged images for each group at each condition and visualized them to evaluate anatomical detail (***SI Appendix*** **Fig. S2D**). To assess the quality of alignments quantitatively, we calculated two commonly used similarity indices: Jaccard Similarity and Normalized Mutual Information (NMI).

Jaccard Similarity is a measure of overlap between the brain volume of each aligned image with a reference, which in this case is our dataset average (***SI Appendix*** **Fig. S2D, left**) (12, 17). To calculate Jaccard Similarity Indices for each image, all images must be converted to a compatible binarized format. Using *fslmaths*, we binarized each aligned image ($M_i$) as well as the dataset average ($M_{Avg}$) (18). All voxel intensities that exceeded 500 were set to 1 and all voxel intensities less than 500 were set to 0. This process isolates brain voxels for each image. The values for $M_i$ or $M_{Avg}$ are the binary value (0,1) at the x-y-z position of each voxel in the 3D brain matrix.

The Jaccard Similarity Index is defined as the size of the intersection of an individual image with the dataset average (numerator), divided by the size of their union (denominator) ***[Eq 1]***.

$$\mathrm{J}(\mathrm{M_i}, \mathrm{M_{Avg}}) = \frac{|\mathrm{M_i} \cap \mathrm{M_{Avg}}|}{|\mathrm{M_i} \cup \mathrm{M_{Avg}}|} \qquad (1)$$

If a brain voxel in the two images overlap, the multiplicand of the voxel intensities is 1. If not, the multiplicand is 0. Thus, the intersection of two binary images is equivalent to the sum of their voxel-wise product ***[Eq 2]***.

$$\left|\mathrm{M_i} \cap \mathrm{M_{Avg}}\right| = \sum_{\mathrm{v}=1}^{\mathrm{Voxels}} (\mathrm{M_{i,v}} \times \mathrm{M_{Avg,v}}) \tag{2}$$

$$\text{where } \mathrm{M_{i,v}} \times \mathrm{M_{Avg,v}} = \begin{cases} 0: \mathrm{M_i} = 0 \ \textbf{or} \ \mathrm{M_{Avg}} = 0 \\ 1: \mathrm{M_i} = 1 \ \textbf{and} \ \mathrm{M_{Avg}} = 1 \end{cases}$$

These calculations can be performed using FSL functions *fslmaths* and *fslstats.* Procedure to obtain the intersection in the command line…

Step 1: Create a voxel-wise binary map of overlapping brain voxels between the two images.

```
fslmaths <M_i filename> -mul <M_Avg filename> <Intersection Output> -odt char
```

Step 2: Calculate the total number of overlapping brain voxels identified in step 1.

```
fslstats <Intersection Output> -V
```

If a brain voxel is in both an individual image and the average image, the sum is 2. If a voxel is only in one of the two brain images, the sum is 1. If a voxel is not in either of the two brain images, the sum is 0. Thus, after summation of the two images, the number of voxels with a value >1 is equivalent to the union. If we binarize the summation image, the union is equivalent to sum of voxel intensities ***[Eq 3]***.

$$\left|\mathrm{M_i} \cup \mathrm{M_{Avg}}\right| = \sum_{\mathrm{v}=1}^{\mathrm{Voxels}} bin(\mathrm{M_{i,v}} + \mathrm{M_{Avg,v}}) \tag{3}$$

$$\text{where } bin(\mathrm{M_{i,v}} + \mathrm{M_{Avg,v}}) = \begin{cases} 0: \mathrm{M_i} = 0 \ \textbf{and} \ \mathrm{M_{Avg}} = 0 \\ 1: \mathrm{M_i} = 1 \ \textbf{or} \ \mathrm{M_{Avg}} = 1 \end{cases}$$

Procedure to obtain the union in the command line…

Step 1: Create a voxel-wise binary map of brain voxels from both images.

```
fslmaths <M_i filename> -add <M_Avg filename> -bin <Union Output> -odt char
```

Step 2: Determine the total number of overlapping brain voxels identified in step 1.

```
fslstats <Union Output> -V
```

The numerical output from the intersection can then be divided by that of the union to calculate the Jaccard Similarity Index between each image $(\mathrm{M_i})$ and the average $(\mathrm{M_{Avg}})$.

For more fine grain detail of anatomical similarity after alignments, we used normalized mutual information (NMI) (***SI Appendix*** **Fig. S2D, right**). NMI is often used as an optimization cost for MR image alignment algorithms (19). NIfTI images were loaded into R and the NMI between brain voxels in each image and the dataset average calculated via the *NMI()* function from the *aricode* package (20). Minimum and maximum intensity values were set to 1000 and 15000, respectively, and the number of histogram bins to 256. Calculated NMI was consistently above 0.9.

*4.5. Modal Scaling Evaluation.* We performed modal scaling in batch to adjust intensity differences across the combined dataset using a random image from the dataset as reference (template) (13). In this approach, the mode, or peak, of the histogram of voxel intensities represents the intensity value for the majority of voxels. We assume this peak is not altered by localized Mn(II) accumulation (1, 2, 13, 21-26). Modal scaling is an automated MATLAB-based set of functions that: 1) determines the intensity values corresponding to image modes; 2) estimates intensity histograms using a nonlinear least-squares curve fitting function and calculates the scaling factor to align histogram modes; and 3) multiplies this scaling factor to adjust the intensity values of the target images to that of the reference image. To be assured that original intensity distributions were preserved and that intensity values across all images fell within the same grayscale, we compared histograms before and after scaling relative to the reference image for every image in each group (***SI Appendix*** **Fig. S2E**). Results for grayscale normalization were consistent between groups and across conditions.

**5. MR Image Analysis.**

*5.1. Comparison of Anatomy Between Groups, Jacobian Determinant of Deformation Fields.* We tested for difference in anatomy between groups by obtaining the warp fields required to align their pre-Mn(II) images to the average of aligned pre-Mn(II) from Std, the minimal deformation target (MDT). The MDT was created by a two-step process as we previously described in Medina et al. 2017 (2, 13). One image from the skull-stripped Std pre-Mn(II) dataset was used as a reference and all other images in the dataset were rigid-body-aligned to this single image by SPM *Realign*. Aligned images were averaged using SPM *ImCalc* to produce a second reference image for another round of rigid body alignments of the dataset. The average from this second round created the MDT. The next steps were all performed with both groups processed in batch. All non-aligned Std and ELA pre-Mn(II) images were then rigid body aligned to the MDT using SPM *Realign* to prepare images for warping. All *Realigned* images' intensity histograms were modally scaled to the histogram mode of the MDT image with our custom MATLAB scripts (13). Then, intensity scaled images were affine aligned and nonlinearly warped to the MDT using SPM8 *Normalize* to obtain the warp fields for each dataset. For the *Normalize* function, estimation options were: Source image smoothing = 0.3mm; Non-linear cutoff frequency = 1; Non-linear regularization = 10; Non-linear iterations = 16. These first steps were performed in SPM8 producing an "sn.mat" file (27). To calculate Jacobian determinants for each voxel, the resultant "sn.mat" files from each pre-Mn(II) image's warping were processed in batch using the SPM12 *Deformations* function and SPM batch scripting (14). This process created voxel-wise Jacobian determinant "j_" files for all images including both groups. We inversely aligned the *InVivo* Atlas grayscale and label images to the MDT using a similar process as described in Extended Methods 5.4 below. Next, atlas masks were generated and applied to Jacobian determinant images to extract segment-wise averages using *fslmaths* and *fslstats*. Data were compiled into a csv file retaining group identity. Two-tailed two-sample t-tests comparing segment-wise Jacobian determinants between groups were performed in Excel for each segment with FDR correction ($p < 0.05$, FDR = 0.05). Total volume differences (in mm) were calculated by multiplying the average group differences of the Jacobian determinant values by the total segmental volume of the aligned *InVivo* Atlas.

*5.2. Statistical Parametric Mapping (SPM): Selection of Statistical Thresholds.* SPM assumptions are that: 1) error fields are a reasonable lattice approximation to an underlying random field with a multivariate Gaussian distribution, and 2) error fields are continuous, with an analytical autocorrelation function (14, 28). Aligned, normalized 3D MEMRI images do not meet these assumptions and hence must be "smoothed". Smoothing involves adjusting voxel intensity values to be continuous using a Gaussian kernel. The minimal "smoothness" is typically considered to be 3x the voxel size (14, 28), i.e., ~0.3 $mm^3$ for our data at 0.1 $mm^3$. A full-width half-maximum (FWHM) Gaussian kernel of 0.15 $mm^3$ returned the required smoothness, as measured by SPM *Results*. This kernel was applied to all images prior to SPM t-tests.

For all SPM t-tests we corrected for multiple comparisons by requiring statistically significant voxels be within a cluster of 8 and that p-value be less than $\alpha$ (0.05). Our goals for the selection of statistical thresholds were to balance specificity, based on simulated data, with estimated sensitivity, based on our power analysis of real data. First, we generated noise-only images (n=24) based on our real data's noise distribution (1000 ± 300 SD, assuming it to be Gaussian). Second, noise images were smoothed with the optimized 0.15 $mm^3$ Gaussian kernel. Lastly, we performed SPM t-tests on these noise-only images, for which any positive results are false. We then quantified the number of false positives, which yielded the false positive rate. We applied different cluster sizes and T-values until we found a combination that produced a false positive rate at or below $\alpha$ (0.05). At a cluster size of 8 and $T \geq 1.72$, our false positive rate was below 0.05, matching that of our power analysis.

To balance correction for multiple comparisons with sensitivity, we also considered the ability to detect moderate effect sizes. a cluster size of 8 ensured both a corrected false positive rate of less than α (0.05) and simultaneously allowed for statistical detection of moderate effect sizes. For comparisons of post-Mn(II) images, these settings captured Cohen's D effect sizes of $d > 0.52$ for within group and $d > 0.72$ for between group comparisons, at $p < 0.05$ cluster corrected. For comparisons of post-Mn(II) versus pre-Mn(II) images, where signal is much stronger, we kept the cluster size the same (8 voxels) but increased t-value stringency, which improved comparisons between segmental fractional activation volumes, as a number of segments were saturated (approached 100% active) at lower stringency.

*5.3. SPM: Adjustment of Post-Mn(II) Images for Average Differences Between Groups' Pre-Mn(II) Images.* We found small, yet statistically significant differences in signal intensity when comparing pre-Mn(II) images between groups. To account for such differences, we "standardized" the ELA group by adjusting all ELA images to the same intensity profile as Standard. To do this, aligned, intensity normalized, and smoothed images of pre-Mn(II) were 1) averaged for each group; 2) the Standard average was subtracted from the ELA average; and 3) the resultant difference image was subtracted from all pre- and post-Mn(II) images of the ELA group. To verify that this adjustment nullified baseline differences, we compared pre-Mn(II) images of Standard with the adjusted pre-Mn(II) images of ELA and found the differences between groups disappeared. All between group comparisons in Fig. 6 used the adjusted post-Mn(II) ELA images. Adjustment calculations were performed with *fslmaths* (18).

*5.4. InVivo Atlas: Alignment and Automated Segmentation.* For this manuscript we added regions to our *InVivo* Atlas previously reported in (1), which is a high resolution MEMRI image of a living mouse brain with annotated segments defined by manual segmentation. New segments were added by drawing them on the "label image" using FSLeyes for visualization. Segment boundaries were based on grayscale contrast and the Allen Institute for Brain Science's Mouse Reference Atlas – the Common Coordinate Framework (CCF v3) – and are annotated according to that terminology (29). Segments were assigned to larger domains based on the CCF v3 hierarchy. The Atlas now includes 117 annotated brain segments in *Version 10.4* versus 87 in our previous report (1) (***SI Appendix*** **Table S6**). New segments include many sub-regions of the cortex, hypothalamus, and midbrain. Of the total 117 segments, 102 were used for statical map segmentation and calculation of fractional volumes. The other 15 were fiber tracts or ventricles/CSF segments, which were used to mask statistical maps. Anatomical volumes were also compared in these 15 other segments as in *5.1* above.

The updated *InVivo* Atlas was inversely aligned to our dataset. First, the lower resolution dataset average image was linearly aligned to the higher resolution atlas via FSL flirt. The transformation matrix was inverted and then applied to the grayscale atlas and the label image with nearest neighbor interpolation. Next, the grayscale atlas was non-linearly aligned to the dataset average and the warp field applied to the labels, again with nearest neighbor interpolation. We used a suite of custom written bash and python scripts with *fslmaths* and *fslstats* to 1) create segment masks, 2) apply segment masks to SPMs, and 3) extract segment-wise statistical summaries, such as average t-value and the number of statistically significant voxels (1, 30). After segment-wise voxel counts were extracted, resultant data were compiled into a csv file and loaded into R for processing and visualization. Fractional volumes, i.e., the fraction of statistically significant to total voxels, were calculated in R (see next section for formulation). Column graphs were generated using the *ggplot2* package in R.

*5.5. Formulas Used to Calculate Fractional Volumes.* Statistical maps were used to identify volumes of statistically significant Mn(II)-enhanced intensity voxel-wise per segment (14), and the fraction of enhanced to total voxels calculated.

Two SPM designs were performed for each of the three conditions: 1) paired t-tests of post-Mn(II) images relative to pre-Mn(II) or relative to other post-Mn(II) images (FAV); and 2) unpaired two-sample t-tests between ELA and Standard post-Mn(II) images (FDV). From segmentation of respective SPM T-maps, we calculated the fraction of total voxels that were significantly enhanced ***[Eq 1]*** or significantly different ***[Eq 2]*** within each segment ($S_x$). We defined the fraction of enhanced to total as "fractional activation volumes" ***[Eq 1]*** and the fraction of statistically different to total as "fractional difference volumes" ***[Eq. 2]***.

$$\text{Fractional Activation Volume}_{S_x} = \frac{\text{Count(Significant Voxels}_{S_x,\text{Paired}})}{\text{Total(Voxels}_{S_x})} \quad (4)$$

$$\text{Fractional Difference Volume}_{S_x} = \frac{\text{Count(Significant Voxels}_{S_x,\text{Unpaired}})}{\text{Total(Voxels}_{S_x})} \quad (5)$$

Where $S_x$ stands for any individual segment "x," *e.g.*, BST, ACA, or DG.

*5.6. Evaluation of Potential Sex Differences.* The ELA group was composed of equal numbers of male (n = 6) and female (n = 6) mice. We sub-divided smoothed post-Mn(II) images form the ELA group by male and female, and compared them by SPM t-tests at each condition ($p < 0.01$, cluster corr.). Resultant statistical maps were displayed in FSLeyes with our *InVivo* Atlas, and the between group (ELA vs Std) statistical maps overlaid (***SI Appendix*** **Fig. S10**).

*5.7 Diagrams of the Balance of Brain Wide Patterns of Activity.* Diagrams of approximate anatomical location of *InVivo* Atlas domains and relative levels neural activities within these domains illustrate brain states characterized by fractional activation (***SI Appendix*** **Fig. S11**) and difference volumes (**Fig. 6**, **right panels**) of Standard and ELA groups over the three conditions. Color gradients were used to depict relative differences in fractional volumes among regions within and between brain states, with darker colors correlating with greater average fractional volumes across segments in that domain. For **Fig. 6**, the proportion of area taken in each box by the two colors represents the proportion of ELA versus Std FDV across segments. Color-coded text blocks of the 10 domains were manually positioned in approximate anatomical locations on a sagittal slice acquired with FSLeyes.

**6. Immunohistochemistry, c-fos Quantification and Correlation to MEMRI.**

*6.1. Immunohistochemical Staining for c-fos and Microscopy.* At the conclusion of MR imaging, subsets of mice (5 Std and 6 ELA) were deeply anesthetized with avertin (250-500 mg/kg i.p., Sigma Aldrich) and subjected to cardiac perfusion as we previously described (1, 31). Fixed brains of each group were embedded in a single block, and the blocks serially sectioned in register at 35 μm intervals (Neuroscience Associates, https://www.neuroscienceassociates.com). Sections were stained by immunohistochemistry using a rabbit polyclonal anti-c-fos primary (Cat# RPCA-c-fos, EnCor Biotechnology) and a biotinylated anti-rabbit secondary with Nickel DAB chromogen. Micrographs were obtained on an Olympus C643 light microscope with a DP27 digital camera and CellSens software. Anatomical sections also identifiable in MR slices were selected for measurement of counts and analysis: hypothalamus (high) and thalamus (low) in MR slice 102; superior central raphe (high) and midbrain (low) in MR slice 107. For c-fos IHC, image intensities visualized in ImageJ/FIJI for counting. Nuclei were counted manually in the micrographs on the computer screen using a 3 x 3 grid of 100 $\mu m^2$ squares in 1-3 sections for each animal (5 Std and 6 ELA). We aimed to count 3 x 35 μm serial sections for a ~100 $\mu m^3$ volume. For regions with more than 1 section counted, we averaged the counts in each anatomically correlated 100 $\mu m^2$ square. MR intensities (SI) were measured from 100 $\mu m^3$ voxels in corresponding slices.

*6.2. Quantification and Correlation c-fos+ counts and MEMRI SI.* <u>Data processing</u>: Counts of c-fos+ nuclei and signal intensities from voxel-wise MEMRI were loaded, processed, and analyzed in R. Raw counts and SI were visualized in quartile boxplots using *ggplot2* in R. To meet the linearity assumption of linear regression and Pearson's correlation, we applied a square root transform to c-fos counts from each 100 $\mu m^2$. Regions counted in the same section or measured in the same slice were compared by linear mixed models, accounting for random effects of individual, and the fixed effect of region by post-hoc paired t-tests with FDR correction. For correlation analysis, the square-root-transformed counts as well as SI values were averaged by individual and region. Counts were paired with SIs for each of the 4 regions for all 5 Std (20 total; 5 mice x 4 regions); or 6 ELA brains (24 total; 6 mice x 4 regions).

Permutation testing: The 20 Std and 24 ELA averaged linearized counts were paired with SI values and randomly permuted 10,000 times each, where n = 20 or 24 per resample, respectively. Pairings of counts and SI values were jumbled simultaneously across individuals and regions. We calculated the Pearson correlation coefficient when the counts and SI values were jumbled, which yielded a null distribution for each group. The p-value, corresponding to the likelihood of the correlation between measures to be due to chance, was determined by calculating the proportion of values in the permuted data that were equal to or greater than the magnitude of the calculated coefficient from the original sample. A one-tailed test was performed to test the hypothesis that correlations between counts and SI would be positive. If the resultant p-value were greater than 0.05, this would refute our hypothesis. If less than 0.05, this would support our hypothesis. See ***SI Appendix*** **Fig. S5C-D**.

Bootstrap resampling: We performed bootstrap resampling to gain a better estimate of the correlation between c-fos counts and MEMRI SI (32). Unlike permutation testing, pairing was maintained for resampling. The 20 normal and 24 ELA count-SI pairs were resampled separately with replacement, each 10,000 times, where n = 20 or 24 per sample. For each re-sample, the Pearson correlation between counts and SI values was calculated, as well as average counts and SI values. This yielded a larger sample size and the bootstrapped sampling distribution, for these statistics, as shown in ***SI Appendix*** **Fig. S5C-D**. We performed a linear regression of the average SI to average count data to determine the equations for the lines shown in **Fig. S5D**.

**SI Figures S1 to S11**

**Fig. S1.**

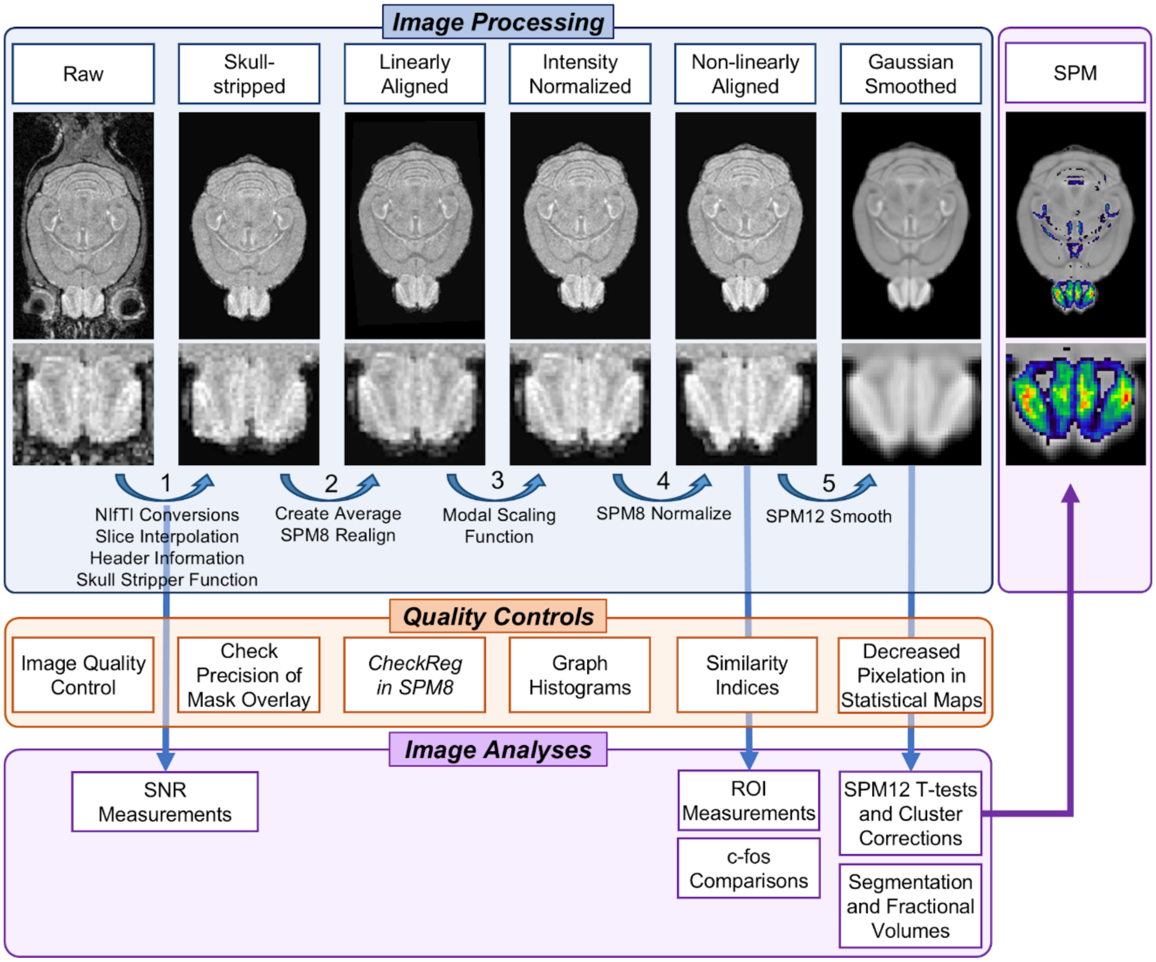


**Fig. S1.** Flow Diagram of MEMRI Image Processing and Analysis Pipeline.

Diagram illustrates image processing, statistical analysis, and quality assurance workflow. Note our processing, as described in Medina et al. 2017, is semi-automated involving a series of automated algorithms that require user input and visual inspection for quality assurance. _Top Panel_: MEMRI Image Processing steps, from raw Bruker images to Gaussian smoothing. Brain-wide axial slices of a representative HC MEMRI image with higher magnification images of olfactory bulb anatomy, corresponding to each processing step. Curved blue arrows represent processing performed between images. Note that anatomical contrast, both whole-brain and olfactory, are preserved through *Step 4: SPM12 Normalize*, i.e., images are skull-stripped, intensity normalized and anatomically aligned. Smoothing, which is required to meet assumptions of Random Field Theory for SPM, decreases effective resolution by applying 0.15 $mm^3$ Gaussian kernel. Non-smoothed images are subject to higher voxel-wise variability and can result in pixelated statistical maps. _Middle Panel_: List of quality controls taken at each processing step. Examples of these quality controls are shown in ***SI Appendix*** **Fig. S2**. _Bottom Panel_: Image analyses from input images corresponding to particular steps in the processing pipeline. SNR measurements are taken from images prior to the final stage of *Step 1: Skull-stripping*. ROI measurements in **Fig. 3-4** and ***SI Appendix*** **Fig. S4-6** are performed after *Step 4: SPM12 Normalize* on warped-aligned images. Note that smoothed images were also compared against warped as a control ***SI Appendix*** **Fig. S6**. SPM12 t-tests in **Figs. 5-6** and ***SI Appendix*** **Fig. S3, S7**, **S8**, and **S10** are performed after *Step 5: SPM12 Smooth*.

**Fig. S2.**

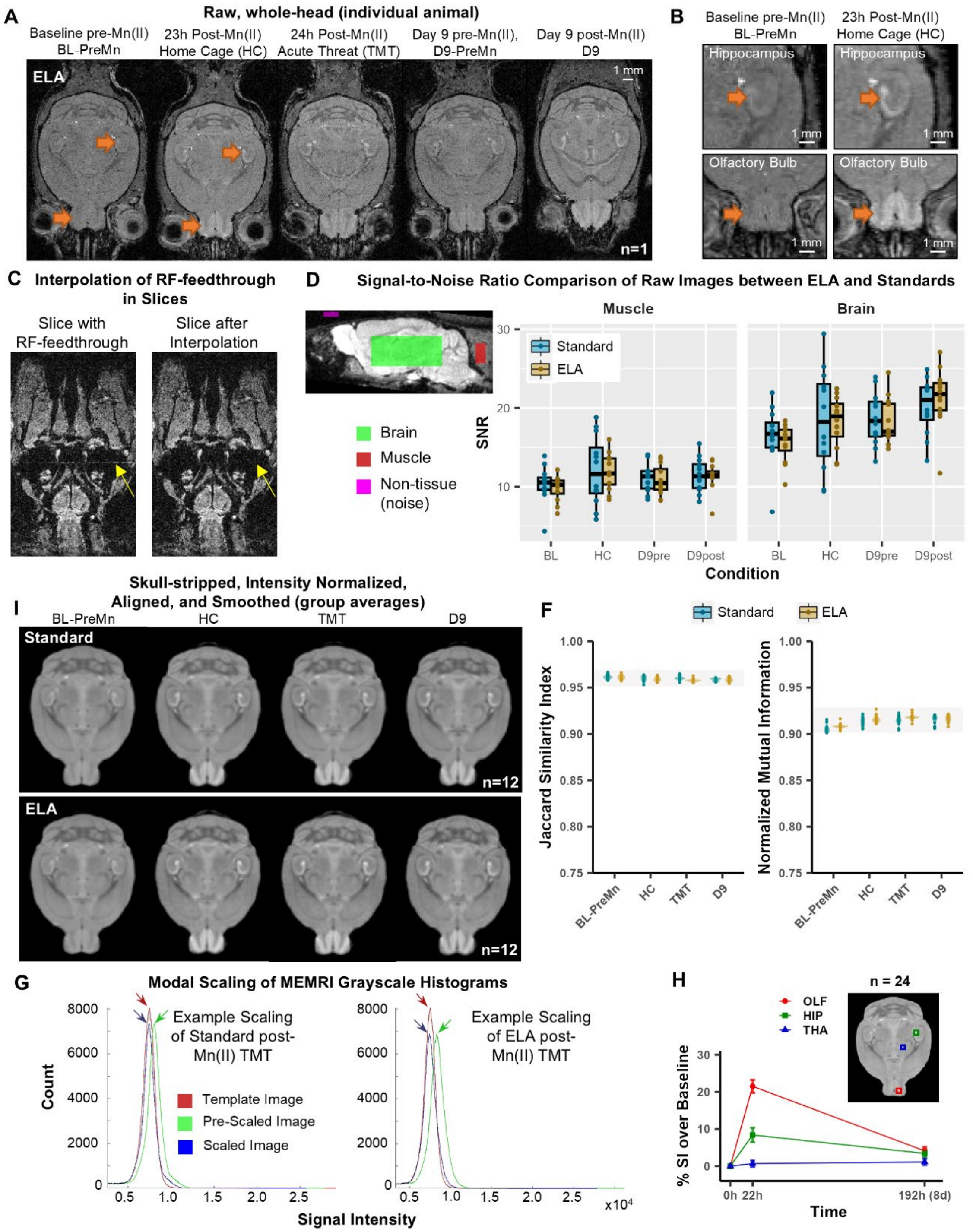


**Fig. S2.** MRI Image Processing Quality Controls and Mn(II)-enhanced Signal Kinetics. *(A)* A representative set of unprocessed whole-head MR images from a single ELA animal across the longitudinal timeline (n = 1). Orange arrows indicate regions that are visually enhanced by Mn(II). See Uselman et al. 2020 for similar projections of Std (2). *(B)* Higher magnification of pre-Mn(II) "baseline" (*left*) and post-Mn(II) images at HC (*right*) from panel *(A)* showing Mn(II)-enhanced signal increases (orange arrows) in hippocampus (*top*) and olfactory bulb (*bottom*). This demonstrates expected Mn(II)-enhancement and anatomical contrast of images *(C)* Axial slices of whole-head images showing an RF-feedthrough "slice" artifact (*left,* arrow) and the same slice after interpolation (see **Extended Methods 4.1**). In these slices where RF-feedthrough artifacts appeared, linear interpolation of adjacent slices successfully removed artifacts. *(D) Left panel*: Representative brain (green), muscle (red), and non-tissue (noise, magenta) volumes overlaid on a representative raw grayscale MEMR image. Intensity measurements from these volumes across pre- and post-Mn(II) images at BL, HC, and D9-Pre, D9-Post (as indicated) were used to compare signal-to-noise ratios (SNR) between groups. *Right panel*: Boxplots of SNRs calculated from average Muscle (left) and Brain (right) tissue intensity after adjustment for noise. Noise levels were consistent throughout. SNR is compared between pre- and post-Mn(II) scans for each injection, and between groups (Std, cyan; ELA, yellow). Comparisons found that MRI acquisition produced similar SNR between groups at all four timepoints in both brain and muscle, validating that our protocol delivered similar $MnCl_2$ doses for all animals in both groups. No main effect of group, or interactions of group with tissue location or Mn(II)-injection were observed by ANOVA (Group: $F_{(1,22)} = 0.469$, $p = 0.500$; Group x Tissue: $F_{(1,160)} = 0.049$, $p = 0.825$; Group x Mn(II): $F_{(3,160)} = 0.354$, $p = 0.786$). *(E)* Standard (Std, *top row*) and ELA (*bottom row*) group averages of skull-stripped, intensity normalized and spatially aligned images across each point in the timeline (n = 12). Note the enhanced anatomical detail. *(F)* Violin plots of Jaccard Similarity Indices (JSI) and Normalized Mutual Information (NMI) for (linearly, non-linearly) aligned Std (cyan) and ELA (yellow) MR images relative to the dataset average. Both groups show JSI > 0.95 and NMI > 0.9 for all images, indicating a high degree of alignment across the dataset. No large-scale difference exists in either JSI ($F_{(1,22)} = 0.3$, $p = 0.61$) or in NMI ($F_{(1,22)} = 1.0$, $p = 0.34$) between Std and ELA. *(G)* Modal scaling of grayscale intensity histograms from representative images in Std (*left*) and ELA (*right*) groups. Intensity normalization occurs through scaling intensity values of all images in the dataset to the mode of the histogram of a template image (red). Precision of grayscale normalization is evidenced by coincidence of intensity histogram modal peaks of images with that of the reference image. *(H)* Average kinetics of Mn(II)-enhanced signal of both ELA and Std datasets in the olfactory bulb (red), hippocampus (green), and thalamus (blue) from BL-PreMn (0 h post-Mn(II)) to HC (22 h post-Mn(II)) to D9-PreMn (192 h post-Mn(II)). Each are shown as Mean ± SEM the percent signal intensity (SI) over BL-PreMn combining groups (n = 24). Colored boxes indicate regions measured. Mn(II) intensity kinetics over time reproduced previous reports (1, 11).

**Fig. S3.**

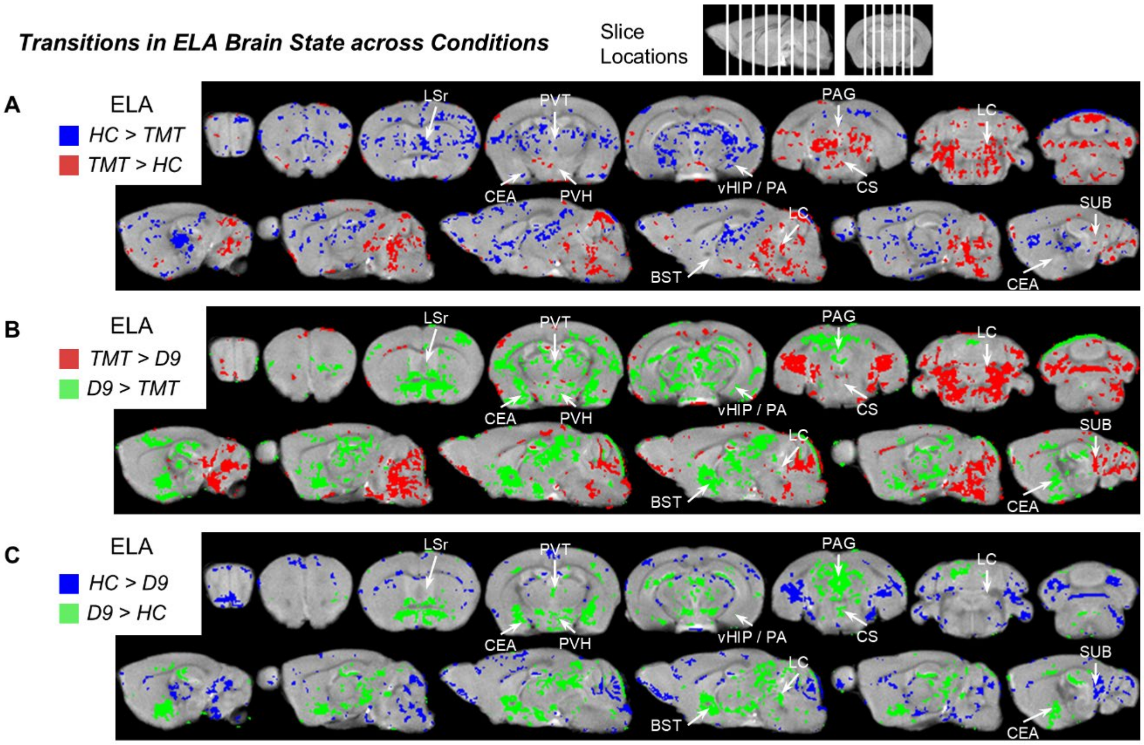


**Fig. S3.** Acute and Longer-term Transitions of ELA Brain States as they Progress across Conditions. *(A-C)* Results from pairwise SPM t-tests of ELA images between sequential post-Mn(**II**) conditions and between HC and D9 (T = 1.80, $p < 0.05$, cluster corrected for multiple comparisons, minimum cluster size = 8). Note this comparison does not show HC activity as does Main Text **Fig. 2** but does show both increased and decreased activity between HC and post-threat conditions and between short- and long-term post-threat conditions. Shown are coronal (*top rows*) and sagittal (*bottom rows*) slices with statistical maps overlaid on the dataset average. White arrows point to segments with large changes in activity between conditions. Colors represent direction of signal differences as indicated. *(A)* HC signal intensity versus acute TMT. TMT elicited an acute shift in activity to include various mid- and hindbrain segments, such as the LC. Whereas activity in the LSr decreased after TMT. *(B)* D9 signal intensity versus acute TMT. Various new volumes of activity emerge at D9 relative to TMT: PVH, PVT, vHIP/PA, CEA, and PAG. Other regions, such as the subiculum (SUB, not segmented), saw a decrease in activity at D9. *(C)* D9 signal intensity versus HC. Various regions show heightened activity at D9 over activity already present at HC or TMT: PVT and PAG. Other regions such as the PVH, PVT, BST, vHIP/PA, CS, and CEA are activated primarily at D9.

**Fig. S4.**

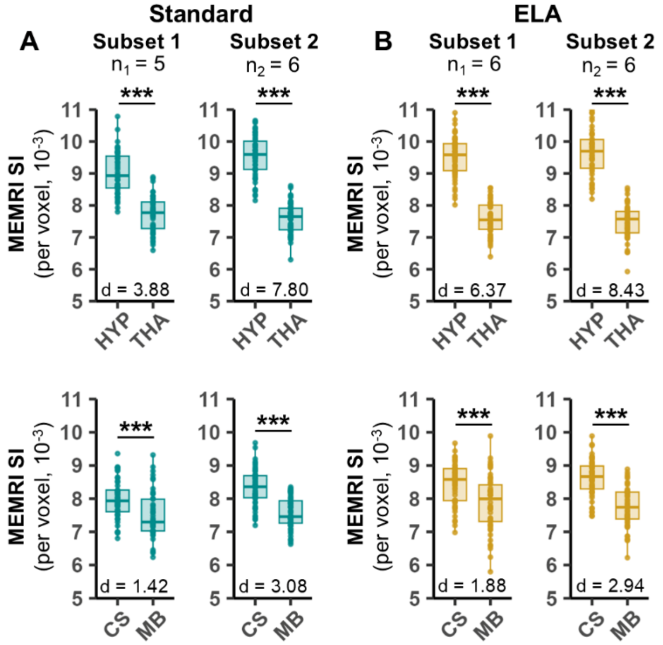


**Fig. S4.** Test-retest Reliability in MEMRI Signal Intensities between Subsets of Std and ELA Mice. We evaluated test-retest reliability by comparing statistical differences between MEMRI signal intensity (SI) values in the two regions, as in Main Text **Fig. 4**. We divided each group into two subsets (Std: $n_1 = 5$, $n_2 = 6$; ELA: $n_1 = 6$, $n_2 = 6$) and compared between the subsets within group. In subset 1 ($n_1$) individuals were tested singly two weeks apart over a 10-12 wk period. Subset 2 ($n_2$) was tested months later, again over a 10-12 wk period. We reasoned that if this protocol were reproducible and reliable, we would find only minimal variance between individuals and subsets, and similar statistical significance of their between region differences.

The twelve remaining post-Mn(II) D9 images from the larger dataset of both *(A)* Std (cyan) and *(B)* ELA (yellow) were used to compare to the subset analyzed in Main Text **Fig. 4**. Quartile boxplots with dot plot overlays display statistical differences between MEMRI SI of regions for each subset of Std or ELA mice. Separate mixed effects models with a fixed effect of region and random effects of individual and voxel location were used for each subset of each group. Post-hoc paired t-tests FDR correction, *** $p < 0.001$. Effect sizes (*d*) were estimated by Cohen's D. Statistical differences between regions with high and low signal intensity were preserved in the two subsets. For example, in *(A)* for Std, the significance between HYP and THA in subset 1 ($n_1$) was $p < 0.001$, FDR, also with the same p-value in subset 2 ($n_2$).

**Fig. S5.**

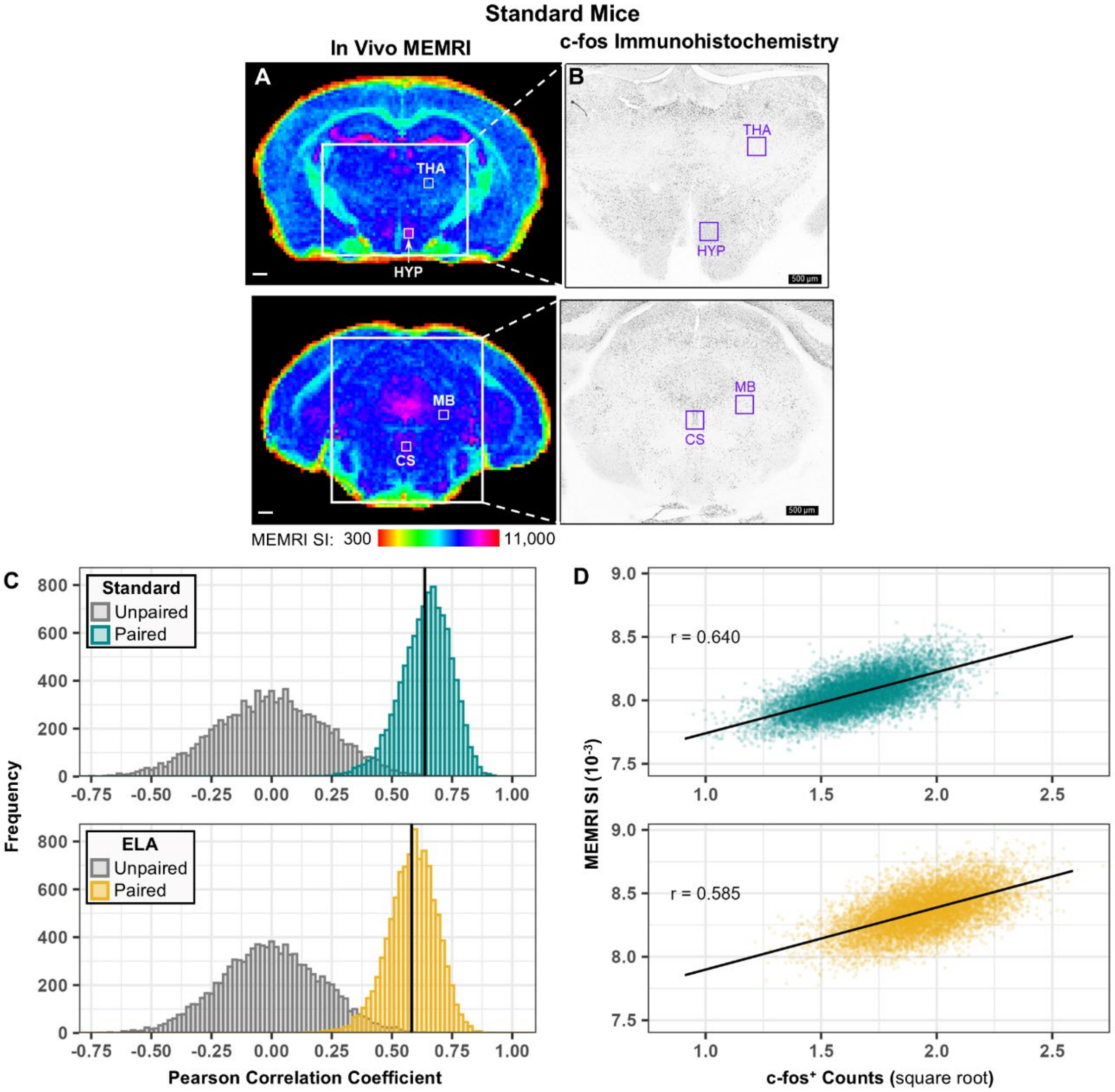


**Fig. S5**. Correlations between MEMRI SI and c-fos Counts. *(A)* Averaged images of Std mice (n = 5) at D9 from the same five brains used for c-fos counts (as shown for ELA in Main Text **Fig. 4**). These Std mice were sectioned and stained for c-fos in parallel with ELA mice shown in **Fig. 4**. Coronal slices are color coded for intensity values (HSV in *FSLeyes*, see MEMRI SI color code). Minimum was set to 300 based on noise and maximum to 11,000 according to the intensity histogram. Large white boxes indicate histological fields of view as shown in *(B)*; small white boxes indicate regions (3 x 3 x 1 voxel grid) where signal intensities (SI) and c-fos+ nuclei were measured. In slice 102, the area with high SI was hypothalamus (HYP) and with low, thalamus (THA). In slice 107, the area with high SI was superior central raphe (CS) and with low, midbrain (MB). White mag bar = 500 µm. *(B)* Representative histologic sections of a single Std brain from the same group of mice shown in (*A*) stained for c-fos. Purple squares indicate locations of 3 x 3 grid of 100 µm$^2$ squares where numbers of c-fos+ nuclei were counted in 1-3 x 35 µm thick sections. Black mag bar = 500 µm. *(C-D)* Correlations between MEMRI SI and c-fos counts. *(C)*

Histograms showing the distributions of bootstrapped resampled correlation coefficients between MEMRI SI and square-root-transformed c-fos+ counts (Std: *cyan, top;* ELA: *yellow, bottom*) and of coefficients when MEMRI SI and c-fos pairing were randomly assigned, i.e., permutation testing (gray histograms). Black line indicates the correlation coefficient of the original data. Probability of whether correlations were due to chance is represented by the area of the gray histograms to the right of the black line – correlations of the unpaired data equal to or greater than the correlation coefficient of the original data (black line; Std: $r = 0.636$, $p = 0.001$; ELA: $r = 0.582$, $p = 0.0013$). The number of bootstrapped samples and permutations were 10,000 each. See Extended Methods 6.2 for details. Note coefficients calculated from original data (black line) are similar to the peak of the histogram of the resampled data when pairing is maintained. *(D)* Scatter plots of bootstrap resampled pairs of average square-root-transformed c-fos counts and average MEMRI SI for Std (*top, cyan*) and ELA (*bottom, yellow*). Regression line (*black*) and corresponding average Pearson correlation coefficient (*r*) from bootstrapped samples from *(C)* are indicated. Note the distributions form a strong linear relationship ($r > 0.5$) in both groups; Std: $r = 0.640$ [0.408, 0.822]; ELA: $r = 0.585$ [0.376, 0.765]; where numbers in brackets are the 95% confidence interval of the bootstrapped data.

**Fig. S6.**

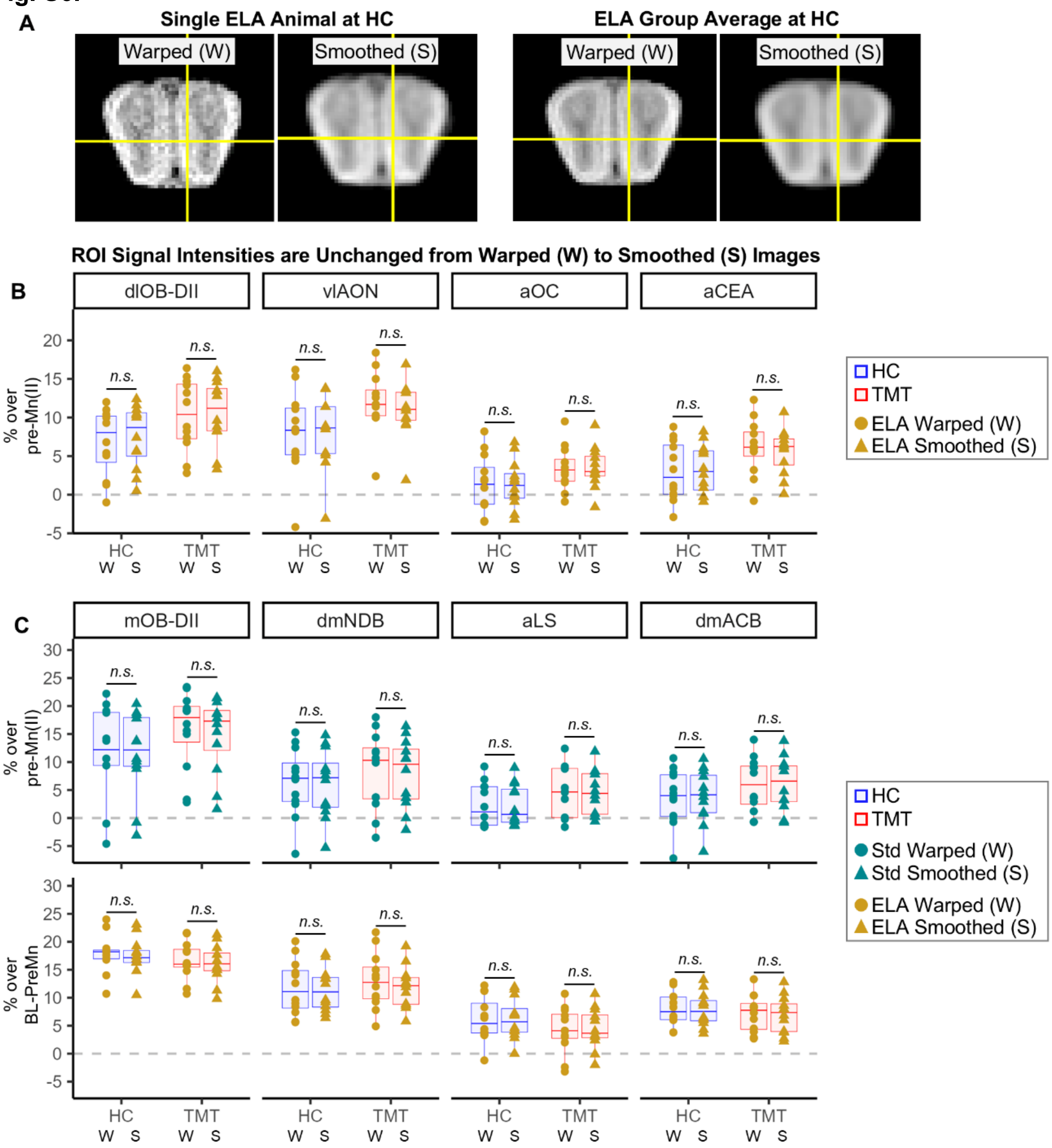


**Fig. S6.** Comparison of ROI Intensities between Warped Images before and after Smoothing at a Kernel Size of 0.15 $mm^3$.

*(A)* Example images of the medial DII subdomain of the olfactory bulb (mOB-DII) of a single ELA individual at HC (left two images) and from ELA group averages at HC (right two images) before (W) and after smoothing (S) with a 0.15 $mm^3$ kernel. Yellow crosshairs indicate center voxel of 3 x 3 x 3 mOB-DII ROI. Note olfactory bulb layers are still distinguishable in smoothed images. *(B)* Shown are signal intensities measured from the same four ROIs in ELA as shown in **Fig. 3A** at two conditions (HC, blue; TMT, red) before and after smoothing. Boxplots for warped (circles, W) and smoothed (triangles, S). The boxplots for W and S are shown side-by-side for each condition at each ROI. See key for color coding. Note no statistical significance between warped and smoothed

(*n.s.*, p > 0.05). *(C)* Shown are signal intensities from four other ROIs from both groups as shown in **Fig. 3B** at two conditions before and after smoothing. Std (*top*, cyan) and ELA (*bottom*, yellow). Boxplots for warped (circles, W) and smoothed (triangles, S). Boxplots for W and S are shown side-by-side for each condition at each ROI. No statistical differences were found between warped and smoothed images for any of the 8 measured ROIs, by repeated measures ANOVA; main effect of smoothing: $\eta_p^2 < 0.02$; post-hoc t-tests with FDR correction: *n.s.*, p > 0.05. No statistical interactions of smoothing with group or condition were found. Note the average difference in signal intensity due to smoothing across all data points was 0.2 ± 0.4% (± SD) in 3 x 3 x 3 ROIs. Smoothing decreases intensity in some voxels and increases intensity in others. The effect of smoothing on a voxel-wise level is complicated. By using the smallest smoothing kernel allowable we mitigate these effects. By using 3 x 3 x 3 voxel ROIs we are replicating the effective resolution of the smoothing process used here with a 0.15 $mm^3$ Gaussian kernel.

**Fig. S7.**

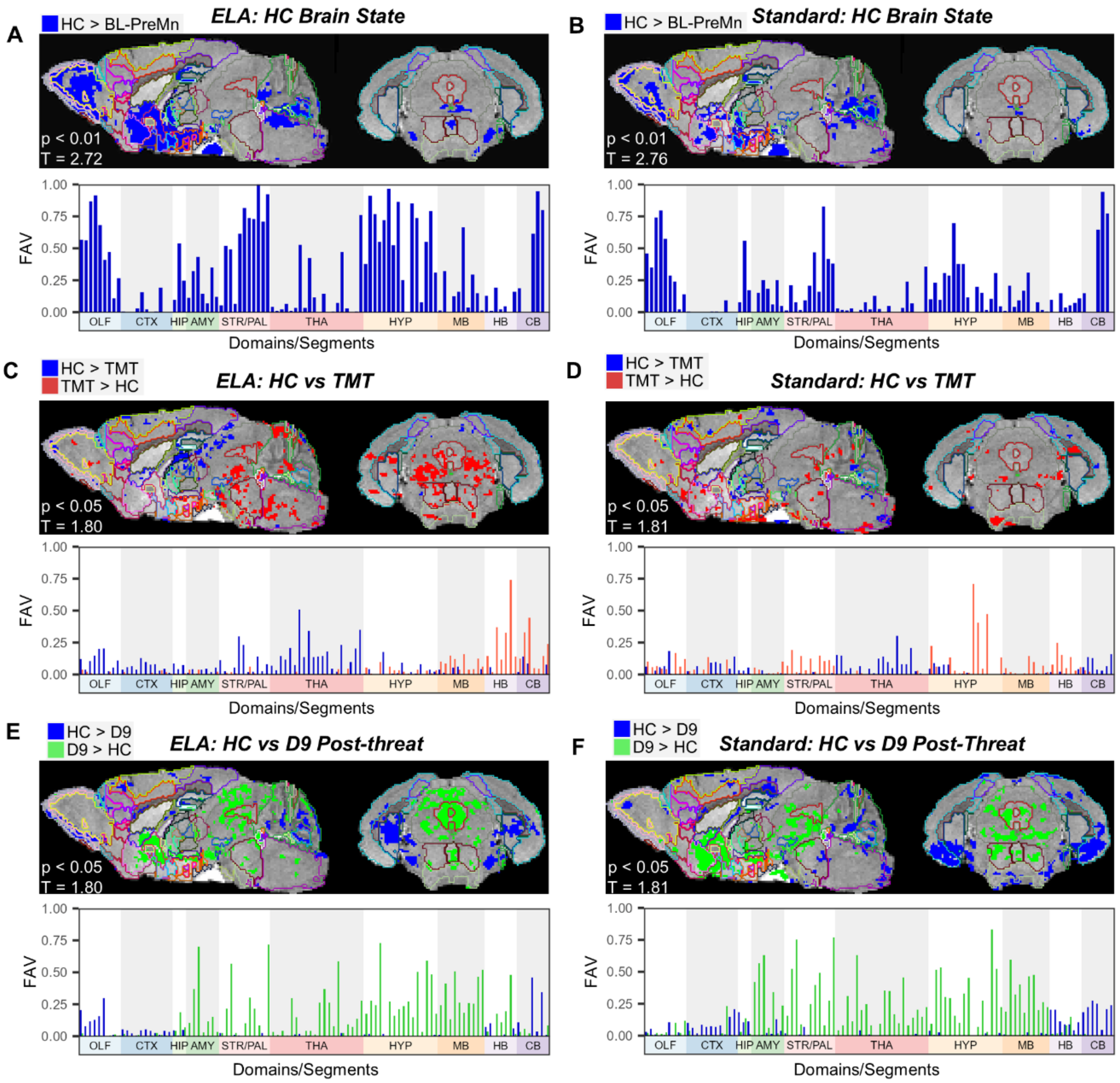


**Fig. S7.** Direct Comparisons between Successive Conditions in ELA and in Std. *Left panels*, ELA *(A, C, E)*; *Right panels*, Std *(B, D, F)*. Shown are statistical maps for each condition projected onto our segmented grayscale *InVivo* Atlas *(top* image*)* together with their corresponding column graphs *(below* each image*).* Column graphs show the fractional volume of activity measured quantitatively as segment-wise fractional activation volumes (FAV, vertical axis). Ten anatomical domains (color-coded on the horizontal axis) are divided into 102 subsegments. See ***SI Appendix* Table S5** for list of domains and subsegment names, and **Table S6B** for FAV values. Neural activity in *(A)* ELA and *(B)* Std at home cage (HC). Post-Mn(II) images at HC compared to that cohorts' pre-Mn(II) images, $p < 0.01$, cluster corr. for multiple comparisons; ELA: $T_{(11)} = 2.72$, n = 12; Std: $T_{(10)} = 2.76$, n = 11. Note FAVs across many segments are greater for ELA than Std. Activity in *(C)* ELA and *(D)* Std shift immediately after threat. Post-Mn(II) images at HC were compared to those at TMT. Direct comparisons enable detection of both activated and silenced regions by the experience of threat.

Colors of statistical maps and columns indicate these increased (red) or decreased (blue) signal intensities at TMT compared to HC. Note that the blue columns represent areas active prior to threat and silenced afterwards. In ELA, threat increased activity primarily in MB and HB ($FAV_{ELA}$ = 0.14-0.38) and silenced activity in anterior domains. In Std, domains that were already highly active in ELA at HC (STR/PAL and HYP) only became active in Std after threat ($FAV_{Std}$ = 0.18-0.21). Activity in *(E)* ELA and *(F)* Std transitions again longer after threat. Comparisons were between post-Mn(II) images at HC and D9. Colors of statistical maps and columns indicate increased (green) and decreased (blue) signal intensities at D9 compared to HC. Note brain-wide activity increased at D9 compared to HC in both groups, but to a greater extent in Std. Neither group regained the home cage pattern. These data suggest amount and anatomical distribution of activity is different after threat experience. For post-Mn(II) versus post-Mn(II) comparisons in *(C-F)*: paired t-tests, $p < 0.05$, cluster corr. for multiple comparisons; ELA: $T_{(11)} = 1.81$; Std: $T_{(10)} = 1.82$.

705 **Fig. S8.**

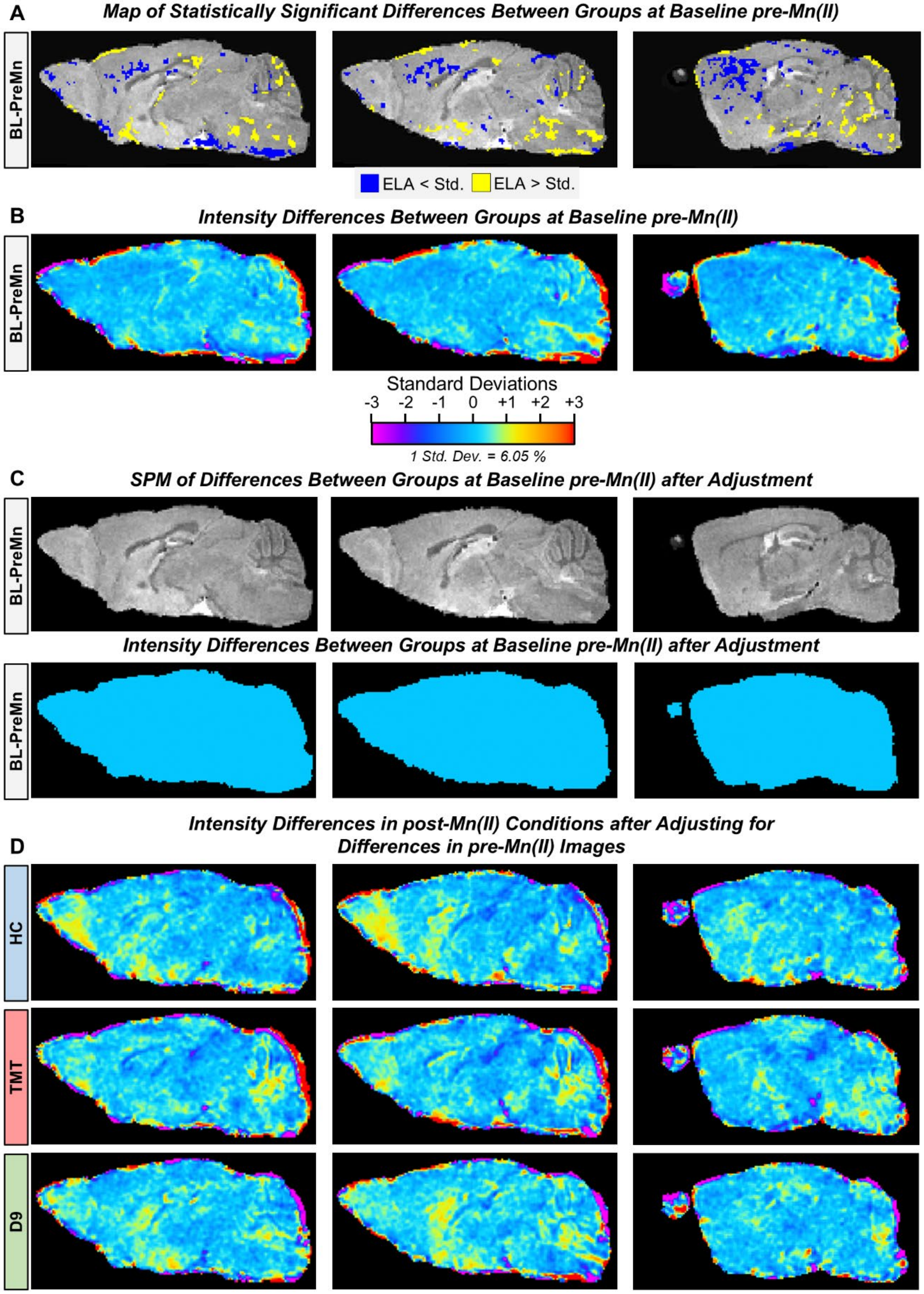

A
Map of Statistically Significant Differences Between Groups at Baseline pre-Mn(II)
BL-PreMn
ELA < Std.
ELA > Std.
B
Intensity Differences Between Groups at Baseline pre-Mn(II)
BL-PreMn
Standard Deviations
-3 -2 -1 0 +1 +2 +3
1 Std. Dev. = 6.05 %
C
SPM of Differences Between Groups at Baseline pre-Mn(II) after Adjustment
BL-PreMn
Intensity Differences Between Groups at Baseline pre-Mn(II) after Adjustment
BL-PreMn
D
Intensity Differences in post-Mn(II) Conditions after Adjusting for Differences in pre-Mn(II) Images
HC
TMT
D9

**Fig. S8.** Comparison of Baseline pre-Mn(II) Images between Groups and Effects of Adjustment on post-Mn(II) Intensity Maps. *(A)* Between-group SPM of pre-Mn(II) images show statistically significant differences in T1-weighted signal intensities between ELA and Std at low stringency ($p < 0.05$). Shown are three sagittal slices of these statistical maps overlaid on the dataset average: ELA > Std, yellow; ELA < Std, blue. $T = 1.72$, $p < 0.05$, cluster corrected for multiple comparisons. *(B)* Measurements of intensity differences shown as maps reveal that the significant voxels are due to minimal actual signal difference between groups. Sagittal slices from the 3D difference map after subtracting the average Std pre-Mn(II) image from that of ELA. Color code represents standard deviations of the percent differences of ELA from Std. Note that many differences in brain signal are less than 1 standard deviation from Std (< 6.05%). Only a few regions have larger differences and these typically correspond to those regions also found to be statistically significant. Differences at baseline between groups are therefore unlikely to contribute to post-Mn(II) results. *(C)* After adjustment of ELA images, no significant differences were found by SPM (*top*) or by average intensity difference maps (*bottom*) between baseline Std and ELA pre-Mn(II) images. The *InVivo* Atlas was used as a mask for the intensity difference maps in GIMP/Photoshop. *(D)* Measurements of post-Mn(II) intensity differences after adjusting for baseline differences between groups. Differences in post-Mn(II) signal remain large after adjusting for baseline pre-Mn(II) differences between groups. Sagittal slices of average difference between groups' post-Mn(II) images at home cage (HC), acute threat (TMT) and at day 9 (D9) after removing for the average differences at baseline as in *(B)*. Color code represents the standard deviation of difference between groups at pre-Mn(II), also as in *(B)*. Note that the magnitudes of signal differences between groups are now greater than 1-2 standard deviations in various brain regions. Thus, magnitudes of differences in post-Mn(II) images are frequently more than 2-fold greater than differences observed in pre-Mn(II) images and also remain significant even after adjusting for those baseline differences (see **Fig. 6**).

**Fig. S9.**

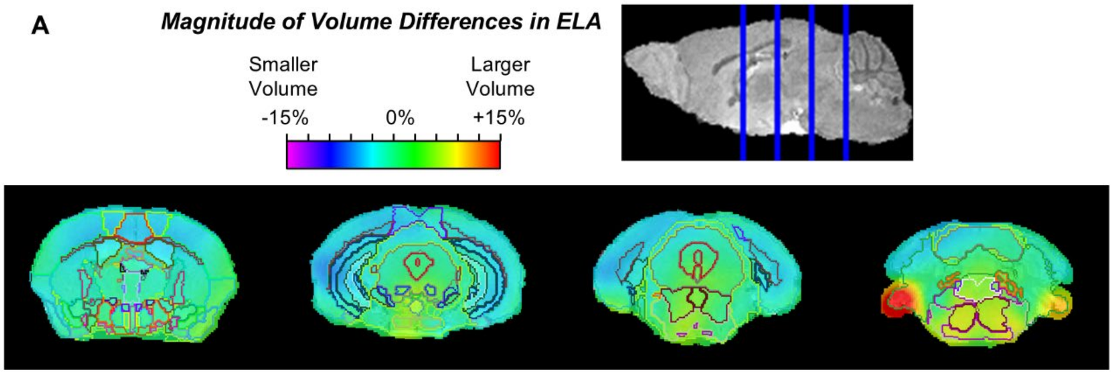


**Fig. S9.** Absence of Anatomical Differences between Adult ELA and Std Mice. Brain volumes were determined by calculating the Jacobian determinants of deformation fields from the warping alignment process. ELA and Std pre-Mn(II) images were warped to the MDT created from Std pre-Mn(II) images. See Extended Methods 5.1. Group-wise average Jacobian determinants (JD) were calculated ($JD_E$: ELA; $JD_S$: Std), and then the voxel-wise percent difference between groups determined ($[JD_S - JD_E] \times 100\%$). *(A)* Shown are coronal slices of images color-coded for $[JD_S - JD_E] \times 100\%$. Color gradient represents the percentage difference in the JD between groups and therefore the percentage volume difference, ranging from -15% to +15%. Slices are overlaid on the aligned *InVivo* Atlas for segmentation, with cross-sectional locations indicated above as blue lines in the sagittal slice. JD values represent the contraction or expansion factor needed to align ELA anatomy to that of the Std MDT. We calculated the segment-wise average JD values and performed two-sample t-tests comparing $JD_E$ to $JD_S$. No statistically significant differences between groups (ELA vs Std) were found at our imaging resolution of $100\mu m^3$ across the 117 segments in our atlas ($p < 0.05$, FDR). Using the same segment-wise JD, we compared segment volumes of ELA females ($n = 6$) to those of ELA males ($n = 6$) and similarly found no significant segmental size differences ($p < 0.05$, FDR).

**Fig. S10.**

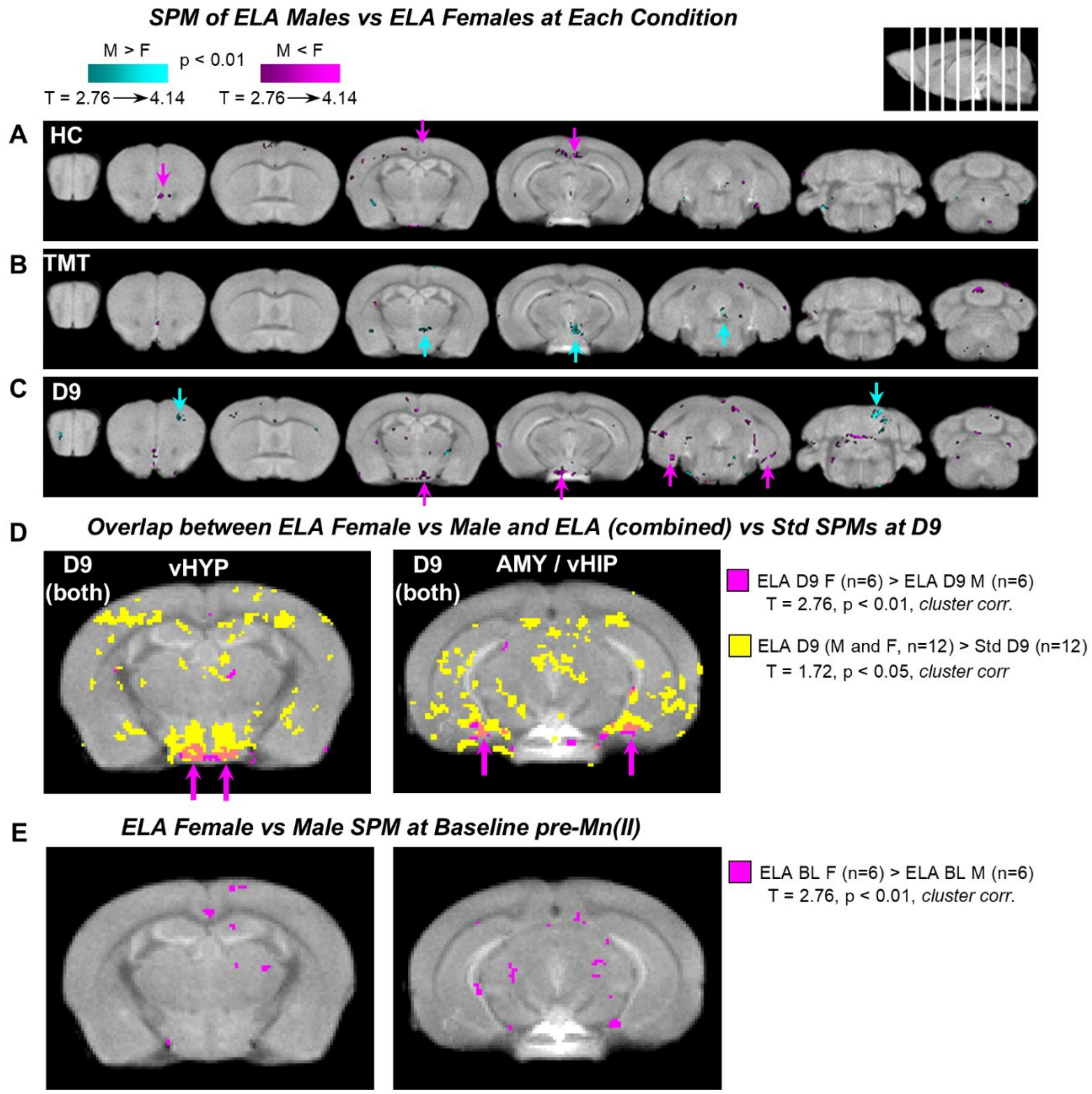


**Fig. S10.** Female Responses Differ Slightly from Male at each Condition.

*(A-C)* Maps from two-sample t-tests comparing ELA females (F) and males (M) at each condition are shown. Statistical maps of M > F (blue) and M < F (pink) are projected onto the dataset average and shown in coronal slices. Coronal slice positions are indicated by white lines in the sagittal slice above. *(D)* Small regions of heightened signal at D9 in ELA females compared to males overlapped with between-group differences in ventral HYP (*left*) and AMY/ventral HIP (*right*). Shown are example slices demonstrating the overlap between the two SPM comparisons projected onto the dataset average: 1) ELA Females > ELA Males at D9 (pink) (n = 6 per group, T = 2.75, p < 0.01, cluster corr.); and 2) ELA Males and Females > Std at D9 (yellow) (n = 12 per group, T = 1.72, p < 0.05, cluster corr.). To assess the degree of overlap, we performed a Jaccard similarity analysis comparing the between-sex with the between-group SPMs at each condition. The Jaccard indices averaged around 0.004 across conditions and only reached as high as 0.025 for the D9 SPMs

shown in *(D)*. This small number of overlapping statistically significant voxels suggested sex-differences were unlikely to contribute to the larger between-group differences. *(E)* An SPM comparison of baseline pre-Mn(II) images between ELA females and males identified only minimal intensity differences. Shown are the same two slices as in *(D)*, of the statistical map for ELA Females > ELA Males at BL pre-Mn(II) (pink) projected onto the dataset average (n = 6 per group, T = 2.75, $p < 0.01$, cluster corr.). Some signal differences between-sexes at baseline corresponded to the anatomy of the posterior cerebral artery. Jaccard similarity analysis of post-Mn(II) versus pre-Mn(II) between-sex SPMs found an average Jaccard index of 0.029. This small value suggested minimal contribution of baseline pre-Mn(II) to post-Mn(II) signal differences for between-sex comparisons.

**Fig. S11.**

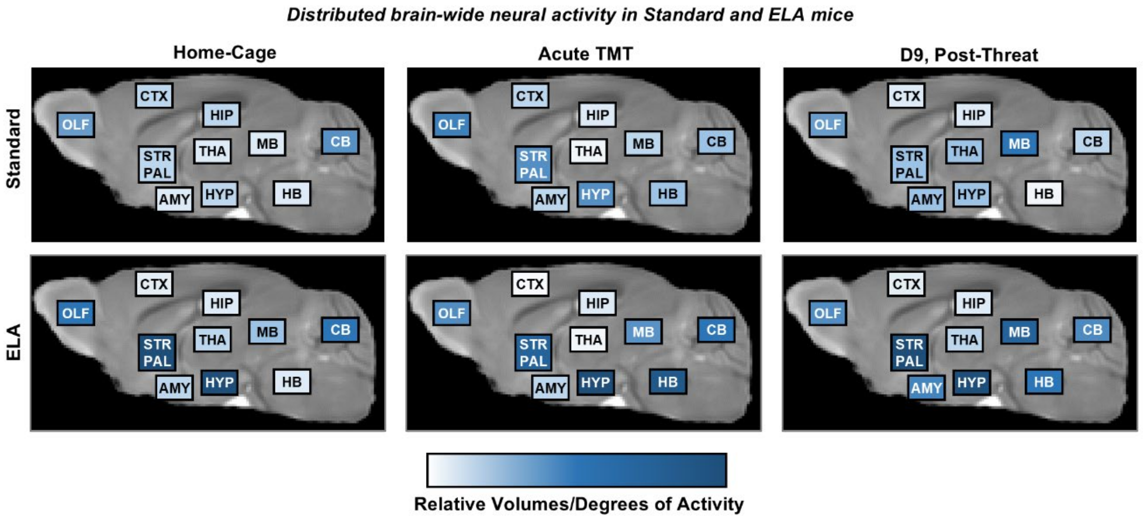


**Fig. S11.** Summary Diagrams of Anatomical Distributions of Neural Activity Patterns Characterizing Brain States of Both Groups at each of the Three Conditions.

Data used for Fig. 5 was diagrammed as in Fig. 6, showing the brain states identified by within-group comparisons of post-Mn(II) > pre-Mn(II); Std (*top*) and ELA (*bottom*); HC (*left*), TMT (*middle*), and D9 (*right*). Rectangular blocks representing the approximate location of 10 anatomical domains as defined by our *InVivo* Atlas were overlaid on a sagittal slice of our grayscale dataset average image. Relative level of activities across domains and between conditions within each group are indicated by the white-to-blue color gradient. Note the relative balance of activity between regions shifts across segments between conditions, even though the anatomical network is likely unchanged.

**SI Tables S1 to S7**

**Table S1.** MRI Hardware and Scan Parameters for ELA and Std Groups

| MRI Hardware and Scan Parameters | | | | | | | | |
|---|---|---|---|---|---|---|---|---|
| **Group** | **Magnet** | **Tesla** | **Bore Size/Type** | **Gradient** | **RF Coil** | **Pulse Sequence** | **TR** | **TE** |
| ELA | Bruker BioSpin Avance DRX500 | 11.7T | 89 mm/vertical | Micro2.5 | 35 mm linear volume coil | 3D-FLASH | 25 ms | 5 ms |
| Standard | Bruker BioSpin Avance DRX500 | 11.7T | 89 mm/vertical | Micro2.5 | 35 mm linear volume coil | 3D-FLASH | 25 ms | 5 ms |
| **Group** | **Flip Angle** | **No. of Averages** | **FOV** | **Voxel Size** | **Scan Time** | **Isoflurane (%)** | **Respiration Rate** | **Body Temp.** |
| ELA | 20° | 8 | 20.0 mm x 12.4 mm x 8.2 mm | 0.1 mm isotropic | 34 min | 1-2% | 100-120 breaths/min | 37°C |
| Standard | 20° | 8 | 20.0 mm x 12.4 mm x 8.2 mm | 0.1 mm isotropic | 34 min | 1-2% | 100-120 breaths/min | 37°C |

**Table S2.** Voxel Coordinates for Regions of Interest (ROI) Analysis

| Region | Hemisphere | Voxel Coordinates | | |
|---|---|---|---|---|
| | | ML | DV | AP |
| MOB-l | Right | 48 | 39 | 49 |
| | Left | 70 | 40 | 49 |
| AON | Right | 46 | 58 | 30 |
| | Left | 73 | 56 | 29 |
| PIR | Right | 22 | 87 | 20 |
| | Left | 90 | 77 | 21 |
| CEA | Right | 37 | 91 | 22 |
| | Left | 86 | 90 | 20 |
| MOB-m | Right | 57 | 46 | 39 |
| | Left | 63 | 46 | 41 |
| NDB | n.a. | 58 | 70 | 26 |
| LSr | Right | 57 | 67 | 29 |
| | Left | 65 | 68 | 30 |
| ACB | Right | 53 | 71 | 26 |
| | Left | 66 | 72 | 26 |

Voxel locations are for centroids of 3x3x3 voxel cubes. Voxel numbers are counted from the right-anterior-ventral edge of MEMRI images' field of view. Voxel coordinates for bregma for this MEMRI data is ML = 65, AP = 89, DV = 80.

**Table S3.** Degree and Statistics of Signal Intensities in Olfactory and Amygdala ROIs of ELA. ROIs correspond to those in **Fig. 3A**. Average and confidence intervals shown as a percentage of pre-Mn(II). All calculations correspond to measurements taken from warped ELA images. FDR adjustments were performed across ROIs but separately for each comparison.

| ROI | Comparison | Average Difference (%) | Standard Deviation (%) | Cohen's D Effect Size | P-value (FDR) |
|---|---|---|---|---|---|
| dlOB-DII | HC > pre-Mn(II) | 6.797 | 4.352 | 1.56 | 0.00036 |
| | TMT> pre-Mn(II) | 10.274 | 4.633 | 2.22 | 0.00000 |
| | TMT > HC | 3.477 | 4.092 | 0.85 | 0.01335 |
| vlAON | HC > pre-Mn(II) | 8.038 | 5.501 | 1.46 | 0.00036 |
| | TMT> pre-Mn(II) | 11.949 | 3.995 | 2.99 | 0.00000 |
| | TMT > HC | 3.910 | 3.326 | 1.18 | 0.00368 |
| aOC | HC > pre-Mn(II) | 1.375 | 3.729 | 0.37 | 0.11397 |
| | TMT> pre-Mn(II) | 3.494 | 2.839 | 1.23 | 0.00067 |
| | TMT > HC | 2.119 | 2.275 | 0.93 | 0.01076 |
| aCEA | HC > pre-Mn(II) | 2.997 | 3.821 | 0.78 | 0.01336 |
| | TMT> pre-Mn(II) | 6.211 | 3.562 | 1.74 | 0.00005 |
| | TMT > HC | 3.213 | 2.689 | 1.20 | 0.00368 |

**Table S4** Degree and statistics of signal intensities between groups in ROIs responsive to TMT in Standard.
ROIs correspond to those in **Fig. 3B**. Average and confidence intervals are shown as a percentage of pre-Mn(II). All calculations correspond to measurements taken from warped images. FDR adjustments were performed across ROIs but separately for each comparison.
* Results from Welch's test, for comparison of heteroscedastic data between groups. Cohen's D and pooled standard deviation from this comparison are referred to as d* and s* in Fig. 3B.

| ROI | Comparison | Average Difference (%) | Pooled Standard Deviation (%) | Cohen's D Effect Size | P-value (FDR) |
|---|---|---|---|---|---|
| mOB-DII | Std HC > pre-Mn(II) | 11.248 | 8.308 | 1.35 | 0.00232 |
| | ELA HC > pre-Mn(II) | 17.894 | 3.459 | 5.17 | 0.00000 |
| | ELA HC ≠ Std HC | 6.646 | *6.364 | *1.04 | *0.02828 |
| dmNDB | Std HC > pre-Mn(II) | 6.129 | 6.391 | 0.96 | 0.00980 |
| | ELA HC > pre-Mn(II) | 11.781 | 4.875 | 2.42 | 0.00000 |
| | ELA HC ≠ Std HC | 5.652 | 5.648 | 1.00 | 0.02828 |
| aLS | Std HC > pre-Mn(II) | 3.642 | 5.260 | 0.69 | 0.02969 |
| | ELA HC > pre-Mn(II) | 7.989 | 3.007 | 2.66 | 0.00000 |
| | ELA HC ≠ Std HC | 4.347 | 4.233 | 1.03 | 0.02828 |
| dmACB | Std HC > pre-Mn(II) | 2.033 | 4.066 | 0.50 | 0.06416 |
| | ELA HC > pre-Mn(II) | 6.138 | 4.005 | 1.53 | 0.00012 |
| | ELA HC ≠ Std HC | 4.105 | 4.034 | 1.02 | 0.02828 |

**Table S5.** Key to *InVivo* Atlas (v10.4) Segmental Abbreviations in Anatomical Order, color-coded per domain, as in the main text figures.

| Domain | Abbreviation | Segment Name |
|---|---|---|
| **Olfactory System: OLF** | | |
| OLF | MOBgl | Main olfactory bulb glomerular layer |
| OLF | MOBgr | Main olfactory bulb granule layer |
| OLF | MOBipl | Main olfactory bulb inner plexiform layer |
| OLF | MOBmi | Main olfactory bulb mitral layer |
| OLF | MOBopl | Main olfactory bulb outer plexiform layer |
| OLF | AOB | Accessory olfactory bulb |
| OLF | AON | Anterior olfactory nucleus |
| OLF | EPd | Endopiriform nucleus dorsal part |
| OLF | TT | Taenia tecta dorsal part |
| OLF | SEZ | Subependymal zone |
| **Cerebral Cortex: CTX** | | |
| CTX | CTX | Cerebral cortex (unsegmented) |
| CTX | ORB | Orbital frontal cortex |
| CTX | PL | Prelimbic cortex |
| CTX | ILA | Infralimbic cortex |
| CTX | DP | Dorsal peduncular cortex |
| CTX | ACA | Anterior cingulate cortex |
| CTX | MO | Primary motor cortex |
| CTX | SS | Somatosensory cortex |
| CTX | PIR | Piriform cortex |
| CTX | RSP | Retrosplenial cortex |
| CTX | PTL | Posterior parietal association area |
| **Hippocampus: HIP** | | |
| HIP | HPF | Hippocampal formation |
| HIP | DG | Dentate gyrus |
| HIP | CA1-CA3 | Field CA1, CA2, CA3 pyramidal layers |
| **Amygdala: AMY** | | |
| AMY | AAA | Anterior amygdala area |
| AMY | MEA | Medial amygdalar nucleus |
| AMY | CEA | Central amygdalar nucleus |
| AMY | LA | Lateral amygdalar nucleus |
| AMY | BLA | Basolateral amygdalar nucleus |
| AMY | PA | Posterior amygdalar nucleus |

| | | |
|---|---|---|
| AMY | COA | Cortical amygdalar area |
| **Striatum and Pallidum: STR/PAL** | | |
| STR/PAL | CP | Caudoputamen |
| STR/PAL | ACB | Nucleus accumbens |
| STR/PAL | FS | Fundus of striatum |
| STR/PAL | OT | Olfactory tubercle |
| STR/PAL | LSc | Lateral septal nucleus caudal part |
| STR/PAL | LSr | Lateral septal nucleus rostral part |
| STR/PAL | GPe | Globus pallidus external |
| STR/PAL | SI | Substantia innominata |
| STR/PAL | MS | Medial septal nucleus |
| STR/PAL | NDB | Diagonal band nucleus |
| STR/PAL | BST | Bed nuclei of the stria terminalis |
| **Thalamus: THA** | | |
| THA | VAL | Ventral anterior lateral complex of the thalamus |
| THA | VM | Ventral medial nucleus of the thalamus |
| THA | VPM | Ventral posteromedial nucleus of the thalamus |
| THA | VPL | Ventral posterolateral nucleus of the thalamus |
| THA | SPA | Subparafascicular area |
| THA | MG | Medial geniculate complex |
| THA | LGd | Dorsal part of the lateral geniculate complex |
| THA | LP | Lateral posterior nucleus of the thalamus |
| THA | AV | Anteroventral nucleus of thalamus |
| THA | IAM | Anteromedial nucleus of thalamus |
| THA | IMD | Intermediodorsal nucleus of the thalamus |
| THA | PVT | Paraventricular nucleus of the thalamus |
| THA | MD | Mediodorsal nucleus of thalamus |
| THA | PT | Parataenial nucleus |
| THA | RE | Nucleus of reunions |
| THA | RT | Reticular nucleus of the thalamus |
| THA | CM | Central medial nucleus of the thalamus |
| THA | PO | Posterior complex of the thalamus |
| THA | PF | Parafascicular nucleus |
| THA | MH | Medial habenula |
| **Hypothalamus: HYP** | | |
| HYP | HY | Hypothalamus (unsegmented) |
| HYP | MPO | Preoptic nuclei |

| | | |
|---|---|---|
| HYP | AVPV | Anteroventral periventricular nucleus |
| HYP | PVH | Paraventricular hypothalamic nucleus |
| HYP | SCH | Suprachiasmatic nucleus |
| HYP | AHN | Anterior hypothalamic nucleus |
| HYP | SBPV | Subparaventricular zone |
| HYP | LHA | Lateral hypothalamic area |
| HYP | ZI | Zona incerta |
| HYP | ARH | Arcuate hypothalamic nucleus |
| HYP | VMH | Ventromedial hypothalamic nucleus |
| HYP | DMH | Dorsomedial hypothalamic nucleus |
| HYP | PVp | Periventricular hypothalamic nucleus (posterior part) |
| HYP | PH | Posterior hypothalamic nucleus |
| HYP | STN | Subthalamic nucleus |
| HYP | MBO | Mammillary nuclei of the hypothalamus |
| **Midbrain: MB** | | |
| MB | MB | Midbrain (unsegmented) |
| MB | VTA | Ventral tegmental area |
| MB | RN | Red nucleus |
| MB | SNc | Substantia nigra compact part |
| MB | SNr | Substantia nigra reticular |
| MB | IPN | Interpeduncular nucleus |
| MB | CLI | Central linear raphe |
| MB | CS | Superior central raphe |
| MB | DR | Dorsal nucleus raphe |
| MB | PAG | Periaqueductal gray |
| **Hindbrain: HB** | | |
| HB | P | Pons |
| HB | PG | Pontine gray |
| HB | PCG | Pontine central gray |
| HB | PRN | Pontine reticular nucleus |
| HB | PB | Parabrachial nucleus |
| HB | LC | Locus coeruleus |
| HB | MY | Medulla oblongata |
| **Cerebellum: CB** | | |
| CB | CB | Cerebellum (unsegmented) |
| CB | DEC | Declive VI |
| CB | FOTU | Folium-tuber vermis VII |

| | | |
|---|---|---|
| CB | PYR | Pyramus VIII |
| CB | NOD | Nodulus X |
| CB | UVU | Uvula IX |
| CB | SIM | Simple lobule |

**Table S6A.** Segment-wise fractional activation volumes (FAV) in Standard and ELA at each condition.

Note that these FAV values are for Post-Mn(II) versus Pre-Mn(II) comparisons in **Fig. 5**. For Post-Mn(II) versus Post-Mn(II) comparisons see **Table S6B**. Columns labeled "HC" are values used for the blue column graphs in **Fig. S7A-B**. Total segment volume is equal to the number of voxels per segment in the corresponding row multiplied by our voxel resolution of 0.1 $mm^3$. In the six columns on the right, the first number is the number of significant voxels and the second number (separated by a "/") is the corresponding fractional activation volume.

| **Domain** | **Segment** | **Total Segment Voxels** | **Significant Voxels / FAV by Condition** | | | | | |
|---|---|---|---|---|---|---|---|---|
| | | | **Standard** | | | **ELA** | | |
| | | | **HC** | **TMT** | **D9** | **HC** | **TMT** | **D9** |
| OLF | MOBgl | 6904 | 3174 / 0.46 | 3373 / 0.49 | 3072 / 0.44 | 3906 / 0.57 | 3703 / 0.54 | 3173 / 0.46 |
| | MOBgr | 4577 | 1597 / 0.35 | 1842 / 0.4 | 1380 / 0.3 | 2584 / 0.56 | 2579 / 0.56 | 1755 / 0.38 |
| | MOBipl | 4912 | 3629 / 0.74 | 3373 / 0.69 | 3429 / 0.7 | 4264 / 0.87 | 4022 / 0.82 | 4054 / 0.83 |
| | MOBmi | 1064 | 848 / 0.8 | 848 / 0.8 | 879 / 0.83 | 975 / 0.92 | 949 / 0.89 | 966 / 0.91 |
| | MOBopl | 1234 | 708 / 0.57 | 765 / 0.62 | 732 / 0.59 | 841 / 0.68 | 818 / 0.66 | 771 / 0.62 |
| | AOB | 599 | 171 / 0.29 | 93 / 0.16 | 132 / 0.22 | 244 / 0.41 | 182 / 0.3 | 164 / 0.27 |
| | AON | 3084 | 743 / 0.24 | 1278 / 0.41 | 1161 / 0.38 | 1451 / 0.47 | 1542 / 0.5 | 1372 / 0.44 |
| | TT | 1097 | 23 / 0.02 | 113 / 0.1 | 22 / 0.02 | 121 / 0.11 | 110 / 0.1 | 134 / 0.12 |
| | EPd | 910 | 128 / 0.14 | 211 / 0.23 | 231 / 0.25 | 241 / 0.26 | 252 / 0.28 | 235 / 0.26 |
| CTX | CTX | 48931 | 234 / 0 | 236 / 0 | 270 / 0.01 | 217 / 0 | 505 / 0.01 | 392 / 0.01 |
| | ORB | 2379 | 0 / 0 | 0 / 0 | 0 / 0 | 2 / 0 | 2 / 0 | 1 / 0 |
| | PL | 1906 | 0 / 0 | 0 / 0 | 0 / 0 | 0 / 0 | 5 / 0 | 2 / 0 |
| | ILA | 1371 | 0 / 0 | 7 / 0.01 | 0 / 0 | 41 / 0.03 | 53 / 0.04 | 59 / 0.04 |
| | DP | 454 | 0 / 0 | 72 / 0.16 | 0 / 0 | 71 / 0.16 | 40 / 0.09 | 39 / 0.09 |
| | ACA | 5311 | 40 / 0.01 | 0 / 0 | 7 / 0 | 119 / 0.02 | 60 / 0.01 | 40 / 0.01 |
| | MO | 16452 | 108 / 0.01 | 65 / 0 | 35 / 0 | 31 / 0 | 107 / 0.01 | 95 / 0.01 |
| | SS | 30181 | 88 / 0 | 128 / 0 | 48 / 0 | 184 / 0.01 | 245 / 0.01 | 194 / 0.01 |
| | PIR | 9948 | 939 / 0.09 | 1435 / 0.14 | 1731 / 0.17 | 1920 / 0.19 | 2264 / 0.23 | 1770 / 0.18 |
| | RSP | 6191 | 8 / 0 | 9 / 0 | 0 / 0 | 2 / 0 | 0 / 0 | 0 / 0 |
| | PTL | 2826 | 0 / 0 | 0 / 0 | 0 / 0 | 0 / 0 | 0 / 0 | 0 / 0 |
| HIP | HPF | 10162 | 773 / 0.08 | 602 / 0.06 | 691 / 0.07 | 989 / 0.1 | 494 / 0.05 | 948 / 0.09 |
| | DG | 2501 | 1401 / 0.56 | 1165 / 0.47 | 1440 / 0.58 | 1348 / 0.54 | 1173 / 0.47 | 1469 / 0.59 |
| | CA1-CA3 | 12826 | 2195 / 0.17 | 1800 / 0.14 | 2180 / 0.17 | 3172 / 0.25 | 2227 / 0.17 | 3737 / 0.29 |
| AMY | AAA | 667 | 0 / 0 | 18 / 0.03 | 154 / 0.23 | 74 / 0.11 | 102 / 0.15 | 155 / 0.23 |
| | MEA | 1092 | 164 / 0.15 | 116 / 0.11 | 692 / 0.63 | 351 / 0.32 | 433 / 0.4 | 743 / 0.68 |
| | CEA | 1578 | 397 / 0.25 | 297 / 0.19 | 1190 / 0.75 | 682 / 0.43 | 830 / 0.53 | 1319 / 0.84 |
| | LA | 1261 | 230 / 0.18 | 99 / 0.08 | 198 / 0.16 | 182 / 0.14 | 63 / 0.05 | 201 / 0.16 |

| | | | | | | | | |
|---|---|---|---|---|---|---|---|---|
| | BLA | 3253 | 196 / 0.06 | 142 / 0.04 | 592 / 0.18 | 228 / 0.07 | 169 / 0.05 | 564 / 0.17 |
| | PA | 463 | 117 / 0.25 | 123 / 0.27 | 98 / 0.21 | 161 / 0.35 | 142 / 0.31 | 249 / 0.54 |
| | COA | 2626 | 133 / 0.05 | 207 / 0.08 | 342 / 0.13 | 313 / 0.12 | 252 / 0.1 | 703 / 0.27 |
| STR/PAL | CP | 22436 | 330 / 0.01 | 344 / 0.02 | 1244 / 0.06 | 1205 / 0.05 | 948 / 0.04 | 1706 / 0.08 |
| | ACB | 5108 | 1081 / 0.21 | 2151 / 0.42 | 2733 / 0.54 | 2658 / 0.52 | 2817 / 0.55 | 2940 / 0.58 |
| | FS | 570 | 43 / 0.08 | 61 / 0.11 | 418 / 0.73 | 281 / 0.49 | 344 / 0.6 | 443 / 0.78 |
| | OT | 2551 | 49 / 0.02 | 137 / 0.05 | 121 / 0.05 | 163 / 0.06 | 178 / 0.07 | 184 / 0.07 |
| | LSc | 338 | 31 / 0.09 | 51 / 0.15 | 21 / 0.06 | 208 / 0.62 | 115 / 0.34 | 75 / 0.22 |
| | LSr | 1952 | 401 / 0.21 | 916 / 0.47 | 1003 / 0.51 | 1596 / 0.82 | 1246 / 0.64 | 1237 / 0.63 |
| | GPe | 1678 | 783 / 0.47 | 666 / 0.4 | 1244 / 0.74 | 1236 / 0.74 | 1340 / 0.8 | 1438 / 0.86 |
| | SI | 2927 | 469 / 0.16 | 708 / 0.24 | 2040 / 0.7 | 2131 / 0.73 | 2179 / 0.74 | 2173 / 0.74 |
| | MS | 220 | 182 / 0.83 | 197 / 0.9 | 215 / 0.98 | 220 / 1 | 218 / 0.99 | 219 / 1 |
| | NDB | 661 | 276 / 0.42 | 363 / 0.55 | 467 / 0.71 | 468 / 0.71 | 478 / 0.72 | 445 / 0.67 |
| | BST | 1800 | 687 / 0.38 | 793 / 0.44 | 1642 / 0.91 | 1657 / 0.92 | 1597 / 0.89 | 1750 / 0.97 |
| THA | VAL | 413 | 12 / 0.03 | 1 / 0 | 13 / 0.03 | 17 / 0.04 | 8 / 0.02 | 26 / 0.06 |
| | VM | 936 | 16 / 0.02 | 3 / 0 | 46 / 0.05 | 8 / 0.01 | 39 / 0.04 | 45 / 0.05 |
| | VPM | 1020 | 16 / 0.02 | 1 / 0 | 78 / 0.08 | 22 / 0.02 | 4 / 0 | 43 / 0.04 |
| | VPL | 1298 | 36 / 0.03 | 8 / 0.01 | 87 / 0.07 | 85 / 0.07 | 53 / 0.04 | 100 / 0.08 |
| | SPA | 202 | 0 / 0 | 0 / 0 | 18 / 0.09 | 2 / 0.01 | 1 / 0 | 32 / 0.16 |
| | MG | 1195 | 21 / 0.02 | 0 / 0 | 25 / 0.02 | 40 / 0.03 | 15 / 0.01 | 82 / 0.07 |
| | LGd | 53 | 4 / 0.08 | 6 / 0.11 | 22 / 0.42 | 28 / 0.53 | 0 / 0 | 34 / 0.64 |
| | LP | 1353 | 27 / 0.02 | 6 / 0 | 109 / 0.08 | 45 / 0.03 | 0 / 0 | 158 / 0.12 |
| | AV | 194 | 25 / 0.13 | 0 / 0 | 4 / 0.02 | 82 / 0.42 | 14 / 0.07 | 38 / 0.2 |
| | IAM | 507 | 13 / 0.03 | 1 / 0 | 57 / 0.11 | 59 / 0.12 | 10 / 0.02 | 102 / 0.2 |
| | IMD | 65 | 0 / 0 | 0 / 0 | 0 / 0 | 0 / 0 | 0 / 0 | 1 / 0.02 |
| | PVT | 1349 | 64 / 0.05 | 8 / 0.01 | 360 / 0.27 | 194 / 0.14 | 164 / 0.12 | 507 / 0.38 |
| | MD | 725 | 0 / 0 | 0 / 0 | 8 / 0.01 | 2 / 0 | 0 / 0 | 15 / 0.02 |
| | PT | 343 | 1 / 0 | 0 / 0 | 1 / 0 | 2 / 0.01 | 1 / 0 | 8 / 0.02 |
| | RE | 376 | 12 / 0.03 | 5 / 0.01 | 73 / 0.19 | 27 / 0.07 | 74 / 0.2 | 160 / 0.43 |
| | RT | 2556 | 606 / 0.24 | 351 / 0.14 | 788 / 0.31 | 1202 / 0.47 | 868 / 0.34 | 1218 / 0.48 |
| | CM | 159 | 11 / 0.07 | 0 / 0 | 35 / 0.22 | 5 / 0.03 | 21 / 0.13 | 22 / 0.14 |
| | PO | 1242 | 0 / 0 | 0 / 0 | 8 / 0.01 | 0 / 0 | 0 / 0 | 2 / 0 |
| | PF | 606 | 0 / 0 | 3 / 0 | 16 / 0.03 | 4 / 0.01 | 0 / 0 | 37 / 0.06 |
| | MH | 212 | 76 / 0.36 | 58 / 0.27 | 139 / 0.66 | 161 / 0.76 | 88 / 0.42 | 191 / 0.9 |
| HYP | HY | 1075 | 107 / 0.1 | 132 / 0.12 | 294 / 0.27 | 407 / 0.38 | 439 / 0.41 | 527 / 0.49 |
| | MPO | 1334 | 306 / 0.23 | 426 / 0.32 | 1029 / 0.77 | 1216 / 0.91 | 1211 / 0.91 | 1150 / 0.86 |
| | AVPV | 82 | 8 / 0.1 | 1 / 0.01 | 50 / 0.61 | 63 / 0.77 | 64 / 0.78 | 62 / 0.76 |
| | PVH | 196 | 60 / 0.31 | 29 / 0.15 | 130 / 0.66 | 108 / 0.55 | 111 / 0.57 | 170 / 0.87 |
| | SCH | 115 | 33 / 0.29 | 22 / 0.19 | 47 / 0.41 | 83 / 0.72 | 88 / 0.77 | 65 / 0.57 |

| | | | | | | | | |
|---|---|---|---|---|---|---|---|---|
| | AHN | 517 | 361 / 0.7 | 335 / 0.65 | 497 / 0.96 | 499 / 0.97 | 509 / 0.98 | 516 / 1 |
| | SBPV | 199 | 75 / 0.38 | 50 / 0.25 | 120 / 0.6 | 104 / 0.52 | 111 / 0.56 | 151 / 0.76 |
| | LHA | 2478 | 931 / 0.38 | 561 / 0.23 | 1871 / 0.76 | 2142 / 0.86 | 2223 / 0.9 | 2306 / 0.93 |
| | ZI | 1764 | 205 / 0.12 | 83 / 0.05 | 772 / 0.44 | 443 / 0.25 | 551 / 0.31 | 1007 / 0.57 |
| | ARH | 152 | 0 / 0 | 6 / 0.04 | 1 / 0.01 | 0 / 0 | 2 / 0.01 | 16 / 0.11 |
| | VMH | 368 | 73 / 0.2 | 207 / 0.56 | 230 / 0.63 | 313 / 0.85 | 311 / 0.85 | 346 / 0.94 |
| | DMH | 390 | 60 / 0.15 | 31 / 0.08 | 216 / 0.55 | 288 / 0.74 | 307 / 0.79 | 360 / 0.92 |
| | PVp | 243 | 4 / 0.02 | 23 / 0.09 | 15 / 0.06 | 19 / 0.08 | 10 / 0.04 | 30 / 0.12 |
| | PH | 570 | 59 / 0.1 | 1 / 0 | 422 / 0.74 | 313 / 0.55 | 393 / 0.69 | 503 / 0.88 |
| | STN | 271 | 83 / 0.31 | 69 / 0.25 | 243 / 0.9 | 215 / 0.79 | 200 / 0.74 | 260 / 0.96 |
| | MBO | 892 | 40 / 0.04 | 15 / 0.02 | 285 / 0.32 | 275 / 0.31 | 388 / 0.43 | 357 / 0.4 |
| MB | MB | 19996 | 194 / 0.01 | 430 / 0.02 | 1993 / 0.1 | 297 / 0.01 | 986 / 0.05 | 2172 / 0.11 |
| | VTA | 764 | 157 / 0.21 | 162 / 0.21 | 516 / 0.68 | 246 / 0.32 | 274 / 0.36 | 499 / 0.65 |
| | RN | 344 | 14 / 0.04 | 1 / 0 | 105 / 0.31 | 0 / 0 | 8 / 0.02 | 23 / 0.07 |
| | SNc | 724 | 67 / 0.09 | 61 / 0.08 | 283 / 0.39 | 91 / 0.13 | 115 / 0.16 | 423 / 0.58 |
| | SNr | 1165 | 195 / 0.17 | 253 / 0.22 | 303 / 0.26 | 185 / 0.16 | 139 / 0.12 | 441 / 0.38 |
| | IPN | 281 | 87 / 0.31 | 165 / 0.59 | 181 / 0.64 | 187 / 0.67 | 196 / 0.7 | 181 / 0.64 |
| | CLI | 132 | 1 / 0.01 | 4 / 0.03 | 46 / 0.35 | 5 / 0.04 | 12 / 0.09 | 18 / 0.14 |
| | CS | 597 | 46 / 0.08 | 87 / 0.15 | 206 / 0.35 | 176 / 0.29 | 236 / 0.4 | 319 / 0.53 |
| | DR | 175 | 3 / 0.02 | 14 / 0.08 | 56 / 0.32 | 26 / 0.15 | 29 / 0.17 | 36 / 0.21 |
| | PAG | 4263 | 22 / 0.01 | 69 / 0.02 | 427 / 0.1 | 61 / 0.01 | 104 / 0.02 | 703 / 0.16 |
| HB | P | 5550 | 536 / 0.1 | 817 / 0.15 | 141 / 0.03 | 713 / 0.13 | 1480 / 0.27 | 270 / 0.05 |
| | PG | 712 | 0 / 0 | 0 / 0 | 3 / 0 | 7 / 0.01 | 4 / 0.01 | 2 / 0 |
| | PCG | 803 | 119 / 0.15 | 133 / 0.17 | 77 / 0.1 | 153 / 0.19 | 351 / 0.44 | 304 / 0.38 |
| | PRN | 5542 | 201 / 0.04 | 237 / 0.04 | 287 / 0.05 | 114 / 0.02 | 342 / 0.06 | 474 / 0.09 |
| | PB | 1151 | 61 / 0.05 | 111 / 0.1 | 101 / 0.09 | 52 / 0.05 | 222 / 0.19 | 151 / 0.13 |
| | LC | 96 | 7 / 0.07 | 3 / 0.03 | 11 / 0.11 | 0 / 0 | 40 / 0.42 | 33 / 0.34 |
| | MY | 28535 | 3114 / 0.11 | 2192 / 0.08 | 1049 / 0.04 | 4521 / 0.16 | 7071 / 0.25 | 1146 / 0.04 |
| CB | CB | 58909 | 8613 / 0.15 | 5866 / 0.1 | 3355 / 0.06 | 11157 / 0.19 | 13661 / 0.23 | 3330 / 0.06 |
| | DEC | 960 | 1 / 0 | 0 / 0 | 0 / 0 | 0 / 0 | 0 / 0 | 0 / 0 |
| | FOTU | 226 | 0 / 0 | 0 / 0 | 0 / 0 | 0 / 0 | 0 / 0 | 0 / 0 |
| | PYR | 609 | 394 / 0.65 | 86 / 0.14 | 9 / 0.01 | 375 / 0.62 | 438 / 0.72 | 79 / 0.13 |
| | NOD | 556 | 523 / 0.94 | 452 / 0.81 | 336 / 0.6 | 526 / 0.95 | 541 / 0.97 | 407 / 0.73 |
| | UVU | 1888 | 1457 / 0.77 | 764 / 0.4 | 513 / 0.27 | 1509 / 0.8 | 1650 / 0.87 | 353 / 0.19 |
| | SIM | 236 | 0 / 0 | 0 / 0 | 0 / 0 | 0 / 0 | 0 / 0 | 0 / 0 |

**Table S6B.** Segment-wise fractional volumes from comparisons between post-Mn(II) images in Standard and ELA mice.

Data in columns labeled HC > TMT, HC < TMT, HC > D9 and HC < D9 correspond to **Fig. S7C-F**. Total segment volume is equal to the number of voxels per segment multiplied by the voxel resolution of 0.1 $mm^3$. In the comparison columns, the first value is the number of significant voxels and the second value (separated by a "/") is the corresponding fractional volume. Note: values are statistical results from t-tests that compared signal intensities directly between post-Mn(II) images.

| Domain | Segment | Total Segment Voxels | Significant Voxels by Comparison | | | | | | | | | | | |
|---|---|---|---|---|---|---|---|---|---|---|---|---|---|---|
| | | | Standard | | | | | | ELA | | | | | |
| | | | HC > TMT | HC < TMT | TMT > D9 | TMT < D9 | HC > D9 | HC < D9 | HC > TMT | HC < TMT | TMT > D9 | TMT < D9 | HC > D9 | HC < D9 |
| OLF | MOBgl | 6904 | 420 / 0.06 | 690 / 0.1 | 508 / 0.07 | 400 / 0.06 | 208 / 0.03 | 143 / 0.02 | 830 / 0.12 | 262 / 0.04 | 735 / 0.11 | 347 / 0.05 | 1410 / 0.2 | 148 / 0.02 |
| | MOBgr | 4577 | 12 / 0 | 266 / 0.06 | 646 / 0.14 | 13 / 0 | 250 / 0.05 | 26 / 0.01 | 165 / 0.04 | 91 / 0.02 | 219 / 0.05 | 3 / 0 | 359 / 0.08 | 4 / 0 |
| | MOBipl | 4912 | 148 / 0.03 | 328 / 0.07 | 148 / 0.03 | 78 / 0.02 | 48 / 0.01 | 109 / 0.02 | 515 / 0.1 | 43 / 0.01 | 140 / 0.03 | 98 / 0.02 | 572 / 0.12 | 6 / 0 |
| | MOBmi | 1064 | 60 / 0.06 | 47 / 0.04 | 7 / 0.01 | 36 / 0.03 | 1 / 0 | 15 / 0.01 | 157 / 0.15 | 4 / 0 | 2 / 0 | 24 / 0.02 | 136 / 0.13 | 6 / 0.01 |
| | MOBopl | 1234 | 80 / 0.06 | 52 / 0.04 | 26 / 0.02 | 82 / 0.07 | 9 / 0.01 | 31 / 0.03 | 247 / 0.2 | 9 / 0.01 | 42 / 0.03 | 65 / 0.05 | 191 / 0.15 | 8 / 0.01 |
| | AOB | 599 | 110 / 0.18 | 4 / 0.01 | 1 / 0 | 8 / 0.01 | 63 / 0.11 | 0 / 0 | 123 / 0.21 | 0 / 0 | 4 / 0.01 | 0 / 0 | 178 / 0.3 | 0 / 0 |
| | AON | 3084 | 0 / 0 | 521 / 0.17 | 79 / 0.03 | 18 / 0.01 | 0 / 0 | 305 / 0.1 | 79 / 0.03 | 69 / 0.02 | 33 / 0.01 | 27 / 0.01 | 10 / 0 | 15 / 0 |
| | TT | 1097 | 0 / 0 | 68 / 0.06 | 109 / 0.1 | 0 / 0 | 30 / 0.03 | 23 / 0.02 | 58 / 0.05 | 0 / 0 | 0 / 0 | 89 / 0.08 | 12 / 0.01 | 14 / 0.01 |
| | EPd | 910 | 0 / 0 | 109 / 0.12 | 52 / 0.06 | 16 / 0.02 | 0 / 0 | 106 / 0.12 | 99 / 0.11 | 2 / 0 | 1 / 0 | 87 / 0.1 | 1 / 0 | 6 / 0.01 |
| CTX | CTX | 48931 | 1207 / 0.02 | 1197 / 0.02 | 3877 / 0.08 | 1080 / 0.02 | 4953 / 0.1 | 1889 / 0.04 | 1893 / 0.04 | 945 / 0.02 | 2418 / 0.05 | 3169 / 0.06 | 2672 / 0.05 | 951 / 0.02 |
| | ORB | 2379 | 5 / 0 | 87 / 0.04 | 336 / 0.14 | 4 / 0 | 147 / 0.06 | 5 / 0 | 139 / 0.06 | 12 / 0.01 | 24 / 0.01 | 10 / 0 | 109 / 0.05 | 3 / 0 |
| | PL | 1906 | 126 / 0.07 | 4 / 0 | 14 / 0.01 | 53 / 0.03 | 36 / 0.02 | 2 / 0 | 124 / 0.07 | 5 / 0 | 25 / 0.01 | 25 / 0.01 | 44 / 0.02 | 0 / 0 |
| | ILA | 1371 | 4 / 0 | 49 / 0.04 | 140 / 0.1 | 3 / 0 | 116 / 0.08 | 8 / 0.01 | 54 / 0.04 | 2 / 0 | 4 / 0 | 100 / 0.07 | 8 / 0.01 | 16 / 0.01 |
| | DP | 454 | 0 / 0 | 62 / 0.14 | 93 / 0.2 | 0 / 0 | 32 / 0.07 | 0 / 0 | 58 / 0.13 | 0 / 0 | 0 / 0 | 20 / 0.04 | 20 / 0.04 | 0 / 0 |
| | ACA | 5311 | 488 / 0.09 | 34 / 0.01 | 356 / 0.07 | 304 / 0.06 | 384 / 0.07 | 28 / 0.01 | 522 / 0.1 | 10 / 0 | 117 / 0.02 | 123 / 0.02 | 287 / 0.05 | 0 / 0 |
| | MO | 16452 | 1579 / 0.1 | 144 / 0.01 | 915 / 0.06 | 523 / 0.03 | 1140 / 0.07 | 13 / 0 | 1288 / 0.08 | 224 / 0.01 | 257 / 0.02 | 244 / 0.01 | 684 / 0.04 | 104 / 0.01 |
| | SS | 30181 | 2547 / 0.08 | 161 / 0.01 | 858 / 0.03 | 1486 / 0.05 | 2432 / 0.08 | 497 / 0.02 | 2475 / 0.08 | 214 / 0.01 | 672 / 0.02 | 3621 / 0.12 | 735 / 0.02 | 744 / 0.02 |
| | PIR | 9948 | 16 / 0 | 1170 / 0.12 | 381 / 0.04 | 706 / 0.07 | 94 / 0.01 | 2317 / 0.23 | 304 / 0.03 | 318 / 0.03 | 357 / 0.04 | 865 / 0.09 | 103 / 0.01 | 92 / 0.01 |
| | RSP | 6191 | 290 / 0.05 | 45 / 0.01 | 342 / 0.06 | 78 / 0.01 | 915 / 0.15 | 30 / 0 | 275 / 0.04 | 108 / 0.02 | 249 / 0.04 | 329 / 0.05 | 266 / 0.04 | 38 / 0.01 |
| | PTL | 2826 | 401 / 0.14 | 0 / 0 | 77 / 0.03 | 88 / 0.03 | 588 / 0.21 | 8 / 0 | 41 / 0.01 | 19 / 0.01 | 48 / 0.02 | 23 / 0.01 | 94 / 0.03 | 0 / 0 |
| HIP | HPF | 10162 | 359 / 0.04 | 113 / 0.01 | 1546 / 0.15 | 347 / 0.03 | 1886 / 0.19 | 290 / 0.03 | 860 / 0.08 | 63 / 0.01 | 533 / 0.05 | 1387 / 0.14 | 473 / 0.05 | 417 / 0.04 |
| | DG | 2501 | 72 / 0.03 | 12 / 0 | 252 / 0.1 | 434 / 0.17 | 261 / 0.1 | 337 / 0.13 | 100 / 0.04 | 3 / 0 | 24 / 0.01 | 549 / 0.22 | 14 / 0.01 | 474 / 0.19 |
| | CA1-CA3 | 12826 | 689 / 0.05 | 91 / 0.01 | 1290 / 0.1 | 784 / 0.06 | 1456 / 0.11 | 185 / 0.01 | 961 / 0.07 | 122 / 0.01 | 780 / 0.06 | 2026 / 0.16 | 642 / 0.05 | 900 / 0.07 |
| AMY | AAA | 667 | 0 / 0 | 35 / 0.05 | 5 / 0.01 | 173 / 0.26 | 0 / 0 | 280 / 0.42 | 8 / 0.01 | 25 / 0.04 | 0 / 0 | 43 / 0.06 | 0 / 0 | 70 / 0.1 |
| | MEA | 1092 | 1 / 0 | 42 / 0.04 | 2 / 0 | 629 / 0.58 | 2 / 0 | 619 / 0.57 | 45 / 0.04 | 8 / 0.01 | 4 / 0 | 421 / 0.39 | 0 / 0 | 402 / 0.37 |
| | CEA | 1578 | 51 / 0.03 | 5 / 0 | 1 / 0 | 1142 / 0.72 | 10 / 0.01 | 994 / 0.63 | 7 / 0 | 61 / 0.04 | 0 / 0 | 1004 / 0.64 | 0 / 0 | 1105 / 0.7 |
| | LA | 1261 | 9 / 0.01 | 12 / 0.01 | 1 / 0 | 181 / 0.14 | 25 / 0.02 | 75 / 0.06 | 49 / 0.04 | 0 / 0 | 0 / 0 | 127 / 0.1 | 0 / 0 | 38 / 0.03 |
| | BLA | 3253 | 9 / 0 | 27 / 0.01 | 46 / 0.01 | 989 / 0.3 | 46 / 0.01 | 1117 / 0.34 | 148 / 0.05 | 8 / 0 | 7 / 0 | 747 / 0.23 | 0 / 0 | 376 / 0.12 |
| | PA | 463 | 0 / 0 | 9 / 0.02 | 66 / 0.14 | 9 / 0.02 | 60 / 0.13 | 9 / 0.02 | 9 / 0.02 | 0 / 0 | 0 / 0 | 46 / 0.1 | 0 / 0 | 70 / 0.15 |

| | | | | | | | | | | | | | | |
|---|---|---|---|---|---|---|---|---|---|---|---|---|---|---|
| | COA | 2626 | 29 / 0.01 | 266 / 0.1 | 177 / 0.07 | 350 / 0.13 | 96 / 0.04 | 622 / 0.24 | 145 / 0.06 | 54 / 0.02 | 79 / 0.03 | 625 / 0.24 | 0 / 0 | 83 / 0.03 |
| STR/PAL | CP | 22436 | 97 / 0 | 1581 / 0.07 | 783 / 0.03 | 1767 / 0.08 | 415 / 0.02 | 3523 / 0.16 | 2505 / 0.11 | 9 / 0 | 58 / 0 | 1991 / 0.09 | 533 / 0.02 | 534 / 0.02 |
| | ACB | 5108 | 1 / 0 | 972 / 0.19 | 68 / 0.01 | 1289 / 0.25 | 0 / 0 | 2664 / 0.52 | 132 / 0.03 | 74 / 0.01 | 5 / 0 | 1187 / 0.23 | 3 / 0 | 1102 / 0.22 |
| | FS | 570 | 0 / 0 | 10 / 0.02 | 0 / 0 | 364 / 0.64 | 0 / 0 | 429 / 0.75 | 0 / 0 | 11 / 0.02 | 0 / 0 | 258 / 0.45 | 0 / 0 | 323 / 0.57 |
| | OT | 2551 | 21 / 0.01 | 369 / 0.14 | 231 / 0.09 | 128 / 0.05 | 23 / 0.01 | 228 / 0.09 | 131 / 0.05 | 26 / 0.01 | 17 / 0.01 | 166 / 0.07 | 21 / 0.01 | 67 / 0.03 |
| | LSc | 338 | 0 / 0 | 12 / 0.04 | 60 / 0.18 | 0 / 0 | 23 / 0.07 | 1 / 0 | 101 / 0.3 | 0 / 0 | 0 / 0 | 23 / 0.07 | 2 / 0.01 | 0 / 0 |
| | LSr | 1952 | 4 / 0 | 246 / 0.13 | 36 / 0.02 | 201 / 0.1 | 13 / 0.01 | 479 / 0.25 | 449 / 0.23 | 0 / 0 | 0 / 0 | 393 / 0.2 | 2 / 0 | 199 / 0.1 |
| | GPe | 1678 | 28 / 0.02 | 38 / 0.02 | 1 / 0 | 870 / 0.52 | 0 / 0 | 668 / 0.4 | 27 / 0.02 | 106 / 0.06 | 17 / 0.01 | 453 / 0.27 | 4 / 0 | 510 / 0.3 |
| | SI | 2927 | 10 / 0 | 435 / 0.15 | 24 / 0.01 | 875 / 0.3 | 0 / 0 | 1442 / 0.49 | 68 / 0.02 | 25 / 0.01 | 6 / 0 | 689 / 0.24 | 1 / 0 | 628 / 0.21 |
| | MS | 220 | 0 / 0 | 23 / 0.1 | 1 / 0 | 20 / 0.09 | 0 / 0 | 18 / 0.08 | 31 / 0.14 | 0 / 0 | 0 / 0 | 30 / 0.14 | 0 / 0 | 19 / 0.09 |
| | NDB | 661 | 0 / 0 | 74 / 0.11 | 0 / 0 | 33 / 0.05 | 0 / 0 | 181 / 0.27 | 18 / 0.03 | 0 / 0 | 11 / 0.02 | 18 / 0.03 | 18 / 0.03 | 1 / 0 |
| | BST | 1800 | 16 / 0.01 | 126 / 0.07 | 3 / 0 | 1224 / 0.68 | 0 / 0 | 1380 / 0.77 | 149 / 0.08 | 5 / 0 | 0 / 0 | 1446 / 0.8 | 0 / 0 | 1289 / 0.72 |
| | VAL | 413 | 63 / 0.15 | 0 / 0 | 0 / 0 | 103 / 0.25 | 0 / 0 | 17 / 0.04 | 29 / 0.07 | 0 / 0 | 0 / 0 | 40 / 0.1 | 0 / 0 | 14 / 0.03 |
| | VM | 936 | 137 / 0.15 | 0 / 0 | 0 / 0 | 494 / 0.53 | 0 / 0 | 77 / 0.08 | 30 / 0.03 | 2 / 0 | 15 / 0.02 | 46 / 0.05 | 0 / 0 | 8 / 0.01 |
| | VPM | 1020 | 25 / 0.02 | 3 / 0 | 0 / 0 | 277 / 0.27 | 3 / 0 | 315 / 0.31 | 148 / 0.15 | 0 / 0 | 1 / 0 | 235 / 0.23 | 0 / 0 | 43 / 0.04 |
| | VPL | 1298 | 64 / 0.05 | 0 / 0 | 0 / 0 | 389 / 0.3 | 0 / 0 | 146 / 0.11 | 149 / 0.11 | 1 / 0 | 6 / 0 | 175 / 0.13 | 4 / 0 | 10 / 0.01 |
| | SPA | 202 | 0 / 0 | 6 / 0.03 | 0 / 0 | 56 / 0.28 | 0 / 0 | 127 / 0.63 | 42 / 0.21 | 0 / 0 | 0 / 0 | 161 / 0.8 | 0 / 0 | 60 / 0.3 |
| | MG | 1195 | 76 / 0.06 | 14 / 0.01 | 4 / 0 | 123 / 0.1 | 50 / 0.04 | 238 / 0.2 | 86 / 0.07 | 25 / 0.02 | 7 / 0.01 | 219 / 0.18 | 17 / 0.01 | 175 / 0.15 |
| | LGd | 53 | 0 / 0 | 0 / 0 | 0 / 0 | 3 / 0.06 | 0 / 0 | 13 / 0.25 | 27 / 0.51 | 0 / 0 | 0 / 0 | 14 / 0.26 | 1 / 0.02 | 0 / 0 |
| | LP | 1353 | 63 / 0.05 | 0 / 0 | 0 / 0 | 192 / 0.14 | 15 / 0.01 | 107 / 0.08 | 182 / 0.13 | 0 / 0 | 0 / 0 | 396 / 0.29 | 0 / 0 | 54 / 0.04 |
| | AV | 194 | 2 / 0.01 | 0 / 0 | 0 / 0 | 9 / 0.05 | 0 / 0 | 0 / 0 | 66 / 0.34 | 0 / 0 | 0 / 0 | 31 / 0.16 | 1 / 0.01 | 8 / 0.04 |
| THA | IAM | 507 | 63 / 0.12 | 0 / 0 | 0 / 0 | 164 / 0.32 | 0 / 0 | 47 / 0.09 | 70 / 0.14 | 1 / 0 | 0 / 0 | 118 / 0.23 | 0 / 0 | 8 / 0.02 |
| | IMD | 65 | 9 / 0.14 | 0 / 0 | 0 / 0 | 36 / 0.55 | 0 / 0 | 23 / 0.35 | 9 / 0.14 | 0 / 0 | 0 / 0 | 30 / 0.46 | 0 / 0 | 17 / 0.26 |
| | PVT | 1349 | 131 / 0.1 | 10 / 0.01 | 1 / 0 | 710 / 0.53 | 0 / 0 | 473 / 0.35 | 195 / 0.14 | 9 / 0.01 | 0 / 0 | 765 / 0.57 | 0 / 0 | 496 / 0.37 |
| | MD | 725 | 35 / 0.05 | 0 / 0 | 0 / 0 | 213 / 0.29 | 0 / 0 | 121 / 0.17 | 129 / 0.18 | 0 / 0 | 0 / 0 | 466 / 0.64 | 0 / 0 | 191 / 0.26 |
| | PT | 343 | 104 / 0.3 | 0 / 0 | 0 / 0 | 257 / 0.75 | 0 / 0 | 33 / 0.1 | 15 / 0.04 | 0 / 0 | 0 / 0 | 79 / 0.23 | 0 / 0 | 26 / 0.08 |
| | RE | 376 | 31 / 0.08 | 0 / 0 | 0 / 0 | 253 / 0.67 | 0 / 0 | 171 / 0.45 | 8 / 0.02 | 15 / 0.04 | 0 / 0 | 194 / 0.52 | 0 / 0 | 221 / 0.59 |
| | RT | 2556 | 202 / 0.08 | 0 / 0 | 1 / 0 | 630 / 0.25 | 0 / 0 | 209 / 0.08 | 595 / 0.23 | 0 / 0 | 11 / 0 | 413 / 0.16 | 66 / 0.03 | 65 / 0.03 |
| | CM | 159 | 33 / 0.21 | 0 / 0 | 0 / 0 | 113 / 0.71 | 0 / 0 | 22 / 0.14 | 0 / 0 | 7 / 0.04 | 2 / 0.01 | 14 / 0.09 | 0 / 0 | 15 / 0.09 |
| | PO | 1242 | 33 / 0.03 | 39 / 0.03 | 0 / 0 | 94 / 0.08 | 15 / 0.01 | 254 / 0.2 | 118 / 0.1 | 0 / 0 | 0 / 0 | 353 / 0.28 | 3 / 0 | 90 / 0.07 |
| | PF | 606 | 6 / 0.01 | 0 / 0 | 0 / 0 | 62 / 0.1 | 0 / 0 | 89 / 0.15 | 124 / 0.2 | 0 / 0 | 0 / 0 | 246 / 0.41 | 0 / 0 | 83 / 0.14 |
| | MH | 212 | 0 / 0 | 0 / 0 | 0 / 0 | 64 / 0.3 | 0 / 0 | 47 / 0.22 | 74 / 0.35 | 0 / 0 | 0 / 0 | 142 / 0.67 | 0 / 0 | 59 / 0.28 |
| | HY | 1075 | 48 / 0.04 | 238 / 0.22 | 184 / 0.17 | 125 / 0.12 | 40 / 0.04 | 101 / 0.09 | 5 / 0 | 56 / 0.05 | 52 / 0.05 | 173 / 0.16 | 15 / 0.01 | 188 / 0.17 |
| | MPO | 1334 | 120 / 0.09 | 81 / 0.06 | 6 / 0 | 556 / 0.42 | 7 / 0.01 | 688 / 0.52 | 54 / 0.04 | 8 / 0.01 | 40 / 0.03 | 676 / 0.51 | 26 / 0.02 | 370 / 0.28 |
| | AVPV | 82 | 5 / 0.06 | 0 / 0 | 0 / 0 | 56 / 0.68 | 2 / 0.02 | 44 / 0.54 | 0 / 0 | 0 / 0 | 0 / 0 | 32 / 0.39 | 0 / 0 | 20 / 0.24 |
| HYP | PVH | 196 | 15 / 0.08 | 0 / 0 | 0 / 0 | 103 / 0.53 | 0 / 0 | 59 / 0.3 | 0 / 0 | 19 / 0.1 | 0 / 0 | 106 / 0.54 | 0 / 0 | 143 / 0.73 |
| | SCH | 115 | 9 / 0.08 | 15 / 0.13 | 4 / 0.03 | 14 / 0.12 | 0 / 0 | 34 / 0.3 | 20 / 0.17 | 0 / 0 | 13 / 0.11 | 41 / 0.36 | 2 / 0.02 | 0 / 0 |
| | AHN | 517 | 1 / 0 | 15 / 0.03 | 0 / 0 | 48 / 0.09 | 0 / 0 | 80 / 0.15 | 0 / 0 | 31 / 0.06 | 0 / 0 | 66 / 0.13 | 0 / 0 | 140 / 0.27 |
| | SBPV | 199 | 0 / 0 | 2 / 0.01 | 2 / 0.01 | 44 / 0.22 | 0 / 0 | 20 / 0.1 | 1 / 0.01 | 7 / 0.04 | 0 / 0 | 52 / 0.26 | 0 / 0 | 32 / 0.16 |
| | LHA | 2478 | 17 / 0.01 | 60 / 0.02 | 12 / 0 | 812 / 0.33 | 1 / 0 | 818 / 0.33 | 28 / 0.01 | 79 / 0.03 | 4 / 0 | 466 / 0.19 | 3 / 0 | 526 / 0.21 |

| | | | | | | | | | | | | | | |
|---|---|---|---|---|---|---|---|---|---|---|---|---|---|---|
| | ZI | 1764 | 22 / 0.01 | 23 / 0.01 | 0 / 0 | 899 / 0.51 | 0 / 0 | 798 / 0.45 | 164 / 0.09 | 39 / 0.02 | 0 / 0 | 547 / 0.31 | 1 / 0 | 418 / 0.24 |
| | ARH | 152 | 0 / 0 | 108 / 0.71 | 98 / 0.64 | 0 / 0 | 0 / 0 | 0 / 0 | 0 / 0 | 1 / 0.01 | 0 / 0 | 11 / 0.07 | 0 / 0 | 41 / 0.27 |
| | VMH | 368 | 0 / 0 | 149 / 0.4 | 20 / 0.05 | 2 / 0.01 | 0 / 0 | 25 / 0.07 | 7 / 0.02 | 0 / 0 | 0 / 0 | 69 / 0.19 | 0 / 0 | 55 / 0.15 |
| | DMH | 390 | 0 / 0 | 17 / 0.04 | 0 / 0 | 90 / 0.23 | 0 / 0 | 107 / 0.27 | 2 / 0.01 | 15 / 0.04 | 0 / 0 | 172 / 0.44 | 0 / 0 | 196 / 0.5 |
| | PVp | 243 | 0 / 0 | 115 / 0.47 | 77 / 0.32 | 0 / 0 | 4 / 0.02 | 7 / 0.03 | 20 / 0.08 | 0 / 0 | 2 / 0.01 | 48 / 0.2 | 0 / 0 | 8 / 0.03 |
| | PH | 570 | 0 / 0 | 2 / 0 | 0 / 0 | 407 / 0.71 | 0 / 0 | 474 / 0.83 | 0 / 0 | 16 / 0.03 | 0 / 0 | 309 / 0.54 | 0 / 0 | 336 / 0.59 |
| | STN | 271 | 0 / 0 | 0 / 0 | 0 / 0 | 162 / 0.6 | 0 / 0 | 142 / 0.52 | 10 / 0.04 | 3 / 0.01 | 0 / 0 | 140 / 0.52 | 1 / 0 | 131 / 0.48 |
| | MBO | 892 | 0 / 0 | 119 / 0.13 | 69 / 0.08 | 82 / 0.09 | 20 / 0.02 | 199 / 0.22 | 21 / 0.02 | 31 / 0.03 | 2 / 0 | 176 / 0.2 | 8 / 0.01 | 157 / 0.18 |
| MB | MB | 19996 | 211 / 0.01 | 956 / 0.05 | 184 / 0.01 | 2114 / 0.11 | 173 / 0.01 | 3752 / 0.19 | 774 / 0.04 | 2030 / 0.1 | 435 / 0.02 | 3630 / 0.18 | 161 / 0.01 | 4887 / 0.24 |
| | VTA | 764 | 0 / 0 | 13 / 0.02 | 3 / 0 | 197 / 0.26 | 2 / 0 | 453 / 0.59 | 11 / 0.01 | 66 / 0.09 | 3 / 0 | 185 / 0.24 | 1 / 0 | 314 / 0.41 |
| | RN | 344 | 0 / 0 | 4 / 0.01 | 0 / 0 | 49 / 0.14 | 0 / 0 | 111 / 0.32 | 0 / 0 | 51 / 0.15 | 0 / 0 | 8 / 0.02 | 0 / 0 | 46 / 0.13 |
| | SNc | 724 | 0 / 0 | 0 / 0 | 0 / 0 | 109 / 0.15 | 0 / 0 | 289 / 0.4 | 38 / 0.05 | 87 / 0.12 | 0 / 0 | 288 / 0.4 | 0 / 0 | 368 / 0.51 |
| | SNr | 1165 | 0 / 0 | 12 / 0.01 | 6 / 0.01 | 80 / 0.07 | 5 / 0 | 208 / 0.18 | 71 / 0.06 | 42 / 0.04 | 1 / 0 | 291 / 0.25 | 0 / 0 | 307 / 0.26 |
| | IPN | 281 | 0 / 0 | 41 / 0.15 | 29 / 0.1 | 89 / 0.32 | 7 / 0.02 | 130 / 0.46 | 1 / 0 | 45 / 0.16 | 14 / 0.05 | 35 / 0.12 | 0 / 0 | 52 / 0.19 |
| | CLI | 132 | 0 / 0 | 13 / 0.1 | 0 / 0 | 35 / 0.27 | 0 / 0 | 63 / 0.48 | 0 / 0 | 5 / 0.04 | 0 / 0 | 0 / 0 | 0 / 0 | 34 / 0.26 |
| | CS | 597 | 4 / 0.01 | 6 / 0.01 | 4 / 0.01 | 111 / 0.19 | 2 / 0 | 118 / 0.2 | 0 / 0 | 24 / 0.04 | 4 / 0.01 | 17 / 0.03 | 0 / 0 | 152 / 0.25 |
| | DR | 175 | 0 / 0 | 12 / 0.07 | 7 / 0.04 | 28 / 0.16 | 4 / 0.02 | 46 / 0.26 | 0 / 0 | 22 / 0.13 | 0 / 0 | 25 / 0.14 | 0 / 0 | 81 / 0.46 |
| | PAG | 4263 | 67 / 0.02 | 74 / 0.02 | 54 / 0.01 | 1027 / 0.24 | 55 / 0.01 | 955 / 0.22 | 170 / 0.04 | 253 / 0.06 | 8 / 0 | 1536 / 0.36 | 2 / 0 | 2208 / 0.52 |
| HB | P | 5550 | 106 / 0.02 | 421 / 0.08 | 1468 / 0.26 | 24 / 0 | 1140 / 0.21 | 72 / 0.01 | 21 / 0 | 794 / 0.14 | 1357 / 0.24 | 40 / 0.01 | 404 / 0.07 | 217 / 0.04 |
| | PG | 712 | 0 / 0 | 177 / 0.25 | 312 / 0.44 | 5 / 0.01 | 145 / 0.2 | 13 / 0.02 | 40 / 0.06 | 14 / 0.02 | 28 / 0.04 | 2 / 0 | 72 / 0.1 | 0 / 0 |
| | PCG | 803 | 16 / 0.02 | 121 / 0.15 | 216 / 0.27 | 13 / 0.02 | 89 / 0.11 | 22 / 0.03 | 0 / 0 | 298 / 0.37 | 26 / 0.03 | 17 / 0.02 | 0 / 0 | 140 / 0.17 |
| | PRN | 5542 | 25 / 0 | 266 / 0.05 | 799 / 0.14 | 273 / 0.05 | 490 / 0.09 | 453 / 0.08 | 0 / 0 | 650 / 0.12 | 117 / 0.02 | 42 / 0.01 | 10 / 0 | 593 / 0.11 |
| | PB | 1151 | 0 / 0 | 156 / 0.14 | 188 / 0.16 | 64 / 0.06 | 101 / 0.09 | 171 / 0.15 | 0 / 0 | 377 / 0.33 | 154 / 0.13 | 37 / 0.03 | 14 / 0.01 | 224 / 0.19 |
| | LC | 96 | 0 / 0 | 3 / 0.03 | 7 / 0.07 | 0 / 0 | 4 / 0.04 | 1 / 0.01 | 0 / 0 | 71 / 0.74 | 13 / 0.14 | 0 / 0 | 0 / 0 | 46 / 0.48 |
| | MY | 28535 | 1222 / 0.04 | 1666 / 0.06 | 3250 / 0.11 | 682 / 0.02 | 2968 / 0.1 | 320 / 0.01 | 190 / 0.01 | 4020 / 0.14 | 5745 / 0.2 | 286 / 0.01 | 1740 / 0.06 | 513 / 0.02 |
| CB | CB | 58909 | 3408 / 0.06 | 2083 / 0.04 | 7951 / 0.13 | 741 / 0.01 | 10822 / 0.18 | 796 / 0.01 | 1106 / 0.02 | 7326 / 0.12 | 12109 / 0.21 | 2664 / 0.05 | 6477 / 0.11 | 2002 / 0.03 |
| | DEC | 960 | 132 / 0.14 | 22 / 0.02 | 49 / 0.05 | 0 / 0 | 215 / 0.22 | 0 / 0 | 134 / 0.14 | 317 / 0.33 | 75 / 0.08 | 206 / 0.21 | 0 / 0 | 54 / 0.06 |
| | FOTU | 226 | 29 / 0.13 | 0 / 0 | 6 / 0.03 | 1 / 0 | 62 / 0.27 | 0 / 0 | 19 / 0.08 | 100 / 0.44 | 34 / 0.15 | 34 / 0.15 | 0 / 0 | 5 / 0.02 |
| | PYR | 609 | 22 / 0.04 | 0 / 0 | 29 / 0.05 | 0 / 0 | 153 / 0.25 | 0 / 0 | 0 / 0 | 30 / 0.05 | 349 / 0.57 | 0 / 0 | 281 / 0.46 | 0 / 0 |
| | NOD | 556 | 17 / 0.03 | 0 / 0 | 37 / 0.07 | 0 / 0 | 25 / 0.04 | 0 / 0 | 0 / 0 | 25 / 0.04 | 140 / 0.25 | 0 / 0 | 20 / 0.04 | 0 / 0 |
| | UVU | 1888 | 126 / 0.07 | 1 / 0 | 93 / 0.05 | 0 / 0 | 394 / 0.21 | 0 / 0 | 0 / 0 | 272 / 0.14 | 948 / 0.5 | 0 / 0 | 656 / 0.35 | 0 / 0 |
| | SIM | 236 | 38 / 0.16 | 0 / 0 | 6 / 0.03 | 0 / 0 | 56 / 0.24 | 0 / 0 | 18 / 0.08 | 56 / 0.24 | 3 / 0.01 | 41 / 0.17 | 2 / 0.01 | 20 / 0.08 |

**Table S7.** Segment-wise fractional difference volumes (FDV) between groups at each condition, as in **Fig. 6** of the main text.

Columns for "Domain", "Segment" and "Total Segment Voxels" are exactly as in **Table S6A-B**. Values in the right columns represent fractional difference volumes for two-sample t-tests at each condition, which compared post-Mn(II) images of Standard with those of ELA after adjusting for pre-Mn(II) differences between groups. Adjustments were made as described in Main Text Methods ***MR Image Analysis***, and ***SI Appendix*** Extended Methods 5.3, where the difference image between the ELA and Std group averages at baseline was subtracted from all ELA pre- and post-Mn(II) images prior to SPM t-tests and segmentation.

| Domain | Segment | Total Segment Voxels | Significant Voxels/FDV by Condition | | | | | |
|---|---|---|---|---|---|---|---|---|
| | | | Std > ELA | | | ELA > Standard | | |
| | | | HC | TMT | D9 | HC | TMT | D9 |
| OLF | MOBgl | 6904 | 0.015 | 0.026 | 0.025 | 0.364 | 0.124 | 0.043 |
| | MOBgr | 4577 | 0.001 | 0.002 | 0.003 | 0.213 | 0.068 | 0.076 |
| | MOBipl | 4912 | 0.002 | 0.003 | 0.003 | 0.461 | 0.157 | 0.110 |
| | MOBmi | 1064 | 0.001 | 0.004 | 0.003 | 0.526 | 0.227 | 0.064 |
| | MOBopl | 1234 | 0.008 | 0.007 | 0.009 | 0.340 | 0.119 | 0.053 |
| | AOB | 599 | 0.000 | 0.005 | 0.082 | 0.144 | 0.132 | 0.008 |
| | AON | 3084 | 0.000 | 0.032 | 0.002 | 0.252 | 0.062 | 0.036 |
| | TT | 1097 | 0.001 | 0.095 | 0.000 | 0.154 | 0.013 | 0.105 |
| | EPd | 910 | 0.000 | 0.170 | 0.023 | 0.171 | 0.035 | 0.036 |
| CTX | CTX | 48931 | 0.105 | 0.106 | 0.104 | 0.046 | 0.028 | 0.054 |
| | ORB | 2379 | 0.014 | 0.066 | 0.027 | 0.076 | 0.029 | 0.061 |
| | PL | 1906 | 0.026 | 0.079 | 0.070 | 0.037 | 0.081 | 0.034 |
| | ILA | 1371 | 0.030 | 0.083 | 0.001 | 0.105 | 0.056 | 0.148 |
| | DP | 454 | 0.000 | 0.165 | 0.000 | 0.192 | 0.013 | 0.187 |
| | ACA | 5311 | 0.031 | 0.079 | 0.031 | 0.143 | 0.099 | 0.047 |
| | MO | 16452 | 0.075 | 0.086 | 0.046 | 0.033 | 0.057 | 0.027 |
| | SS | 30181 | 0.110 | 0.110 | 0.058 | 0.019 | 0.047 | 0.061 |
| | PIR | 9948 | 0.006 | 0.052 | 0.038 | 0.205 | 0.065 | 0.039 |
| | RSP | 6191 | 0.126 | 0.071 | 0.047 | 0.010 | 0.017 | 0.025 |
| | PTL | 2826 | 0.107 | 0.031 | 0.040 | 0.021 | 0.077 | 0.051 |
| HIP | HPF | 10162 | 0.090 | 0.136 | 0.031 | 0.051 | 0.021 | 0.138 |
| | DG | 2501 | 0.075 | 0.112 | 0.014 | 0.052 | 0.029 | 0.079 |
| | CA1-CA3 | 12826 | 0.032 | 0.062 | 0.019 | 0.098 | 0.077 | 0.199 |
| AMY | AAA | 667 | 0.000 | 0.015 | 0.046 | 0.234 | 0.063 | 0.045 |
| | MEA | 1092 | 0.001 | 0.016 | 0.017 | 0.109 | 0.073 | 0.055 |
| | CEA | 1578 | 0.012 | 0.024 | 0.009 | 0.100 | 0.235 | 0.104 |
| | LA | 1261 | 0.042 | 0.093 | 0.021 | 0.019 | 0.019 | 0.059 |

| | | | | | | | | |
|---|---|---|---|---|---|---|---|---|
| | BLA | 3253 | 0.009 | 0.031 | 0.040 | 0.147 | 0.053 | 0.065 |
| | PA | 463 | 0.017 | 0.004 | 0.000 | 0.091 | 0.028 | 0.307 |
| | COA | 2626 | 0.011 | 0.034 | 0.008 | 0.252 | 0.036 | 0.157 |
| STR/PAL | CP | 22436 | 0.020 | 0.121 | 0.063 | 0.194 | 0.026 | 0.063 |
| | ACB | 5108 | 0.011 | 0.057 | 0.011 | 0.302 | 0.090 | 0.040 |
| | FS | 570 | 0.000 | 0.021 | 0.002 | 0.302 | 0.256 | 0.025 |
| | OT | 2551 | 0.011 | 0.130 | 0.024 | 0.124 | 0.034 | 0.081 |
| | LSc | 338 | 0.009 | 0.104 | 0.000 | 0.444 | 0.033 | 0.361 |
| | LSr | 1952 | 0.000 | 0.029 | 0.002 | 0.587 | 0.077 | 0.248 |
| | GPe | 1678 | 0.000 | 0.002 | 0.004 | 0.289 | 0.371 | 0.029 |
| | SI | 2927 | 0.000 | 0.043 | 0.004 | 0.415 | 0.177 | 0.107 |
| | MS | 220 | 0.000 | 0.000 | 0.000 | 0.886 | 0.282 | 0.286 |
| | NDB | 661 | 0.000 | 0.008 | 0.064 | 0.413 | 0.038 | 0.018 |
| | BST | 1800 | 0.000 | 0.002 | 0.000 | 0.481 | 0.159 | 0.267 |
| THA | VAL | 413 | 0.017 | 0.022 | 0.046 | 0.027 | 0.034 | 0.048 |
| | VM | 936 | 0.071 | 0.049 | 0.200 | 0.007 | 0.089 | 0.005 |
| | VPM | 1020 | 0.048 | 0.102 | 0.133 | 0.120 | 0.006 | 0.023 |
| | VPL | 1298 | 0.040 | 0.042 | 0.125 | 0.111 | 0.042 | 0.011 |
| | SPA | 202 | 0.040 | 0.203 | 0.025 | 0.000 | 0.000 | 0.000 |
| | MG | 1195 | 0.040 | 0.054 | 0.053 | 0.088 | 0.008 | 0.060 |
| | LGd | 53 | 0.038 | 0.075 | 0.075 | 0.245 | 0.000 | 0.075 |
| | LP | 1353 | 0.068 | 0.085 | 0.056 | 0.035 | 0.000 | 0.038 |
| | AV | 194 | 0.000 | 0.000 | 0.000 | 0.227 | 0.000 | 0.144 |
| | IAM | 507 | 0.024 | 0.024 | 0.000 | 0.079 | 0.032 | 0.079 |
| | IMD | 65 | 0.215 | 0.169 | 0.231 | 0.000 | 0.000 | 0.000 |
| | PVT | 1349 | 0.030 | 0.069 | 0.000 | 0.104 | 0.050 | 0.152 |
| | MD | 725 | 0.254 | 0.312 | 0.051 | 0.030 | 0.000 | 0.023 |
| | PT | 343 | 0.087 | 0.035 | 0.082 | 0.017 | 0.070 | 0.012 |
| | RE | 376 | 0.005 | 0.021 | 0.005 | 0.059 | 0.234 | 0.064 |
| | RT | 2556 | 0.006 | 0.013 | 0.047 | 0.242 | 0.105 | 0.076 |
| | CM | 159 | 0.013 | 0.000 | 0.126 | 0.019 | 0.126 | 0.006 |
| | PO | 1242 | 0.060 | 0.197 | 0.132 | 0.039 | 0.000 | 0.032 |
| | PF | 606 | 0.040 | 0.243 | 0.059 | 0.030 | 0.013 | 0.046 |
| | MH | 212 | 0.000 | 0.014 | 0.000 | 0.302 | 0.071 | 0.297 |
| HYP | HY | 1075 | 0.007 | 0.056 | 0.002 | 0.218 | 0.138 | 0.397 |
| | MPO | 1334 | 0.000 | 0.000 | 0.000 | 0.545 | 0.360 | 0.394 |
| | AVPV | 82 | 0.000 | 0.000 | 0.000 | 0.451 | 0.671 | 0.293 |
| | PVH | 196 | 0.041 | 0.000 | 0.000 | 0.082 | 0.163 | 0.250 |
| | SCH | 115 | 0.000 | 0.000 | 0.000 | 0.435 | 0.357 | 0.235 |
| | AHN | 517 | 0.000 | 0.000 | 0.000 | 0.191 | 0.029 | 0.257 |
| | SBPV | 199 | 0.015 | 0.025 | 0.000 | 0.035 | 0.015 | 0.176 |

| | | | | | | | | |
|---|---|---|---|---|---|---|---|---|
| | LHA | 2478 | 0.000 | 0.002 | 0.003 | 0.372 | 0.270 | 0.228 |
| | ZI | 1764 | 0.005 | 0.052 | 0.043 | 0.166 | 0.132 | 0.077 |
| | ARH | 152 | 0.000 | 0.355 | 0.000 | 0.053 | 0.000 | 0.553 |
| | VMH | 368 | 0.000 | 0.003 | 0.000 | 0.576 | 0.052 | 0.856 |
| | DMH | 390 | 0.003 | 0.000 | 0.000 | 0.505 | 0.479 | 0.800 |
| | PVp | 243 | 0.025 | 0.173 | 0.012 | 0.111 | 0.000 | 0.214 |
| | PH | 570 | 0.000 | 0.000 | 0.002 | 0.165 | 0.207 | 0.174 |
| | STN | 271 | 0.000 | 0.004 | 0.000 | 0.196 | 0.107 | 0.066 |
| | MBO | 892 | 0.018 | 0.071 | 0.024 | 0.111 | 0.053 | 0.178 |
| MB | MB | 19996 | 0.060 | 0.068 | 0.050 | 0.028 | 0.048 | 0.061 |
| | VTA | 764 | 0.003 | 0.009 | 0.020 | 0.076 | 0.084 | 0.030 |
| | RN | 344 | 0.035 | 0.006 | 0.163 | 0.000 | 0.012 | 0.003 |
| | SNc | 724 | 0.061 | 0.041 | 0.012 | 0.062 | 0.047 | 0.151 |
| | SNr | 1165 | 0.058 | 0.121 | 0.036 | 0.040 | 0.003 | 0.055 |
| | IPN | 281 | 0.000 | 0.011 | 0.000 | 0.310 | 0.274 | 0.064 |
| | CLI | 132 | 0.038 | 0.182 | 0.152 | 0.061 | 0.076 | 0.023 |
| | CS | 597 | 0.002 | 0.000 | 0.000 | 0.191 | 0.337 | 0.221 |
| | DR | 175 | 0.126 | 0.011 | 0.000 | 0.000 | 0.000 | 0.000 |
| | PAG | 4263 | 0.128 | 0.097 | 0.030 | 0.020 | 0.045 | 0.079 |
| HB | P | 5550 | 0.118 | 0.028 | 0.012 | 0.043 | 0.112 | 0.063 |
| | PG | 712 | 0.003 | 0.042 | 0.004 | 0.219 | 0.032 | 0.091 |
| | PCG | 803 | 0.037 | 0.000 | 0.000 | 0.004 | 0.127 | 0.278 |
| | PRN | 5542 | 0.192 | 0.073 | 0.015 | 0.033 | 0.102 | 0.040 |
| | PB | 1151 | 0.093 | 0.011 | 0.006 | 0.019 | 0.092 | 0.047 |
| | LC | 96 | 0.135 | 0.000 | 0.000 | 0.000 | 0.333 | 0.344 |
| | MY | 28535 | 0.057 | 0.016 | 0.046 | 0.067 | 0.200 | 0.069 |
| CB | CB | 58909 | 0.099 | 0.052 | 0.050 | 0.037 | 0.150 | 0.050 |
| | DEC | 960 | 0.286 | 0.165 | 0.031 | 0.000 | 0.050 | 0.036 |
| | FOTU | 226 | 0.642 | 0.124 | 0.000 | 0.000 | 0.004 | 0.000 |
| | PYR | 609 | 0.010 | 0.000 | 0.049 | 0.043 | 0.302 | 0.016 |
| | NOD | 556 | 0.000 | 0.000 | 0.009 | 0.297 | 0.631 | 0.112 |
| | UVU | 1888 | 0.010 | 0.001 | 0.017 | 0.064 | 0.447 | 0.056 |
| | SIM | 236 | 0.275 | 0.148 | 0.000 | 0.000 | 0.059 | 0.055 |

**Legend for Movie S1**

**Movie S1.** Projections of Statistical Maps of Between-Group Differences on the *InVivo* Atlas. Movie shows coronal slices of 3D statistical maps projected onto the *InVivo* Atlas, stepping at 100 μm from anterior to posterior. Segment boundaries are represented by colored lines. These maps were used to obtain the FDV values shown as column graphs in **Fig. 6**. *Upper right*: Range of t-values corresponding to the color gradient of statistical maps. *Lower left*: Range of voxel-wise Cohen's D effect sizes: minimum - maximum. Visualization in FSLeyes was used to step through brain images while recording in Screen Recorder Pro, and Microsoft Clipchamp generated the MP4 movie.

**SI References**


1. Uselman TW, Barto DR, Jacobs RE, & Bearer EL (2020) Evolution of brain-wide activity in the awake behaving mouse after acute fear by longitudinal manganese-enhanced MRI. *Neuroimage* 222:116975.
2. Uselman TW, Medina CS, Gray HB, Jacobs RE, & Bearer EL (2022) Longitudinal manganese-enhanced magnetic resonance imaging of neural projections and activity.e4675.
3. Garcia-Arocena D (2015) More researchers are using B6J mice than ever before. (The Jackson Laboratory).
4. Care IoLARCo & Animals UoL (1986) *Guide for the care and use of laboratory animals* (US Department of Health and Human Services, Public Health Service, National …).
5. Laboratory TJ (2024) Mouse Room Conditions.
6. Gozalo AS & Elkins WR (2023) A Review of the Effects of Some Extrinsic Factors on Mice Used in Research %J Comparative Medicine. 73(6):413-431.
7. Ben-Shachar M, Lüdecke D, & Makowski D (2020) effectsize: Estimation of Effect Size Indices and Standardized Parameters. *Journal of Open Source Software* 5(56).
8. Frahm J, Haase A, & Matthaei D (1986) Rapid NMR imaging of dynamic processes using the FLASII technique. 3(2):321-327.
9. Haase A, Frahm J, Matthaei D, Hanicke W, & Merboldt KD (1986) FLASH imaging. Rapid NMR imaging using low flip-angle pulses. *Journal of Magnetic Resonance (1969)* 67(2):258-266.
10. Wickham H (2016) ggplot2: Elegant Graphics for Data Analysis.
11. Fendt M, Endres T, Lowry CA, Apfelbach R, & McGregor IS (2005) TMT-induced autonomic and behavioral changes and the neural basis of its processing. *Neuroscience and biobehavioral reviews* 29(8):1145-1156.
12. Delora A*, et al.* (2016) A simple rapid process for semi-automated brain extraction from magnetic resonance images of the whole mouse head. *J Neurosci Methods* 257:185-193.
13. Medina CS, Manifold-Wheeler B, Gonzales A, & Bearer EL (2017) Automated Computational Processing of 3-D MR Images of Mouse Brain for Phenotyping of Living Animals. *Curr Protoc Mol Biol* 119:29A 25 21-29A 25 38.
14. Ashburner J*, et al.* (2014) SPM12 manual.
15. Schindelin J*, et al.* (2012) Fiji: an open-source platform for biological-image analysis. *Nature Methods* 9(7):676-682.
16. Pinheiro J, Bates D, & Team RC (2022) nlme: Linear and Nonlinear Mixed Effects Models.
17. Jaccard P (1912) THE DISTRIBUTION OF THE FLORA IN THE ALPINE ZONE. 11(2):37-50.

18. Jenkinson M, Beckmann CF, Behrens TE, Woolrich MW, & Smith SMJN (2012) Fsl. 62(2):782-790.
19. Studholme C, Hawkes D, & Hill D (1998) *Normalized entropy measure for multimodality image alignment* (SPIE).
20. Vinh NX, Epps J, & Bailey J (2009) Information theoretic measures for clusterings comparison: is a correction for chance necessary? in *Proceedings of the 26th Annual International Conference on Machine Learning* (Association for Computing Machinery, Montreal, Quebec, Canada), pp 1073–1080.
21. Bearer EL*, et al.* (2007) Role of neuronal activity and kinesin on tract tracing by manganese-enhanced MRI (MEMRI). *Neuroimage* 37 Suppl 1:S37-46.
22. Bearer EL, Zhang X, & Jacobs RE (2007) Live imaging of neuronal connections by magnetic resonance: Robust transport in the hippocampal–septal memory circuit in a mouse model of Down syndrome. *NeuroImage* 37(1):230-242.
23. Gallagher JJ*, et al.* (2013) Altered reward circuitry in the norepinephrine transporter knockout mouse. *PLoS One* 8(3):e57597.
24. Medina CS*, et al.* (2017) Hippocampal to basal forebrain transport of Mn(2+) is impaired by deletion of KLC1, a subunit of the conventional kinesin microtubule-based motor. *Neuroimage* 145(Pt A):44-57.
25. Medina CS*, et al.* (2019) Decoupling the Effects of the Amyloid Precursor Protein From Amyloid-beta Plaques on Axonal Transport Dynamics in the Living Brain. *Front Cell Neurosci* 13:501.
26. Zhang X*, et al.* (2010) Altered neurocircuitry in the dopamine transporter knockout mouse brain. *PLoS One* 5(7):e11506.
27. Ashburner J*, et al.* (2012) SPM8 manual.
28. Flandin G & Friston K (2008) Statistical parametric mapping (SPM). *Scholarpedia* 3(4):6232.
29. Wang Q*, et al.* (2020) The Allen Mouse Brain Common Coordinate Framework: A 3D Reference Atlas. *Cell* 181(4):936-953.e920.
30. Bearer EL, Medina CS, Uselman TW, & Jacobs RE (2023) Harnessing axonal transport to map reward circuitry: Differing brain-wide projections from medial prefrontal cortical domains. *Frontiers in Cell and Developmental Biology* 11.
31. Tyszka JM, Readhead C, Bearer EL, Pautler RG, & Jacobs RE (2006) Statistical diffusion tensor histology reveals regional dysmyelination effects in the shiverer mouse mutant. *NeuroImage* 29(4):1058-1065.
32. Seamster PE*, et al.* (2012) Quantitative measurements and modeling of cargo–motor interactions during fast transport in the living axon. *Physical Biology* 9(5):055005.
